# Interplay of adherens junctions and matrix proteolysis determines the invasive pattern and growth of squamous cell carcinoma

**DOI:** 10.1101/2021.11.24.469674

**Authors:** Takuya Kato, Robert P. Jenkins, Stefanie Derzsi, Melda Tozluoglu, Antonio Rullan, Steven Hooper, Raphael A. G. Chaleil, Xiao Fu, Holly Joyce, Selvam Thavaraj, Paul A. Bates, Erik Sahai

**Affiliations:** Tumour Cell Biology Laboratory, The Francis Crick Institute, 1 Midland Road, London, NW1 1AT, UK; Department of Pathology, Kitasato University, 1-15-1 Kitasato, Sagamihara, Japan; Hoffman La-Roche, 4070 Basel, Switzerland; Biomolecular Modelling Laboratory, The Francis Crick Institute, 1 Midland Road, London, NW1 1AT, UK; Institute of Cancer Research, 237 Fulham Road, London, SW3 6JB, UK; Centre of Clinical, Oral and Translational Science, King’s College London, Great Maze Pond, London, SE1 9RT, UK

## Abstract

Cancers, such as squamous cell carcinoma, frequently invade as multicellular units. However, these invading units can be organized in a variety of ways, ranging from thin discontinuous strands to thick ‘pushing’ collectives. Here we employ an integrated experimental and computational approach to identify the factors that determine the mode of collective cancer cell invasion. We find that matrix proteolysis is linked to the formation of wide strands, but has little effect on the maximum extent of invasion. Cell-cell junctions also favour wide strands, but our analysis also reveals a requirement for cell-cell junctions for efficient invasion in response to uniform directional cues. Unexpectedly, the ability to generate wide invasive strands is coupled to the ability to grow effectively when surrounded by ECM in 3D assays. Combinatorial perturbation of both matrix proteolysis and cell-cell adhesion demonstrates that the most aggressive cancer behaviour, both in terms of invasion and growth, is achieved at high levels of cell-cell adhesion and high levels of proteolysis. Contrary to expectation, cells with canonical mesenchymal traits – no cell-cell junctions and high proteolysis – exhibit reduced growth and lymph node metastasis. Thus, we conclude that the ability of squamous cell carcinoma cells to invade effectively is also linked to their ability to generate space for proliferation in confined contexts. These data provide an explanation for the apparent advantage of retaining cell-cell junctions in SCC.

## Introduction

Tumours exhibit a variety of histological patterns that inform pathological diagnosis and that are frequently linked to prognosis (Mishra et al., 2014). This link with outcome suggests that the mechanisms specifying histological pattern are related to tumour malignancy. This may be due to some coupling between how cancer cells invade and their ability to proliferate. Epithelial cancer cells, including squamous cell carcinoma, frequently invade in collective units (Friedl and Gilmour, 2009; Khalil et al., 2017; Wang et al., 2016). The importance of collective invasion is underscored by several recent studies showing that collective seeding of metastases is more efficient than single cell seeding (Cheung et al., 2016; Fischer et al., 2015; Khalil et al., 2017). Despite the prevalence and importance of collective patterns of cancer cell invasion, it remains less well understood than single cell forms of invasion. Collectively invading cancer strands can be organized in a variety of different ways, from single file strands that characterize invasive lobular breast cancer and diffuse gastric cancer to broad cohesive units found in basal cell carcinoma (Boelens et al., 2016; Carneiro et al., 2004; Friedl et al., 2012; Pandya et al., 2017). Histological analysis indicates that even within a single disease type there is considerable heterogeneity in the pattern of invasion; for example, both broad ‘pushing’ and strand-like infiltrative invasion can be observed in squamous cell carcinoma (Dissanayaka et al., 2012). In this study, we set out to explore the key parameters that determine the pattern of collective invasion using a combination of computational and experimental approaches.

Several parameters might be expected to modulate tumour histology and, more specifically, collective cancer cell invasion. The ability of cancer cells to adhere to each other through cadherin mediated junctions is linked to their organization into tightly packed clusters. E-cadherin/CDH1 and, to a lesser extent, P-cadherin/CDH3 are the predominant cadherins in mucosal squamous cell carcinomas (SCC or muSCC specifically for mucosal SCC) (Nieman et al., 1999) that typically do not undergo a clear epithelial to mesenchymal transition (EMT). These cadherins are coupled to the actin cytoskeleton via a complex containing α-catenin and β-catenin (Nelson et al., 2013). Cell adhesion to the extracellular matrix (ECM) is also critical for cell migration and invasion in many contexts (Cooper and Giancotti, 2019; Hamidi and Ivaska, 2018). This is primarily mediated by integrin receptors (Hamidi and Ivaska, 2018; Janes and Watt, 2006), with *ITGB1* particularly highly expressed in SCC (Janes and Watt, 2006). The extracellular matrix presents a barrier to migration if the gaps between fibres are smaller than 3-5μm (Wolf et al., 2013, 2009). The dermal ECM underlying SCC lesions is predominantly composed of type I collagen (Watt and Fujiwara, 2011) and numerous studies have demonstrated that MMP14/MT-1MMP is the critical protease for degrading this type of matrix (Castro-Castro et al., 2016; Gifford and Itoh, 2019). The ECM can also be physically moved by forces generated by the contractile cytoskeleton (Mohammadi and Sahai, 2018; Wolf et al., 2013). In many cases, stromal cells are the major source of both matrix proteolytic and force-mediated matrix remodelling in tumours (Conklin and Keely, 2012; Kalluri and Zeisberg, 2006). Cancer-associated fibroblasts (CAFs, sometimes referred to as stromal fibroblasts) can promote the invasion of SCC by providing these functions and are frequently observed leading the migration of cancer cells that retain epithelial characteristics (Gaggioli et al., 2007).

Understanding the relative contributions of the multiple parameters outlined above to cell invasion is a complex multi-dimensional problem with non-linear relationships between parameters and migratory capability. This complexity means that developing a holistic and predictive framework for collective cancer cell invasion using empirical methods only is challenging. For this reason, several studies have sought to utilize computational models. Many different types of model have been used including those based on evolutionary game theory (Basanta et al., 2008; Swierniak and Krzeslak, 2013), Bayesian networks (Katz et al., 2011), differential equations (Gerisch and Chaplain, 2008; Peng et al., 2017; Weekes et al., 2014), agent-based models including cellular automata (Alarcón et al., 2003; Bull et al., 2020; Fiore et al., 2020; Gralka and Hallatschek, 2019; Karolak et al., 2019; Norton et al., 2017; Talkenberger et al., 2017) and hybrids of the above (Anderson, 2005; Anderson et al., 2006; Osborne et al., 2010). Cellular Potts modelling (Cickovski et al., 2007; Graner and Glazier, 1992; Hallou et al., 2017; Pramanik et al., 2021; Scianna et al., 2013; Shirinifard et al., 2009; Szabó and Merks, 2013; Turner and Sherratt, 2002) is a flexible approach that uses voxels to represent different parts of cells or their environment. Changes in the properties associated with each voxel are determined at each time step using principles of probabilistic energy minimization. The behaviour of the model therefore emerges from iterative application of rules that describe the relative favourability of different events or changes. Here we combine a Potts modelling with extensive experimentation to unpick the determinants of the mode of collective cancer cell invasion and their linkage to cancer cell growth, both *in vitro* and *in vivo.*

## Results

### Diverse modes of collective invasion within individual SCC

We began by surveying the diversity of invasive pattern in mucosal squamous cell carcinoma (muSCC). Figure 1a shows considerable diversity in the nature of collective invasion. Further, it illustrates how initial invasion involves cells moving from the epithelial layer into the lamina propria (often termed epidermis and dermis, respectively, in cutaneous skin). Following the invasion into the dermis, cancer cells become surrounded on all sides by ECM. Interestingly, different patterns of invasion were observed in different regions of the same tumour (Figure 1a). These ranged from broad ‘pushing’ invasive masses of cells (box I), thinner strands of cells (box II), to single cell width strands and apparently isolated single cells (box III - although this could not be definitively determined from single H&E sections). Quantitative analysis of the number of cell neighbours provided a more objective metric of invasion type, with high neighbour numbers (typically 4 to 7) indicating broad invasion patterns and low neighbour numbers (2 or 3) indicating thin strand-like invasion, respectively (Figure 1b). Similar patterns were observed in other muSCC biopsies with different strand thickness apparent (Supp. Figure 1a-c). Analysis of neighbour number suggested that strand thickness does not fall into distinct categories, with neighbour number varying continuously between 1 and 9.

**Figure 1:**
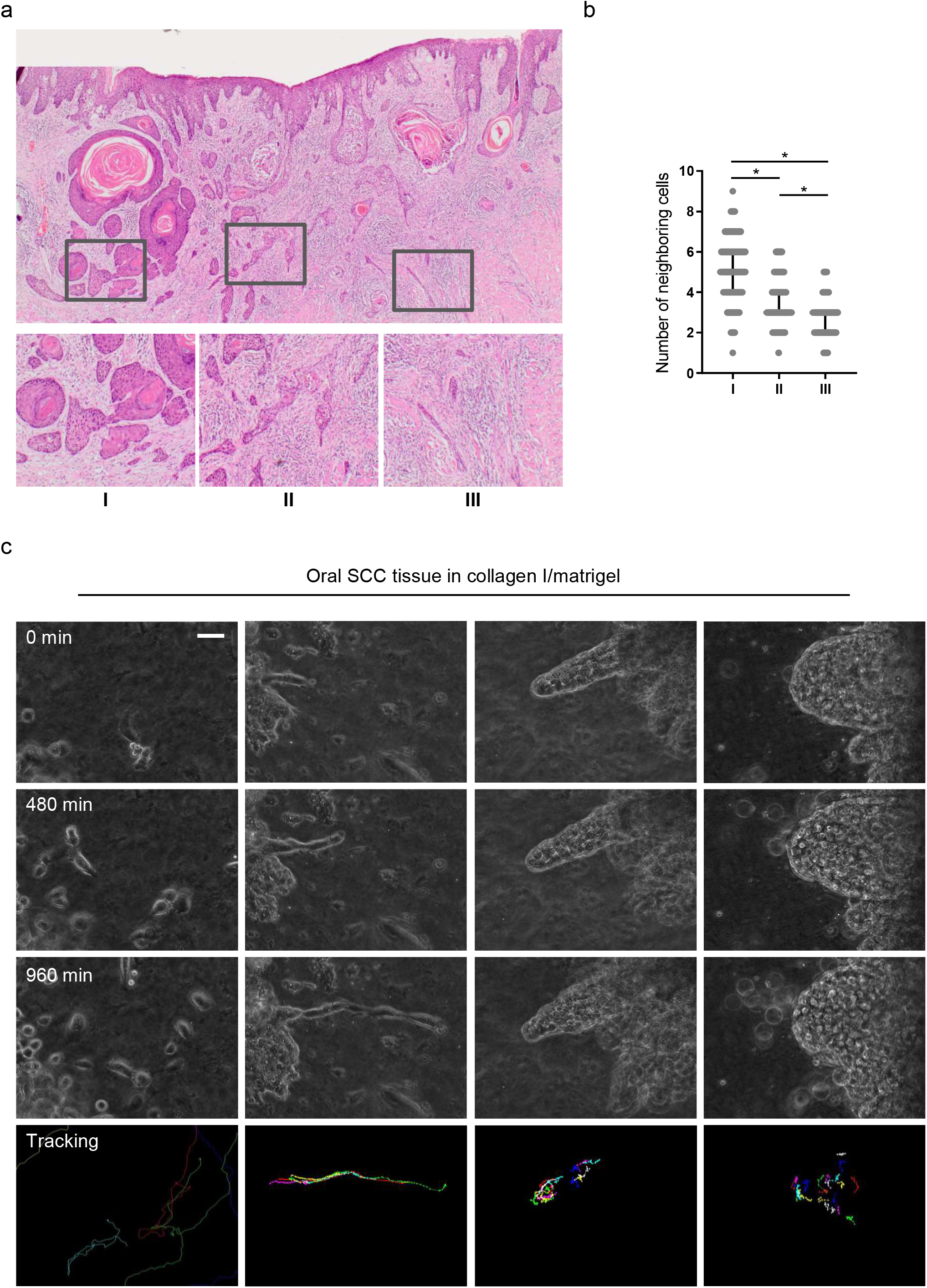
Diversity of collective invasion in squamous cell carcinoma. a) Images show a human invasive head and neck squamous cell carcinoma stained with haematoxylin and eosin. Inset regions show different patterns of collective invasion: I – large rounded clusters, II – intermediate clusters, III – elongated strands only one or two cells wide. b) Plot shows the mean number of cancer cell neighbours for each cell within invasive strands with the morphologies exemplified in panel a) I, II, & III. c) Images show phase contrast microscopy of a human oral squamous cell carcinoma invading into a collagen/matrigel mixture. Scale bar is 100μm. Lower panels show manual tracking of individual cells within the clusters.

To gain insight into the dynamics of squamous cell carcinoma invasion, we performed time-lapse imaging of primary patient explants. Small pieces of tissue, roughly 1 mm^3^ in size, were embedded in a collagen-rich matrix and observed by time-lapse microscopy. Similar to the diversity observed in histological sections, this revealed a variety of behaviours, including single cell ‘follow the leader’ migration through to large ‘dome-like’ multi-cellular invasion, even in samples from a single patient (Figure 1c). Cell tracking revealed that in the larger invading structures there was movement both in the direction of invasion and retrograde back to the main bulk of the explant. The diversity of collective invasion phenotype within a single tumour suggests that the type of collective invasion is not irreversibly determined by early events in the history of the tumour, but can be influenced by variations in cell state that may occur later in tumorigenesis or local environmental differences.

### Generation of an agent-based model of collective cancer invasion

To explore the possible variables responsible for the different collective invasive behaviours observed, we set up both experimental and computational models. Two different experimental settings were implemented. First, an ‘organotypic’ invasion assay in which the SCC cells are cultured as a layer on top of a collagen-rich matrix and exposed to a gas-liquid interface. This recapitulates the early invasion of disease from the epidermis into the dermis (as in the top region of Figure 1a). Second, a ‘spheroid’ assay was used in which the SCC cells are encapsulated in a collagen-rich matrix, mimicking the more confined environment of disease that has already penetrated into the dermis (as in the bottom region of Figure 1a). Alongside these two experimental contexts, we developed a cellular Potts model that incorporated both SCC cells and stromal fibroblasts. The interaction of cancer cells with extracellular matrix and fibroblasts during invasion has been extensively modelled computationally in recent years (Arduino and Preziosi, 2015; Kim et al., 2015; Kumar et al., 2016; Norton et al., 2018; Pally et al., 2019; Sfakianakis et al., 2020). In our 3D model, the voxel size was such that cells typically consisted of 400-800 (~8^3^) voxels. Cell invasion could occur by a cell moving a voxel to a position that was previously occupied by matrix. To determine whether such a change might be favourable, the model included parameters that we anticipated would influence on cancer cell invasion; including cancer cell – cancer cell adhesion, cancer cell – matrix adhesion, cancer cell – fibroblast adhesion, fibroblast – matrix adhesion, cell intrinsic motility, matrix displacement, and matrix proteolysis. The relative influence of these parameters on changes in the position of voxels that defined a cell between time-steps was determined along energy minimization principles, with penalties of differing magnitudes for unfavourable changes in any single parameter (Figure 2a).

**Figure 2:**
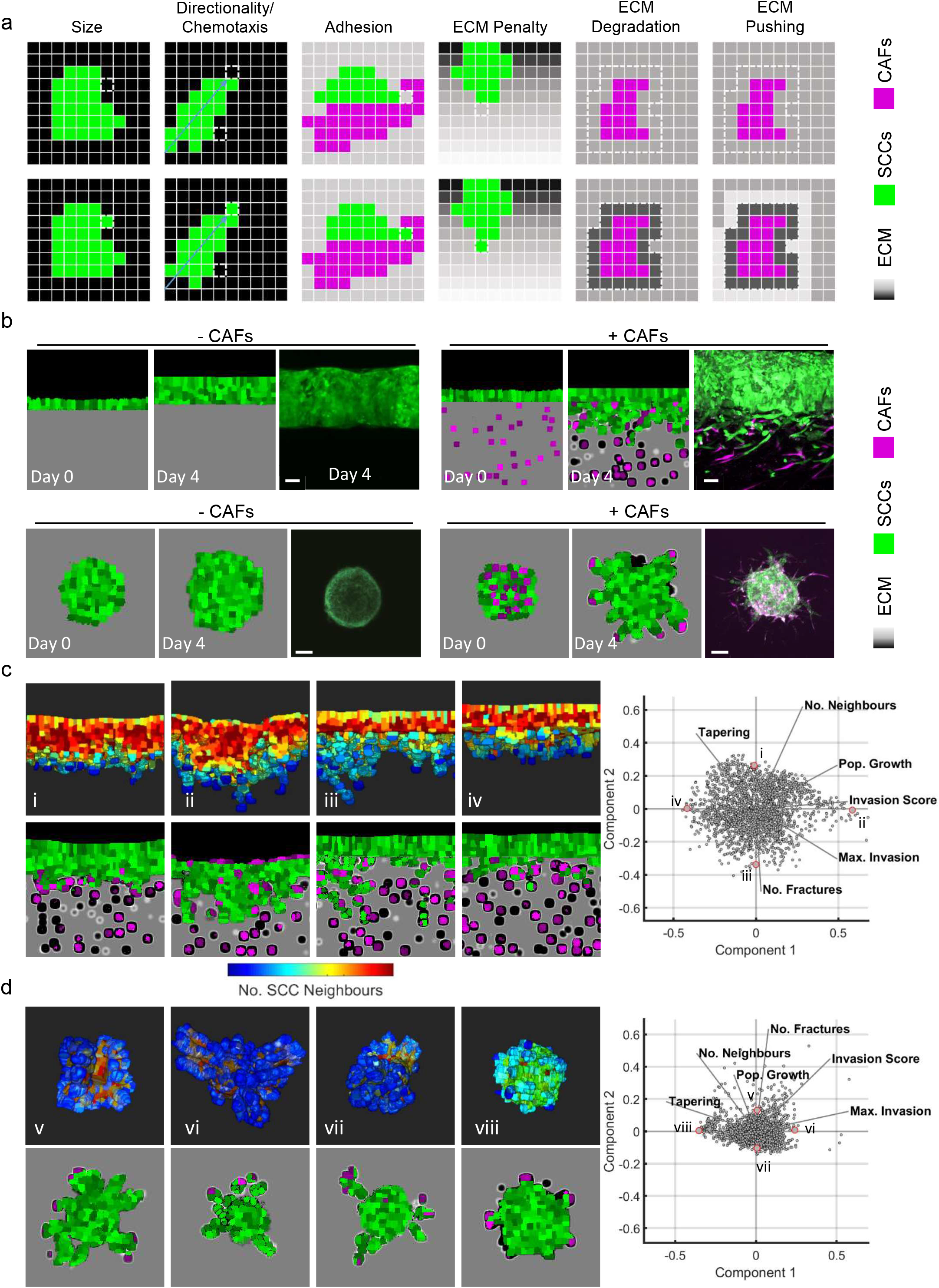
Agent-based modelling recapitulates diversity of collective invasion patterns. a) Images show the key steps and principles driving the agent-based model. At each time step voxels are updated and this can include growth – matrix replaced by cell, directional movement – cell voxel transposition, cell-cell adhesion – increase of voxels at the interface between cells, and ECM remodelling or degradation – change in the ‘quantity’ of matrix in a voxel. Cancer cells are represented in green, fibroblasts in magenta, and matrix in greyscale. b) Images show model outputs (panel columns 1,2,4 &5) next to experimental data when fibroblasts are either absent or present in organotypic models (upper panels) or spheroids (lower panels). Cancer cells are green, fibroblasts are magenta. Scale bar = 100μm. c&d) Diverse patterns of collective invasion are shown in organotypic (c) and spheroid (d) models. PCA plot shows the metrics derived from over 2,000 simulations in the presence of fibroblasts covering variation in cancer cell proteolysis, cancer cell – matrix adhesion, cancer cell – cancer cell adhesion. The additional lines indicate how different metrics contribute to the first two components. The model runs corresponding to the exemplar images are indicated with roman numerals with 3D images heatmapped according to mean number of SCC neighbours (blue low, red high.

Experimental analysis using A431 SCC cells demonstrated that effective invasion required the addition of CAFs and, in both cases, the invasion was almost entirely collective (Figure 2b panels iii & vi). Careful parameterization was performed, including analysis of the relative adhesive properties of the different cells to each other and the collagen-rich matrix used in our assays (Supp. Figure 2a and Supp. Tables 1&2). This enabled the *in silico* replication of the fibroblast-dependent invasion observed in both organotypic and spheroid assays (Figure 2b). In line with previous experimental reports (Gaggioli et al., 2007), the extent of increased invasion scaled with the number of fibroblasts (Supp. Figure 2a).

### *In silico* generation of diverse collective invasion behaviours

Having established an *in silico* model, we then explored parameter space to investigate if different patterns of invasion could be generated by varying the combinations of input parameters. To quantitatively capture the range of invasive behaviours a range of output metrics were collected; including, total invasive extent, maximal invasion, number of cell neighbours, and cell proliferation (Supp. Figure 2b&c). The tapering metric recorded how the number of immediately neighbouring cells varied with the cell’s position in the invasive strand (cells were considered invasive if they had moved beyond the starting position of the interface between cancer cells and the matrix). A uniformly low neighbour number would indicate a long thin strand (Supp. Figure 2cI), a decreasing number of neighbours with increasing invasive would indicate a tapering strand (Supp. Figure 2cII), while a higher number of neighbours would suggest a bulkier form of collective invasion (Supp. Figure 2cIII). A critical function of fibroblasts is to generate permissive tracks for cancer cells to subsequently utilize. To mimic this without the variability generated by the somewhat stochastic behaviour of fibroblasts, we additionally ran simulations with a narrow track that could be permissive for invasion, but no fibroblasts. This confirmed that cancer cells were able to exploit permissive tracks in the extracellular matrix (Supp. Figure 2d). Invasion in this context, termed track invasion score, was quantified based on the extent of matrix remodelling by invading cancer cells with weighting for the distance invaded (Supp. Figure 2e).

The outputs of the model in the presence of CAFs were analysed in two ways: using Principal Component Analysis (PCA) and visual inspection (Figure 2c&d). PCA revealed a wide and continuous spread of invasion patterns, with the first two dimensions of the PCA accounting for 75% (organotypic) and 65% (spheroid) of the variation (Supp. Table 3). Notably, there was no indication of discrete sub-classes of invasive pattern, suggesting a continuous spectrum of invasive behaviours. The continuous spectrum implied by PCA was in line with the range of invasive strand geometries observed in clinical muSCC samples (Figure 1). We additionally, generated visual outputs of the model runs that lay at the edges of the PCA. This revealed diverse patterns of invasion, ranging from large rounded multicellular strands to single cells breaking off the main mass of tumour cells. The diversity in collective invasion observed in the presence of fibroblasts was in contrast to behaviours observed in the absence of CAFs. PCA analysis of the metrics generated from model runs without CAFs shows that the data reduces to a single dimension (Supp. Figure 2f & Supp. Table 3) with remarkably similar behaviour in both organotypic and spheroid data. PCA combining runs with and without CAFs confirmed that fibroblasts boost invasion (Supp. Figure 2f).

### Matrix proteolysis drives strand widening, but not the extent of invasion

Having established that our model could generate diverse types of invasion, we undertook a more systematic analysis of parameter space to determine the contribution of specific parameters to both the extent and pattern of invasion. Figure 3a shows the PCA plots overlaid with shading for the input variable of cancer cell proteolysis, with high levels of proteolysis trending along the vector for number of neighbours in both organotypic and spheroids. Somewhat contrary to expectation, we found that increasing cancer cell proteolysis led to only modestly elevated invasion scores in organotypic contexts. Moreover, the maximum invasive depth did not correlate with matrix proteolysis. Instead the width of the strands (neighbour numbers) increased as a function of proteolysis, especially in organotypic assays. In simulations with low proteolysis, the model predicted thin strands (low neighbour & low tapering scores). To measure the effect of proteolysis on the shape of the invading front of cell clusters, we ran simulations initiated with a cluster of cells and a uniform directional cue, either without the complicating factor of pre-existing tracks or a simple single permissive track. Supp. Figure 3d shows that increasing proteolysis leads to reduced curvature and a ‘pushing’ front in the absence of a track. When a track was present, it was favoured for invasion and interfered with the generation of a pushing front most strikingly at intermediate levels of proteolysis.

**Figure 3:**
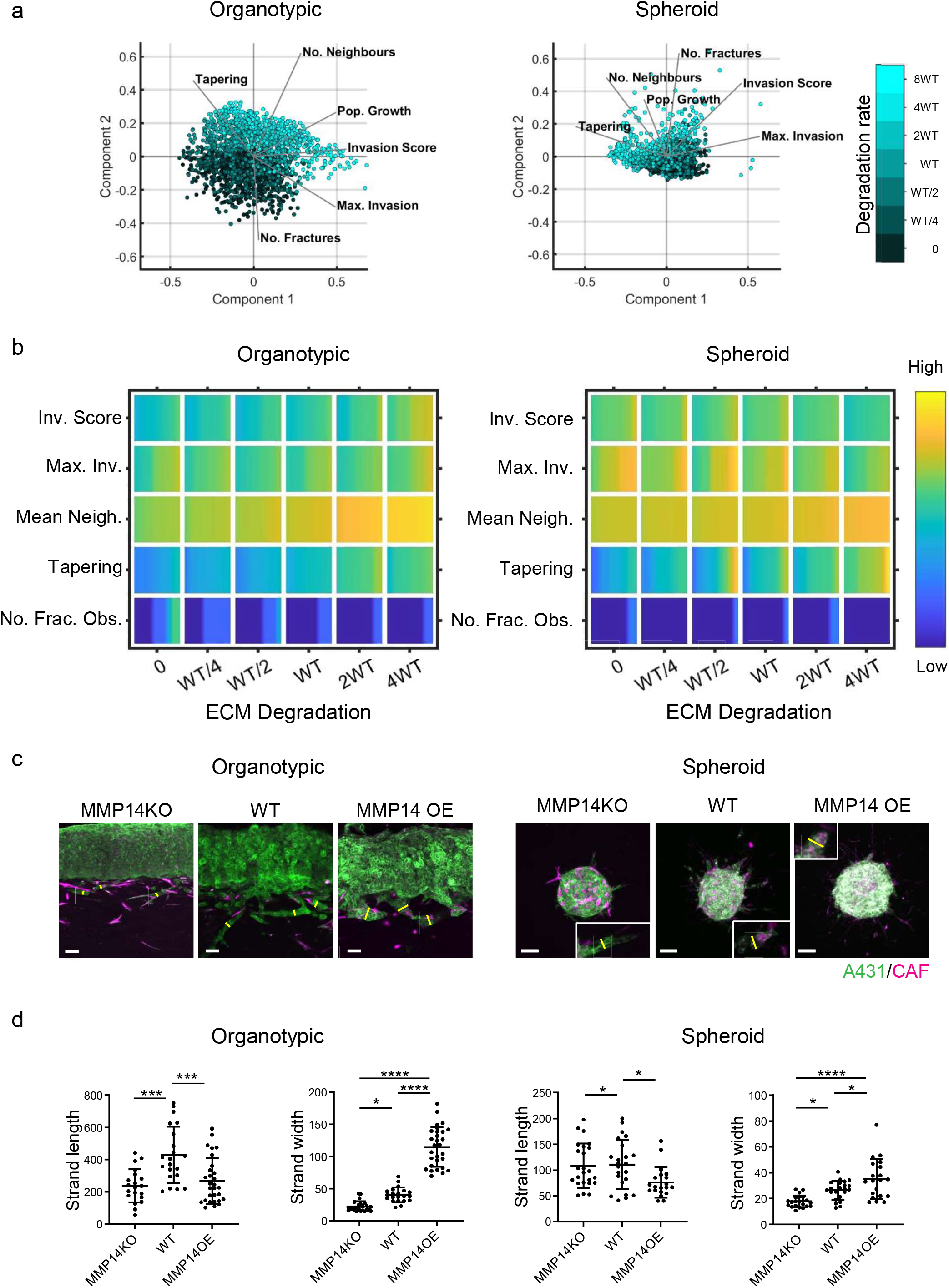
Matrix proteolysis determines strand width, but not the distance invaded. a) PCA plots show the metrics derived from over 2,000 simulations in the presence of fibroblasts covering variation in cancer cell proteolysis with values indicated by the intensity of cyan, cancer cell – matrix adhesion (not colour coded), cancer cell – cancer cell adhesion (not colour coded). b) Heatmaps show how varying the cancer cell proteolysis value (x axis) impacts on different metrics when fibroblasts are included in all simulations. WT indicates the ‘wild-type’ value based on experimental parameterisation using A431 cancer cells. Yellow indicates a high value, dark blue a low value. c) Images show the effect of modulating matrix proteolysis via either MMP14 Crispr KO or MMP14 over-expression in cancer cells (green) both organotypic and spheroid assays including fibroblasts (magenta). Scale bar = 100μm d) Quantification of three biological replicates of the experiment shown in panel c) with strand length and strand width shown – 1 unit is equivalent to 0.52μm.

Analysis of spheroid contexts yielded a different picture; with reduced maximum invasion depth with increasing proteolysis values. Notably, the very highest matrix degradation value yielded significantly lower maximum invasion depth than the intermediate and lowest level. There was less difference in the overall invasion score as increasing proteolysis was linked to slightly wider strands, which counter-balanced the reduction in maximum invasion depth (Figure 3a&b). Both organotypic assays and spheroids without CAFs exhibited low levels of invasion (Supp. Figure 3a&b). Comparative plots of the metrics in simulations with and without CAFs confirm this (note the red colour) and indicate that fibroblasts favour narrower strands (note the blue colour in the neighbour and tapering rows). Overall, cancer cell proteolysis is primarily predicted to regulate strand width in both organotypic and spheroid contexts.

We tested the predictions that cancer cell matrix protease function was linked to width of invasion strands by generating A431 cancer cells that either over-expressed *MMP14,* the major collagen protease, or had it deleted via Crispr/Cas9 editing methods (Supplementary Figure 3c). Figure 3c&d shows that experimentation confirmed the major predictions of our model. In particular, the maximum invasion depth in the organotypic context did not simply increase with MMP14 levels, with strand lengths similar between *MMP14* KO and over-expressing cells. In contrast, the strand width was notably affected by MMP14 levels in both organotypic and spheroid assays (yellow lines in Figure 3c indicate strand width), with KO cells generating thin strands and over-expressing cells generating thick strands. The positive relationship between ECM proteolysis and strand width was particularly strong in organotypic contexts (Figure 3c&d). These results are highly concordant with the model predictions and confirm that MMP14 is a major determinant of the mode of collective cancer cell invasion, but plays little role in determining the maximum distance invaded.

### Cell-cell, but not cell-matrix, adhesion promotes wide invasive strands

We turned our attention to investigate how cancer cell adhesion to either other cancer cells or the matrix influenced the mode of collective invasion. Figure 4a shows PCA plots of invasion characteristics with the strength of cancer cell – matrix adhesion overlaid in green shading. There was no consistent association between cancer cell-matrix adhesion and invasive pattern in the organotypic context, with high adhesion values distributed across the PCA plot. The relationship between cell-matrix adhesion and invasion score was relatively flat, with only very high cell-matrix adhesion values boosting invasion. This prediction is supported by the lack of effect of *ITGB1* deletion on cancer invasion in the experimental organotypic model (Figure 4c&d). In the spheroid context, there was a somewhat stronger association between matrix adhesion and invasion. Minimal invasion was observed in the absence of fibroblasts (Supp. Figure 4a). Intriguingly, the strongest correlation was with the tapering metric that reflects whether strands have a uniform breadth or taper as they invade deeper (Figure 4b – row four). Experiments using *ITGB1* KO A431 cells provided support for this prediction with greater tapering observed in *ITGB1* KO spheroids (Figure 4c&d). Interestingly, and in line with model predictions, this was not observed in organotypic assays (Figure 4b-d).

**Figure 4:**
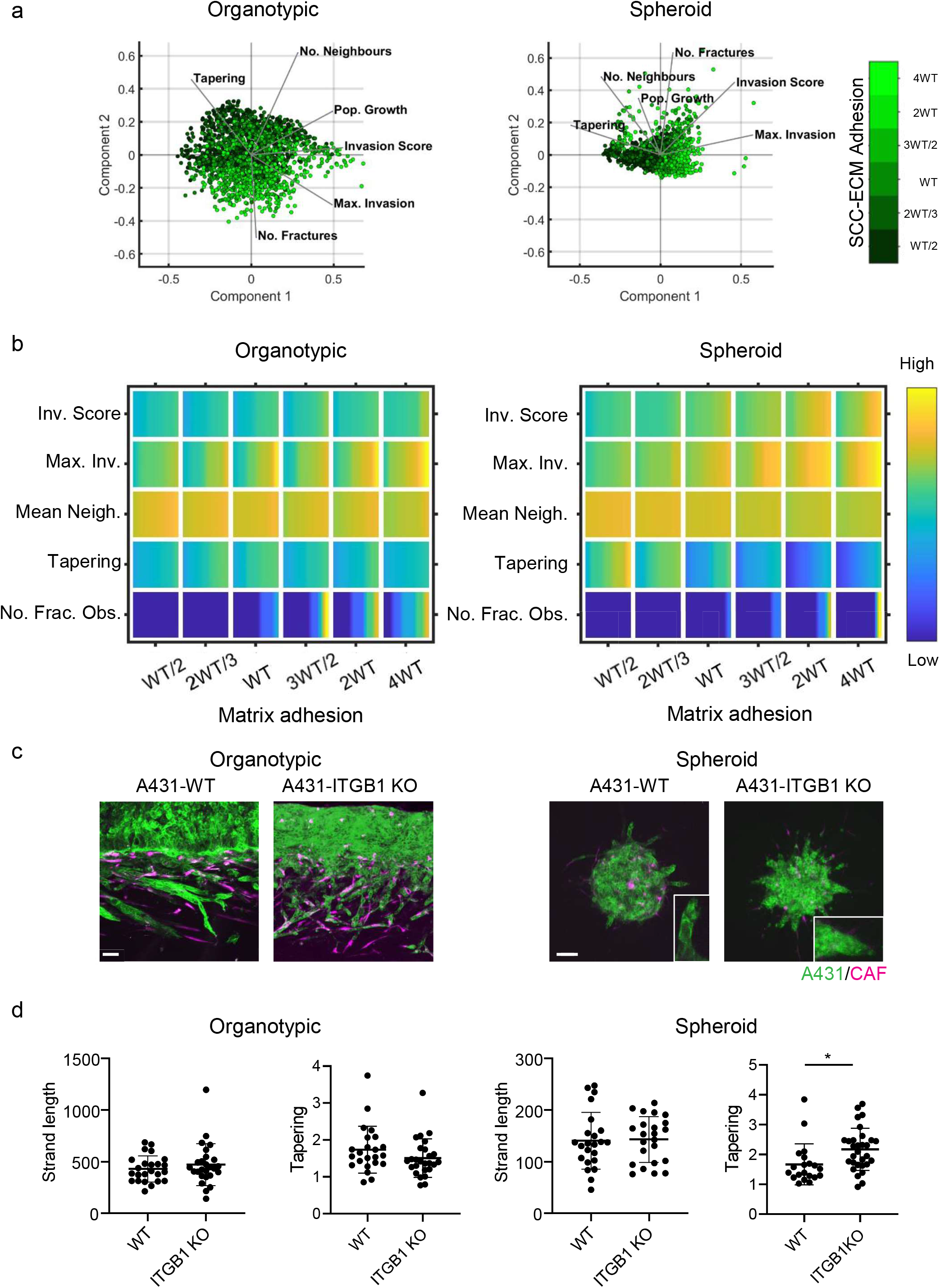
Cancer cell – matrix adhesion modulates the tapering of strands. a) PCA plots show the metrics derived from over 2,000 simulations in the presence of fibroblasts covering variation in cancer cell – matrix adhesion with values indicated by the intensity of green, cancer cell proteolysis (not colour coded), cancer cell – cancer cell adhesion (not colour coded). b) Heatmaps show how varying the cancer cell – matrix adhesion value (x axis) impacts on different metrics when fibroblasts are included in all simulations. WT indicates the ‘wild-type’ value based on experimental parameterisation using A431 cancer cells. Yellow indicates a high value, dark blue a low value. c) Images show the effect of modulating matrix adhesion via Crispr KO of ITGB1 in cancer cells (green) in both organotypic and spheroid assays including fibroblasts (magenta). Scale bar = 100μm d) Quantification of three biological replicates of the experiment shown in panel C) with strand length and tapering metric shown – 1 unit is equivalent to 0.52μm.

Next, we explored the relationship between cancer cell – cancer cell adhesion and invasion when fibroblasts were present (Figure 5a). These analyses yielded several predictions that caught our attention. First, reducing cancer cell – cancer cell adhesion reduced the total invasion score in organotypic assays across relatively large ranges of parameter space (Figure 5a&b – note the association of increasing magenta intensity and invasion score vectors in the PCA plot). This is counter to the widely held view that Epithelial to Mesenchymal Transition (EMT) and increased single cell characteristics promote invasion. Specifically, in organotypic contexts, lower cancer cell – cancer cell adhesion resulted in shorter invasive strands that thinned rapidly as they invaded (this is reflected in the Max. Invasion, Mean Neighbour and Tapering rows in Figure 5b). Once again, little invasion was observed in the absence of fibroblasts (Supp. Figure 5a). Supp. Figure 5b explicitly plots the change in strand width as a function of depth for varying cancer cell – cancer cell adhesion. The simpler context of cell invasion into a thin permissive gap further supported the prediction that cancer cell – cancer cell adhesion is linked to wider invading strands (Supp. Figure 5c). The situation in spheroid assays was more subtle, with increases in invasion only predicted at very high values ≥2WT (Figure 5b). Of note, the Neighbour and Tapering metrics did not vary much depending on cancer cell – cancer cell adhesion. To test these predictions, we generated A431 cells defective in cell – cell adhesion as a result of Crispr-mediated deletion of α-catenin/*CTNNA1* (Supp. Figure 3c). Strikingly, and, in line with the model predictions, these cells lacking adherens junctions were significantly less invasive, both in terms of strand length and strand width, in organotypic assays (Figure 5d). In spheroid assays, loss of α-catenin did not affect strand length and had only a modest effect on strand width (~20% reduction compared to a 60% reduction in width in organotypic assays).

**Figure 5:**
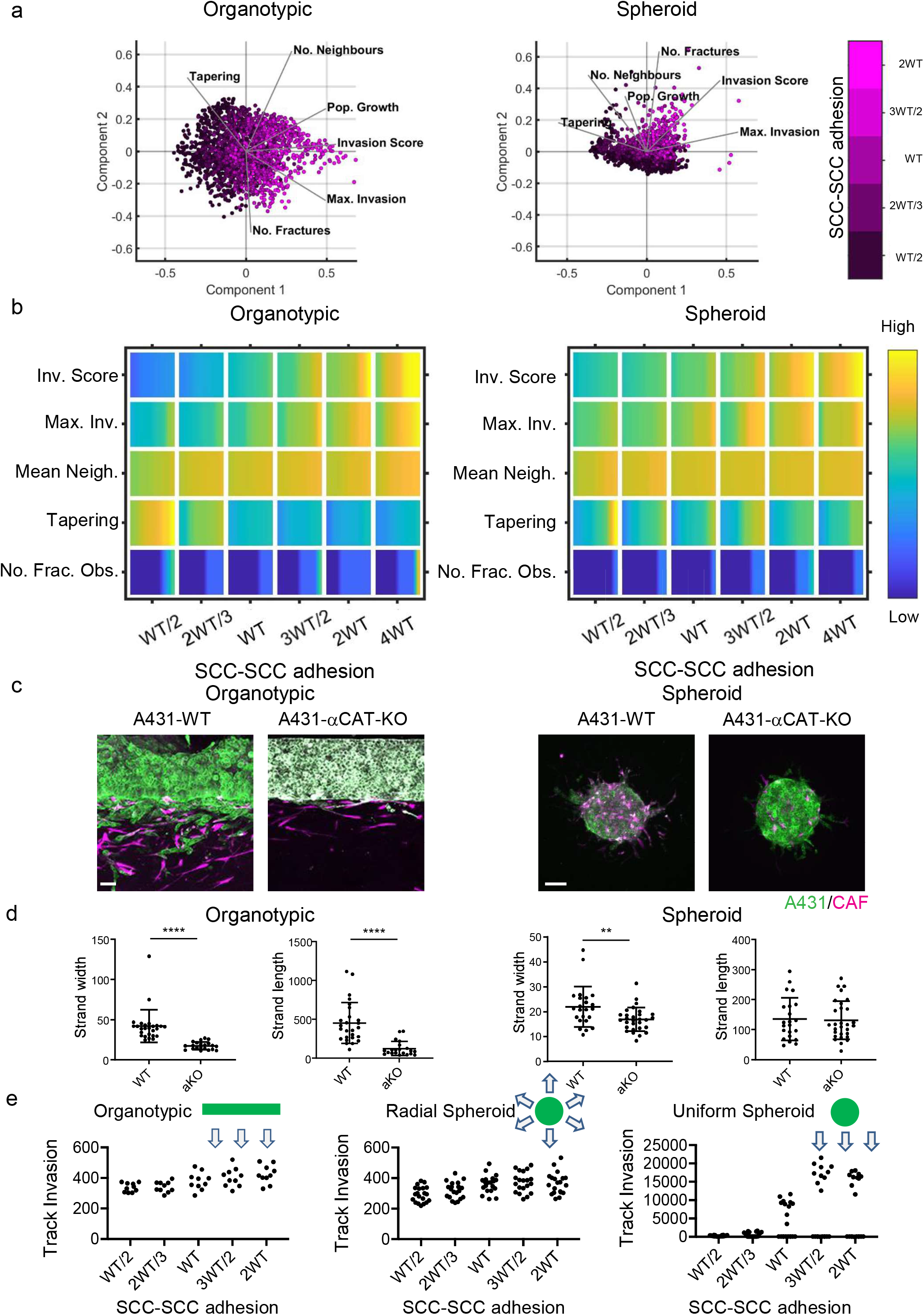
Cancer cell – cancer adhesion is required for efficient invasion in response to uniform directional cues. a) PCA plots show the metrics derived from over 2,000 simulations in the presence of fibroblasts covering variation in cancer cell – cancer cell adhesion with values indicated by the intensity of magenta, cancer cell proteolysis (not colour coded), cancer cell – matrix adhesion (not colour coded). b) Heatmaps show how varying the cancer cell – cancer cell adhesion value (x axis) impacts on different metrics when fibroblasts are included in all simulations. WT indicates the -type’ value based on experimental parameterisation using A431 cancer cells. Yellow indicates a high value, dark blue a low value. c) Images show the effect of modulating cancer cell – cell adhesion via Crispr KO of CTNNA1 in cancer cells (green) in both organotypic and spheroid assays including fibroblasts (magenta). Scale bar = 100μm. d) Quantification of three biological replicates of the experiment shown in panel c) with strand length and strand width shown – 1 unit is equivalent to 0.52μm. e) Plots show the track invasion score with varying cancer cell – cancer cell adhesion in simulations lacking fibroblasts, but with a single permissive track favouring invasion. Cartoons indicate the initial set up of cell positions and the directional cue in the simulation.

### The pro-invasive role of cell-cell junctions depends on a uniform directional cue and supra-cellular coordination of the actomyosin cytoskeleton

The data described above establish an intriguing context dependent role for cell-cell junctions in collective invasion – with a positive relationship between cell-cell adhesion and invasion in organotypic contexts, but not in spheroid contexts. One key difference between these two contexts is that cancer cells in the organotypic context are subject to a uniform gradient of chemotactic cues, whereas in the spheroid context the cancer cells are subject to a radial chemotactic cue. We used our model to test if switching to a uniform chemotactic gradient in the spheroid context would generate a positive relationship between cell-cell adhesion and invasion. Figure 5e quantifies track invasion score in simulations of spheroids with either uniform or radial chemotactic cues. These analyses indicate that cancer cell junctions are favourable for invasion when cells are subject to a uniform directional cue. The importance of junctions only when there is uniform directional cue suggests that it may not be cell-cell adhesion per se that is important, but some linkage between cell-cell adhesions and coordination of collective invasion. Consistent with this idea, cadherin-mediated coordination of actin and myosin dynamics is important for effective collective migration of neural crest cells during cranio-facial development and for border cell migration in the *Drosophila* egg chamber (Geisbrecht and Montell, 2002; Shellard et al., 2018). We hypothesized that a similar mechanism might also underlie the context depend importance of adherens junctions in cancer cell invasion.

Previous work revealed that collectively invading cancer cells have a supra-cellular actomyosin network that enables the coordinated migration of cell groups. Figure 6a confirms control A431 cells exhibit supra-cellular organization of their actomyosin network (Hidalgo-Carcedo et al., 2011). Furthermore, knockout of *CTNNA1* disrupts the formation of a supra-cellular actomyosin network (Figure 6a). To experimentally disrupt the supra-cellular actomyosin network while retaining cell-cell junctions, we utilized two experimental tools, ROCK:ER and ROCK kinase inhibition. The former ectopically boosts actomyosin contractility at cell-cell interfaces, and the latter reduces that activity of the supra-cellular actomyosin belt. Figure 6b&c show that these manipulations have the desired effect on active actomyosin, as determined by pS19-MLC staining (Supp. Figure 6a&b confirm these observations with staining for MYH9). We next tested the effect of these perturbations on A431 *MMP14* over-expressing cells that generate wide invasive strands. Figure 6d&e show that both manipulations reduce the width of invading strands, demonstrating that disrupting actomyosin coordination mechanisms phenocopy loss of adherens junctions with respect to the width of invading strands. Further, the data support a model in which adherens junctions influence invasive pattern by enabling supra-cellular coordination of actomyosin, and not simply determining whether cancer cells are able to maintain contact with one another.

**Figure 6:**
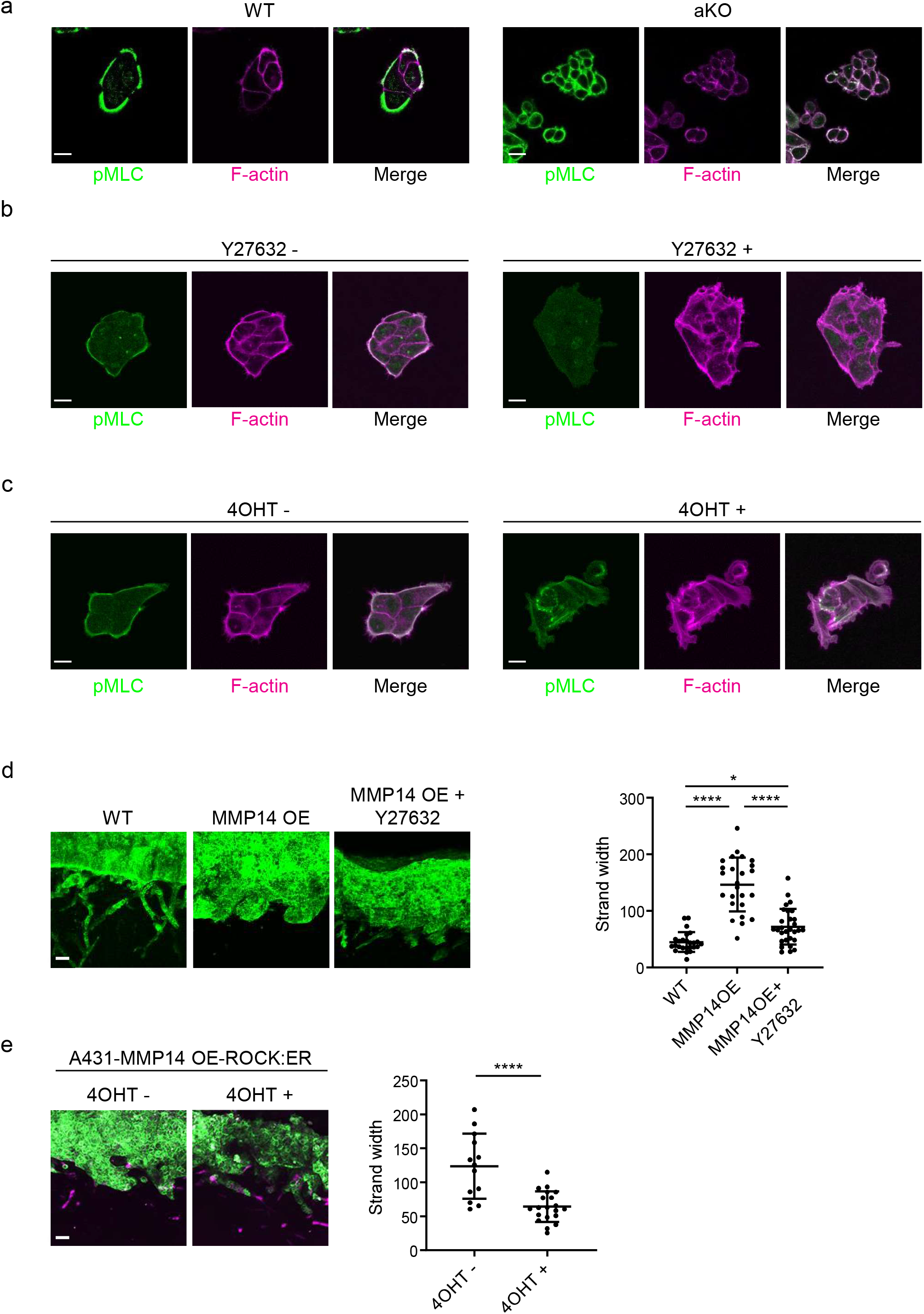
Supra-cellular coordination of actomyosin organisation by cell – cell junctions enables wide invading strands. a) Images show the F-actin (magenta) and active myosin (pS19-MLC – green) networks in control A431 and CTNNA1 KO A431 cells. b) Images show the F-actin (magenta) and active myosin (pS19-MLC – green) networks in control A431 and 10μM Y27632 treated cells. Scale bar = 20μm. c) Images show the F-actin (magenta) and active myosin (pS19-MLC – green) networks in control A431 ROCK:ER and 4-OHT-treated cells. Scale bar = 20μm. d) Images show organotypic invasion assays using control or MMP14 over-expressing A431 cells in the presence or absence of 10μM Y27632. Plot shows the quantification of strand width from three biological replicates – 1 unit is equivalent to 0.52μm. Scale bar = 100μm. e) Images show organotypic invasion assays using MMP14 over-expressing A431 cells additionally engineered to contain ROCK:ER in the presence or absence of 4-OHT. Plot shows the quantification of strand width from three biological replicates. Scale bar = 100μm.

### Protease-driven strand widening requires cell-cell junctions

The analyses above investigate the relationship between individual cancer cell parameters and invasion; we additionally explored how combinations of parameter variations influenced invasive pattern and extent. The data described above argue that, by virtue of their role in coordinating supra-cellular actomyosin, cell-cell junctions would be required for high levels of proteolysis to generate wide invasive tracks. We, therefore, explored the interplay between cancer cell – cancer cell adhesion and proteolysis in determining SCC invasion using both modelling and experimental strategies. Potts modelling predicted that the high neighbour number observed when matrix proteolysis is high would depend upon cell-cell junctions in organotypic assays (note the higher values in the top right regions on the plots in Figure 7ai). Interestingly, this cooperative interaction between proteolysis and cell-cell adhesion was not predicted to influence the extent of maximum invasion, which was dominated by cell-cell adhesion alone (Figure 7aii). These predictions were supported by experimentation: deletion of *CTTNA1* prevented the formation of wide invasive strands by *MMP14* over-expressing A431 cells in the organotypic invasion assays (Figure 7c), with more subtle effects observed in the spheroid assays (Figure 7a,d&e). Supp. Figure 3a&b indicate that CAFs favour narrower invasive strands; therefore, to more fully explore how ECM proteolysis and cell-cell adhesion co-ordinately determine the geometry of collective invasion, we revisited simulations, without CAFs, designed to monitor the curvature of the invading cell cluster (Supp. Figure 3d). Supp. Figure 7a&b show that if both ECM proteolysis and cell-cell adhesion are high then a broad, virtually flat, invasive front is generated. Reducing either proteolysis or cellcell adhesion leads to increased curvature. Together, these analyses establish that a broad ‘pushing’ front of invasion requires both high proteolysis and high cancer cell – cancer cell adhesion.

**Figure 7:**
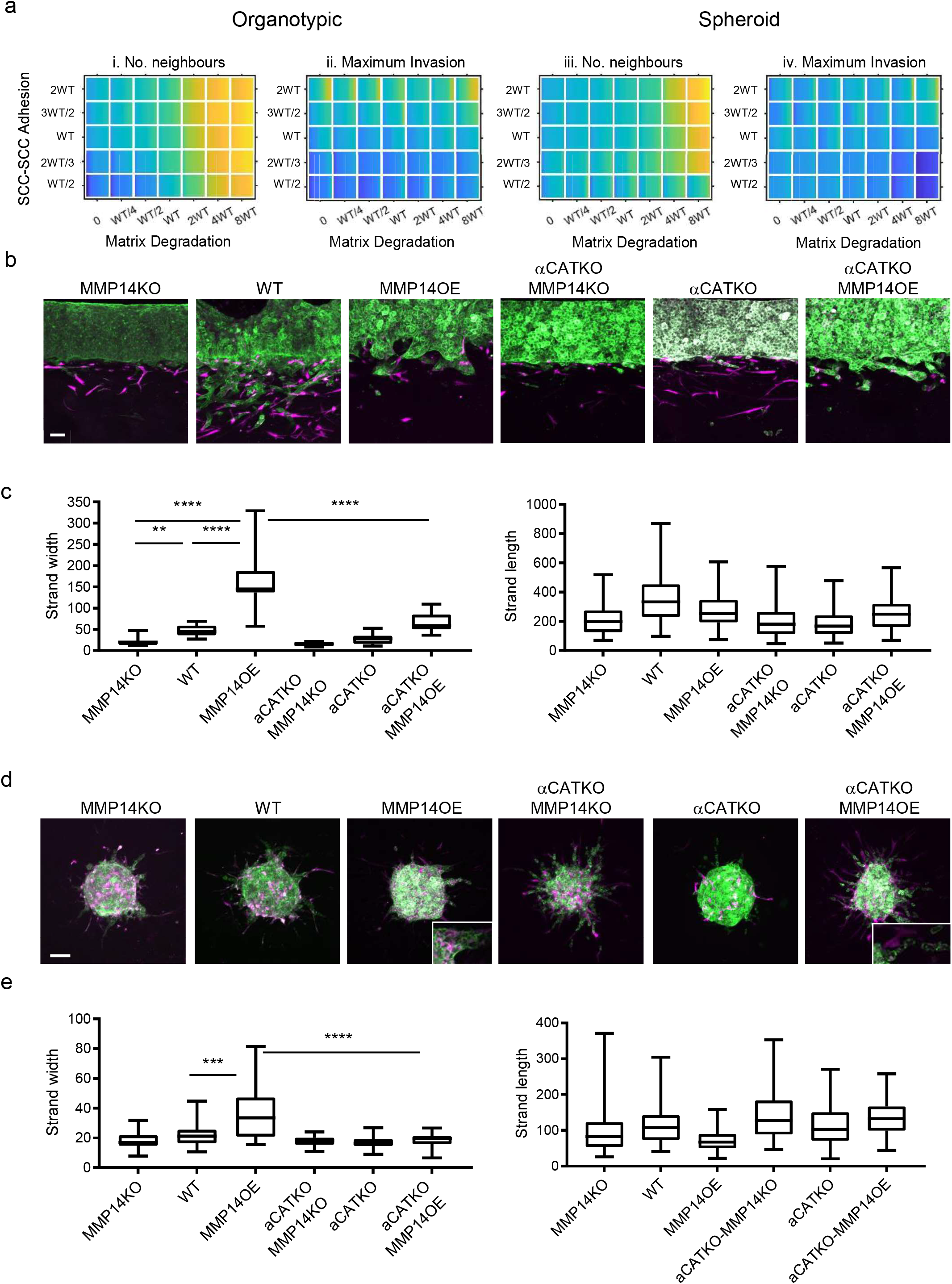
Proteolysis-driven strand widening requires adherens junctions. a) Heatmaps show how varying the matrix proteolysis (x-axis) and cancer cell – cancer cell adhesion value (y axis) impacts on different metrics when fibroblasts are included in all simulations. WT indicates the ‘wild-type’ value based on experimental parameterisation using A431 cancer cells. Yellow indicates a high value, dark blue a low value. b) Images show the effect of combinatorial modulation of matrix proteolysis and cancer cell – cell adhesion via Crispr KO of CTNNA1 and/or MMP14 and/or MMP14 over-expression in cancer cells (green) in both organotypic assays including fibroblasts (magenta). Scale bar= 100μm. c) Quantification of three biological replicates of the experiment shown in panel b) with strand length and strand width shown – 1 unit is equivalent to 0.52μm. d) Images show the effect of combinatorial modulation of matrix proteolysis and cancer cell – cell adhesion via Crispr KO of CTNNA1 and/or MMP14 and/or MMP14 over-expression in cancer cells (green) in both spheroid assays including fibroblasts (magenta). e) Quantification of three biological replicates of the experiment shown in panel d) with strand length and strand width shown. Scale bar = 100μm.

### Strand widening is coupled to cancer cell growth

While the focus of our analysis has been the pattern of invasion, the widening of tracks might also represent a mechanism for generating additional space for cell growth in confined environments. As proliferation is a feature of our model, we additionally investigated whether cancer cell growth might be impacted as a result of change in cancer cell – cancer cell adhesion and proteolysis. Interestingly, the vectors reflecting cell growth and neighbour number in the PCA analysis were closely aligned (Figure 2c&d). This alignment was particularly pronounced in the spheroid simulations lacking fibroblasts (Supp. Figure 2f). Given that we have established matrix proteolysis and cancer cell – cancer cell adhesion as the major determinants of neighbour number and strand width, we therefore investigated the relationship between these parameters and cell growth. Neither was predicted to have a strong effect on cell growth in organotypic assays, either in the presence or absence of CAFs (Figure 8a). In contrast, a strong positive relationship between proteolysis and growth was predicted in the context of spheroids lacking CAFs (Figure 8a). Cancer cell – cancer cell adhesions were also predicted to make a positive contribution to growth, albeit smaller than the effect of proteolysis (Figure 8a). We proceeded to test these predictions experimentally. Manipulation of *MMP14* and *CTTNA1* had minimal effect on cell growth in unconfined 2D culture conditions (Supp. Figure 8a). Figure 8b&c confirm that both proteolysis and cancer cell – cancer cell adhesion are required for effective cell growth in 3D collagen matrices. Moreover, the positive effect of boosting proteolysis required cell-cell adhesions (Figure 8b&c compare MMP14 OE with αCATKO MMP14 OE). Ectopic activation of ROCK2, which disrupts cytoskeletal cohesion in cell clusters, also reduced growth in 3D collagen; further reinforcing the link between the determinants of invasive strand width and cancer cell growth.

**Figure 8:**
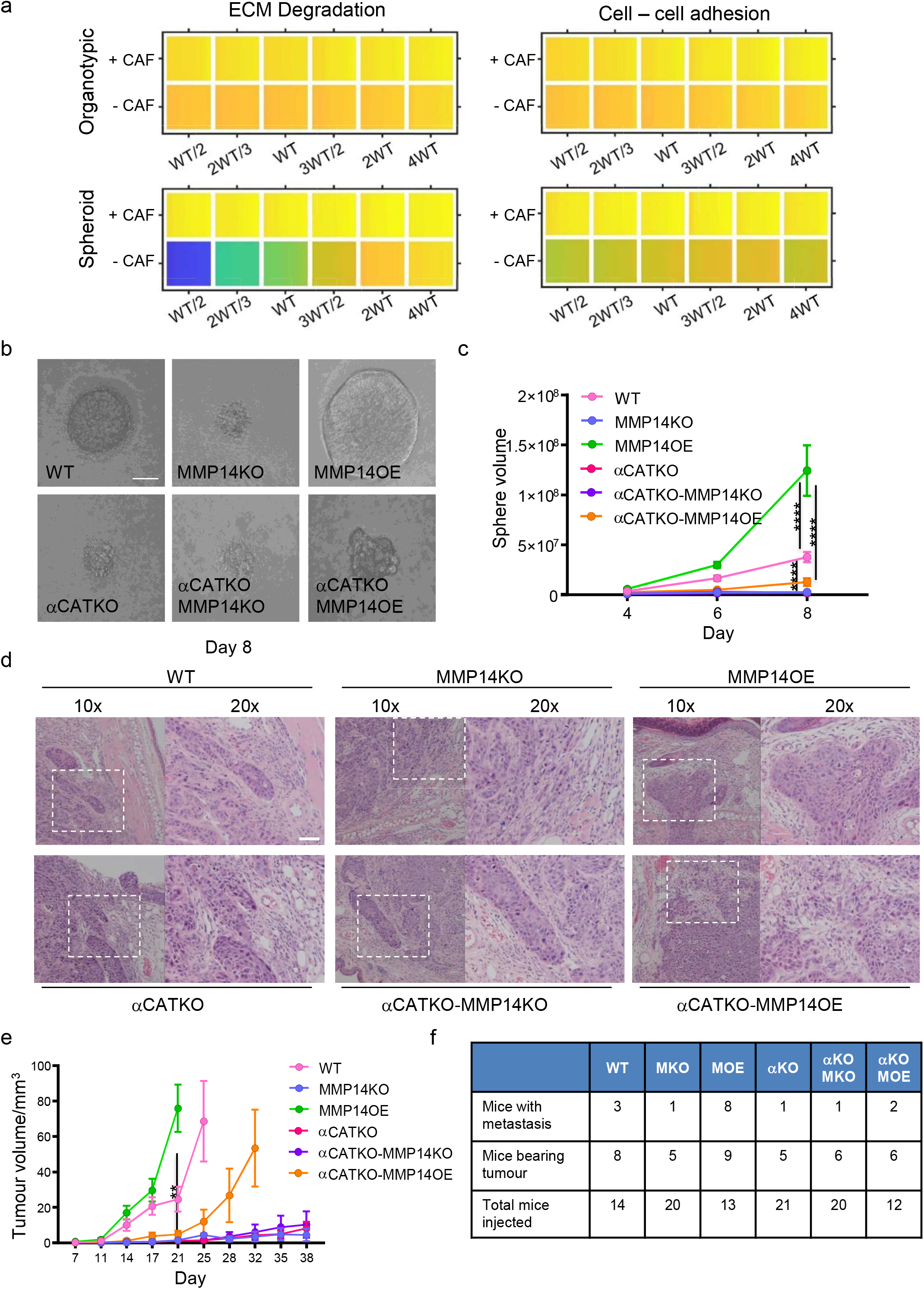
Strand widening is linked to tumour growth and metastasis. a) Heatmaps show how varying the matrix proteolysis (left) or cancer cell – cancer cell adhesion value (right) impacts on predicted cell growth in the presence or absence of fibroblasts. WT indicates the ‘wild-type’ value based on experimental parameterisation using A431 cancer cells. Yellow indicates a high value, dark blue a low value. b) Phase contrast images show the growth of cancer cell colonies with the indicated manipulations of MMP14 and CTNNA1 after 8 days surrounded by matrix. Scale bar = 50μm. c) Plot shows quantification of the growth assay shown in b). Data from three biological replicates. d) H&E images are shown on tumours growing the ears of mice with the indicated manipulations of MMP14 and CTNNA1. Scale bar = 50μm. e) Plot shows quantification of A431 tumour growth with the indicated manipulations of MMP14 and CTNNA1. f) Table shows quantification of mice with primary tumours and mice with lymph node metastases when injected with A431 cells with the indicated manipulations of MMP14 and CTNNA1. The total number of mice for each condition also applies to the data plotted in e).

### Protease-driven tumour growth and lymph node metastasis requires cell-cell junctions

Finally, we sought to test whether key findings of our integrated *in silico* and *in vitro* analysis also applied in an *in vivo* context with a heterogeneous environment including a greater diversity of stromal cell types not included in our model. A431 cells engineered to have different levels of *MMP14* and *CTTNA* levels were injected into the dermal space within the ears of mice. This anatomical location was chosen because the dermis represents the first tissue that squamous cell carcinoma invade into and cells can spread from the dermis to local lymph nodes, which reflects the clinical spread of the disease. This environment is spatially confined with some fibroblasts in addition to thin layers of fat, cartilage, and muscle. It was not possible to include all these additional factors with appropriately controlled parameterization. Therefore, we concentrated on validating the relationship between matrix proteolysis, cancer cell-cell adhesion, and invasive spread *in vivo.* In addition, if stromal support, such as that provided by fibroblasts is limited then the mechanisms that promote wide invasive strands also favour growth. To test these ideas, we injected A431 cells with combinations of MMP14 and α-catenin manipulations into the intradermal space of mouse ears. This environment is spatially restrictive with lymphatic drainage to local lymph nodes. Of note, MMP over-expressing cells generated tumours with particularly wide, bulging, margins (Figure 8d). Strikingly, there was a strong correlation between the levels of MMP14 and tumour growth (Figure 8e). Metastatic spread to lymph nodes also correlated with MMP14 levels, which is in line with previous reports (Bartolomé et al., 2009; Devy et al., 2009; Wang et al., 2021). Notably, and in contrast to the prevailing dogma, reducing cancer cell – cancer cell adhesion did not lead to a more aggressive tumour phenotype, but reduced both tumour growth and very few mice were observed to have lymph node metastases (Figure 8f). This could be partly compensated by over-expression of MMP14, suggesting that a defect in ‘space’ generation might underpin the defect in the *CTNNA1* KO cells. However, the growth and lymph node metastasis of *MMP14 o.e./CTNNA1* KO cells was reduced compared to the *MMP14* o.e. cells (Figure 8f), indicating that the tumour promoting effect of elevated MMP14 levels depends on cell-cell adhesion. Together, these analyses demonstrate that MMP14-driven matrix proteolysis promotes invasion in wide collective units and tumour growth in spatially confined contexts. Furthermore, the widening of invasive units, tumour growth, and lymph node metastases depends upon adherens junction-mediated supra-cellular coordination of the actomyosin network.

## Discussion

The combined computational and experimental analysis of collective cancer cell invasion presented here raises several findings that warrant further consideration. Although, matrix proteolysis was broadly associated with higher levels of invasion (Castro-Castro et al., 2016; Egeblad and Werb, 2002), it was not a simple linear relationship (Supp. Figure 8d). Most notably, high proteolysis reduces the maximal extent of invasion but increases the strand width in both the model and experiments. The ability of cells with high levels of proteolysis to generate space means that there is less pressure to constrict cells into longer thinner strands. The importance of space limitation for effective invasion is underscored by the reduced invasion observed when spheroids have a ‘choice’ between invasion and spreading over an unimpeded matrix layer. High proteolysis essentially reduces the space limitation. This is also linked with high levels of proliferative capacity in 3D environments. Our analysis demonstrated that this growth effect was clearly observed in vivo. *MMP14* over-expressing tumours grew and metastasized aggressively. This argues that SCC cells invading in thick strands are efficient at metastasis. Crucially, the aggressive behaviour of *MMP14* over-expressing cells is reduced by depletion of α-catenin. This argues strongly against a single cell form of migration being optimal for lymph node metastasis of SCC cells. The importance of adherens junctions for efficient metastasis is increasingly appreciated, this work suggests that one advantage of both adherens junctions and matrix proteases in collective migration is that the cells remain in a state capable of generating the space required for growth.

*In silico* analysis revealed that reducing cancer cell-ECM adhesion had a minor effect on determining the mode of invasion. Experimentation using *ITGB1* knock-out cells supported this analysis and notably confirmed the relationship between strand tapering and cancer cell-ECM adhesion. Moreover, unless cancer cell-ECM adhesion was very strong, the relationship between this variable and extent of invasion was rather weak in both organotypic and spheroid assays. Broadly, these data are consistent with the integrin-independence of amoeboid forms of migration in 3D and hint at a role for either adhesion forces mediated by the glycocalyx or a role for outward forces that enable a ‘chimneying’ type of migration. However, it is worth noting that we only targeted *ITGB1* experimentally. In the future, it will be interesting to explore the effect of targeting other integrins and cell-matrix adhesion molecules, and increasing cancer cell-ECM adhesion to very high levels. Cancer cell-cell adhesions exert a greater influence on collective cancer invasion than cell-ECM adhesions (Supp. Figure 8d). Intriguingly, the positive role of cancer cell-cell adhesions was most pronounced in simulations with a uniform chemotactic gradient. We propose that this reflects a crucial role of cell-cell adhesions in coordinating a supra-cellular actomyosin cytoskeleton in collectively invading clusters. This is likely to involve coordination of cell polarity complexes at sites of cell-cell contact. Interestingly, loss of cell-cell junctions was not sufficient to promote a clear switch to single cell invasion. This is likely due to the lack of available space in 3D contexts. This observation is consistent with *CDH1* deficient tumours, such as invasive lobular carcinoma of the breast and some gastric cancers, typically showing thin strand-like patterns of invasion.

To conclude, our integrated *in silico* and experimental approach reveals some of the key determinants of the mode of collective cancer invasion. Broad pushing fronts are associated with high matrix proteolysis and strong cancer cell-cell junctions and a lower dependence on cancer-associated fibroblasts. Reducing either proteolysis or cancer cell-cell adhesions leads to thinner invasive strands, with cell-matrix adhesions tuning strand tapering. We observe and experimentally demonstrate an unexpected linkage between the mechanisms that promote the widening of invasive strands and ability of cancer cells to grow when surrounded by ECM.

## Materials and Methods

### Experimental

#### Cell culture

Human vulval CAFs are described in Gaggioli et al. (Gaggioli et al., 2007). CAFs were cultured in DMEM supplemented with 10% FBS and 1% insulin–transferrin–selenium (Invitrogen, no. 41400-045) and 100U/ml penicillin, 100μg/ml streptomycin. Human vulval squamous cell carcinoma cell line A431 cells were grown in DMEM supplemented with 10% FBS, 100U/ml penicillin, 100μg/ml streptomycin. For ROCK inhibitor treatment cells were treated with 10μM Y27632.

#### Stable cell lines

E-cadherin, E- and P-cadherin, alpha-catenin, *MMP14* KO A431 cells were generated by CRISPR-Cas9 as previously described (Labernadie et al., 2017). Briefly, pX458 vectors encoding gRNA sequences were transfected into A431 cells and single GFP positive cells were sorted into 96-well plate 2 days after transfection. Cells were grown for 2 weeks and KO was checked by western blot and sequencing of genome DNA. For *MMP14* overexpressing cells, A431 cells were transfected with pMMP14-mCherry (generous gift from Dr. Machesky at CRUK Beatson Institute) and selected by G418 for 2 weeks. mCherry positive cells were sorted by flow cytometry. Stably labelled A431 cells and CAFs were obtained by infecting lentivirus containing fluorescent protein gene. 293FT cells were transfected with pCSII-mCherry-CAAX, pCSII-ECFP-CAAX or pCSII-KEIMA-CAAX construct and lentiviral RRE, REV and VSVG encoding plasmids (5μg each) by Xtremegene HP (Roche) according to the manufacturer’s recommendation. Resulting supernatant containing lentivirus was then infected to target cells.

#### Western blotting

Cells were lysed with Laemmli sample buffer containing 2.5% β-mercaptoethanol and heated at 95 °C for 5 min. Samples were loaded to 4-15% polyacrylamide gels (Bio-Rad) for electrophoresis. Proteins were then transferred to a PVDF membrane (Merck), which was blocked with 5% dry milk, Tris buffered saline, 0.2% Tween, and incubated with primary antibodies (overnight at 4° C) followed by secondary antibodies (1:10000) for 1 h at room temperature. Proteins were detected by using Luminata Crescendo (Merck) and LAS600 (GE Healthcare). The following antibodies were used: anti-MMP14 rabbit monoclonal (1:1000, EP1264Y, Abcam), anti-alpha-catenin rabbit monoclonal (1:1000, EP1793Y, Abcam), anti-Vimentin mouse monoclonal (1:1000, 1A4, Sigma), anti-Fibronectin rabbit polyclonal (1:1000, Sigma) and anti-actin mouse monoclonal antibody (1:2000, AC-40, Sigma).

#### Spheroid invasion assay

A431 and CAF cells were detached from the cell culture dishes with trypsin and re-suspended in sterile 0.25% methylcellulose solution in DMEM. The cellulose solution contained a 1:1 ratio of A431 and CAF cells at a concentration of 1 × 10^5^ cells/ml. Twenty microliter droplets were plated onto the underside of a 10 cm culture dish and allowed to form spheroids in a 37 °C incubator overnight. The spheroids were then embedded in a collagen l/Matrigel gel mix at a concentration of approximately 4 mg/ml collagen I and 2 mg/ml Matrigel (BD Bioscience) in 24-well glass-bottomed cell culture plates (MatTek) on a 37 °C hot block. The gel was incubated for at least 30 min at 37 °C with 5% CO2. The gel was covered with DMEM media containing 10% FCS. Sixty hours later, the spheroids embedded in the gel were washed with PBS and then fixed for 30 min at room temperature with 4% paraformaldehyde. The spheroids were then imaged with an inverted Zeiss LSM780 at a magnification of ×10, ×20 and ×63. z stack images spanning 100–150 μm were collected and image stacks were processed by ZEN software (Carl Zeiss) to yield maximum-intensity projections.

For quantification of the images, strand length and width were measured using Fiji software. Strand tapering was calculated by the following formula: strand width at 20% from the root/ strand width at 80% from the root.

#### Organotypic invasion assay

Organotypic invasion assays were performed as previously described^27^. Briefly, collagen I (BD Biosciences cat. no. 354249) and Matrigel (BD Biosciences cat. no. 354234) were mixed to yield a final collagen concentration of 4 mg/ml and a final Matrigel concentration of 2 mg/ml. After the gel had been left to set at 37 °C for 1 h, mixture of 5 × 10^5^ A431 cells and 5 × 10^5^ vulval CAFs were plated on the top in complete medium. Twenty-four hours later, the gel was then mounted on a metal bridge and fed from underneath with complete medium (changed daily). After 6 days the cultures were fixed with 4% PFA plus 0.25% glutaraldehyde in PBS and imaged using Zeiss LSM780 at a magnification of ×10 and ×20. z stack images spanning 100-150 μm were collected and image stacks were processed by ZEN software to yield maximum-intensity projections.

For quantification of the images, strand length and width were measured using Fiji software. Strand tapering was calculated by the following formula: strand width at 20% from the root/ strand width at 80% from the root.

Wound healing assay: 4×10^4^ cells in 70μL medium were seeded into each well of 2-well culture insert (Ibidi) and cultured overnight. After removing culture insert complete medium was added to the dish and images were taken at 0, 9 and 24h. Empty area was measured using Fiji and the results of 9 and 24h were normalised to that of 0h.

#### Proliferation assays

2D assay – 5×10^4^ cells were seeded in 24 well plate and the number of cells was counted everyday using Countess II automated cell counter (Thermo Fisher Scientific). Results were normalised to day 1. 3D ‘confined’ assay – SCC cells were mixed in collagen I/Matrigel at a concentration of 3×10^3^/ml, and 100μL of the mixture was put in 96 well plate and incubated for an hour at 37 degree. After the incubation 150μL of complete medium was added to each well. Images of growing cells were taken at indicated time points with EVOS FL microscope system (Thermo Fisher Scientific).

#### Immunostaining

Cells were fixed with 4% paraformaldehyde for 10 min and permeabilized in 0.1 % Triton X-100 for 10 min. Cells were blocked in 1% BSA for 1 h before incubation with primary antibodies at 4 degree overnight. After incubation the appropriate fluorescence-conjugated secondary antibodies for 1 h, cells were washed with PBS. Images were acquired with an inverted Zeiss LSM780 at a magnification of ×20 and ×63.

#### *In vivo* tumour growth

Cells were detached from culture flask and resuspended in 4mg/ml Matrigel/PBS at a concentration of 2.5×10^7^. Twenty micro litre of cell suspension was injected into mouse ear intradermis using 31G needle (BD). The tumour size was measured every 3-4 days using caliper until it reached 0.6mm in diameter. At the end point mice were sacrificed and the tumour samples were fixed with 4% PFA overnight and processed by standard methods for haematoxylin/eosin staining. Cervical lymph node was taken out and analysed for metastatic seeding.

### Computational

#### Cellular Potts Model

Detailed information on mathematical background and C++ coding implementation for each cellular mechanism within the model can be found in Supplementary Methods.

#### Simulation Quantification

For each simulation outcome of interest (for example, each combination of parameter values) ten CC3D simulations were run to generate invasion metrics. All invasion metrics were calculated in MATLAB (version 2019a). Unless stated, all invasion metrics were recorded at day four. Invading cells are classed as all cells beyond the tumour interface at day zero. Maximum invasion is given by the maximum distance of any invasive SCC centroid to the initial tumour interface. Invasion Score is equal to the total number of invasive SCCs multiplied by the mean distance of invasive centroids to the initial tumour interface. For mean number of SCC neighbours, tapering and number of fractured objects the bulk tumour mass at day four is found. The mean number of SCC neighbours is calculated for all cells in the bulk tumour mass that are invading. For these cells the gradient of line of best fit between the number of neighbours and distance from initial tumour interface is calculated to give the tapering metric. Fractured objects are defined as objects unconnected to the bulk tumour mass and containing at least one SCC. The number of these distinct objects is counted for the fractured object metric. For cell growth the total number of SCCs versus time is recorded and an exponential fitted to the resulting curve. For the combination of very large SCC-degradation (8WT), SCC-SCC adhesion (2WT) and SCC-ECM adhesion (4WT) in the presence of CAFs, spheroids can become hollow and break apart. In such circumstances, there is no bulk tumour mass resulting in a mean number of neighbours of zero for the main tumour mass and a large number of fractured objects. There are 4 instances of this and these data have been removed (leaving 6 simulations for this region of parameter space) prior to PCA analysis and heatmap generation. The track invasion score is taken at day 5. It is calculated by finding all points around a permissive track, beyond the initial tumour boundary where the ECM density is 0.75 or below (initial condition set to 1). These points are then weighted according to their distance from the boundary and then summed. For the spheroid both permissive tracks, either side of the initial tumour mass are quantified. Track width is calculated as the maximum width of the invading strand. Strands that are either non-invasive or where the entire tumour mass has invaded uniformly do not record track width values. Curvature is quantified on day seven. The leading invasive edge is reduced to two one-dimensional signals in x-z and y-z for mid-points of y and x respectively. Each one-dimensional signal is then smoothed with a smoothing window of 50 pixels. The LineCurvature2D function (Dirk-Jan Kroon (2021). 2D Line Curvature and Normals, MATLAB Central File Exchange. Retrieved 9 November 2021 (https://www.mathworks.com/matlabcentral/fileexchange/32696-2d-line-curvature-and-normals)) is used to calculate curvature for each signal and the average taken. For all heatmaps, for each box the x-axis represents the percentiles from 0.5 to 99.5 (left to right) of all ten simulations for that outcome of interest.

## Supporting information

Supplementary CC3D Code Walkthrough

## Acknowledgements

We thank the Francis Crick Institute Advanced Light Microscopy facility, Cell Services, Flow Cytometry, the Biological Research facility and Experimental Histopathology facility for scientific and technical support. We thank Dr. Laura Machesky for kindly providing an *MMP14* plasmid. We are grateful to lab members, past and present, for help and advice throughout this work.

The work was funded by the Francis Crick Institute which receives its core funding from Cancer Research UK (FC001144, FC001003), the UK Medical Research Council (FC001144, FC001003 FC001144) and the Wellcome Trust (FC001144, FC001003). T. K. was supported by JSPS Kakenhi grant number JP19K21262, Marie-Curie action (HeteroCancerInvasion no. 708651), The Uehara Memorial Foundation and Kitasato University Research Grant for Young Researchers.

---

**Supplementary Figure 1: Diversity of collective invasion in squamous cell carcinoma**

a) Images show a human invasive head and neck squamous cell carcinoma stained with haematoxylin and eosin (left) and anti-α-SMA (right). Inset regions show different patterns of collective invasion: I – large rounded clusters, II – intermediate clusters, III – elongated strands only one or two cells wide. b-d) Images show three further human invasive head and neck squamous cell carcinoma stained with haematoxylin and eosin and cytokeratin (bottom right). Inset regions show different patterns of collective invasion: I – large rounded clusters, II – intermediate clusters, III – elongated strands only one or two cells wide. e) Plot shows the average number of cancer cell neighbours each cell within invasive strands with the morphologies exemplified in panels a) & b-d), respectively. ***p<0.0005, ****p<0.0001. One way ANOVA with post-hoc multiple comparisons.

**Supplementary Figure 2: Agent-based modelling recapitulates diversity of collective invasion patterns**

a) Images show model outputs next to experimental data for increasing numbers of fibroblasts. Cancer cells are green, fibroblasts are magenta. Scale bar = 100μm. b) Cartoon illustrates the different metrics of invasion. c) Cartoons illustrate different types of invasive strand and the metrics that they would generate. d) Images show model output in an organotypic assay in the absence of fibroblasts, but with a single permissive track. e) Images illustrate how the Track Invasion Score is calculated from the simulations illustrated in panel d). Briefly, the extent of matrix remodelling in each ECM voxel is weighted based on the vertical distance from the starting cell – matrix interface. f) PCA plots showing the metrics derived from thousands of simulations in the absence of fibroblasts (grey-level plots) and both the presence and absence of fibroblasts (greenyellow plots) covering variation in cancer cell proteolysis, cancer cell – matrix adhesion, cancer cell – cancer cell adhesion.

**Supplementary Figure 3: Matrix proteolysis determines strand width, but not the distance invaded**

a) Heatmaps show how varying the cancer cell proteolysis value (x axis) impacts on different metrics in the absence of fibroblasts. WT indicates the ‘wild-type’ value based on experimental parameterisation using A431 cancer cells. b) Heatmaps show the differential in values resulting from the inclusion of fibroblasts (effectively a comparison of Figure 3b and Supp. Figure 3a). Red indicates an increase when fibroblasts are present, dark blue a reduction when in the presence of fibroblasts. c) Western blots of MMP14, CTNNA1, VIM, FN1, and ACTB in A431 cells engineered using Crispr/Cas9 to delete MMP14 or CTNNA1, or to over-express MMP14. d) Images show simulation output initiated with a spheroid, no fibroblasts, a uniform chemotactic cue and varying cancer cell proteolysis. Left panel · day 7output in absence of permissive track, right panel day 5 output in presence of permissive track.

**Supplementary Figure 4: Cancer cell – matrix adhesion modulates the tapering of strands**

a) Heatmaps show how varying the cancer cell – matrix adhesion value (x axis) impacts on different metrics in the absence of fibroblasts. WT indicates the ‘wild-type’ value based on experimental parameterisation using A431 cancer cells.

**Supplementary Figure 5: Cancer cell – cancer adhesion is required for efficient invasion in response to uniform directional cues**

a) Heatmaps show how varying the cancer cell – cancer cell adhesion value (x axis) impacts on different metrics in the absence of fibroblasts. WT indicates the ‘wild-type’ value based on experimental parameterisation using A431 cancer cells. b) Plots show how the mean number of neighbours of cancer cells vary as a function of invasion depth in simulations with varying cancer cell – cancer cell adhesion (c.f. Supp. Figure 2b). c) Plots show how the strand width varies depending on cancer cell – cancer cell adhesion in permissive track simulations.

**Supplementary Figure 6: Supra-cellular coordination of actomyosin organisation by cell – cell junctions enables wide invading strands**

a) Images show the F-actin (magenta) and myosin (MYH9/MHCIIa – green) networks in control A431 and 10μM Y27632 treated cells. Scale bar = 20μm. b) Images show the F-actin (magenta) and myosin (MYH9/MHCIIa – green) networks in control A431 ROCK:ER and 4-OHT-treated cells. Scale bar = 20μm.

**Supplementary Figure 7: Proteolysis-driven strand widening requires adherens junctions**

a) Images show simulations initiated with a spheroid, no fibroblasts, and a uniform downward chemotactic cue. Matrix proteolysis and cancer cell – cancer cell adhesion are varied. b) Heatmaps show how varying cancer cell – cancer cell adhesion value (x axis) and matrix proteolysis (y-axis) impacts on the curvature of the front of the invading cluster.

**Supplementary Figure 8: Strand widening is linked to tumour growth and metastasis**

a) Plot shows quantification of growth of A431 cells with the indicated manipulations of MMP14 and CTNNA1 in 2D cell culture. b) Phase contrast images show the growth of A431 ROCK:ER cancer cell colonies in the presence or absence of 4-OHT. Scale bar = 50μm. c) Plot shows quantification of the growth assay shown in b). Data from three biological replicates. d) Table relating patterns of collective invasion to cancer cell properties and the presence or absence of fibroblasts.

**Supplementary Table 1: Key CC3D parameter values**

Table of key parameter values used in Cellular Potts model and their real-world corresponding values.

**Supplementary Table 2: Additional CC3D parameter values**

Table of remaining parameter values used in Cellular Potts model.

**Supplementary Table 3: PCA loadings and variance explained**

Table of principal component loadings and variance explained for key simulation settings.

