## Supplementary figures and images for "Interplay of adherens junctions and matrix proteolysis determines the invasive pattern and growth of squamous cell carcinoma"

### Supplementary Figure 1

Supplementary Figure 1: Diversity of collective invasion in squamous cell carcinoma

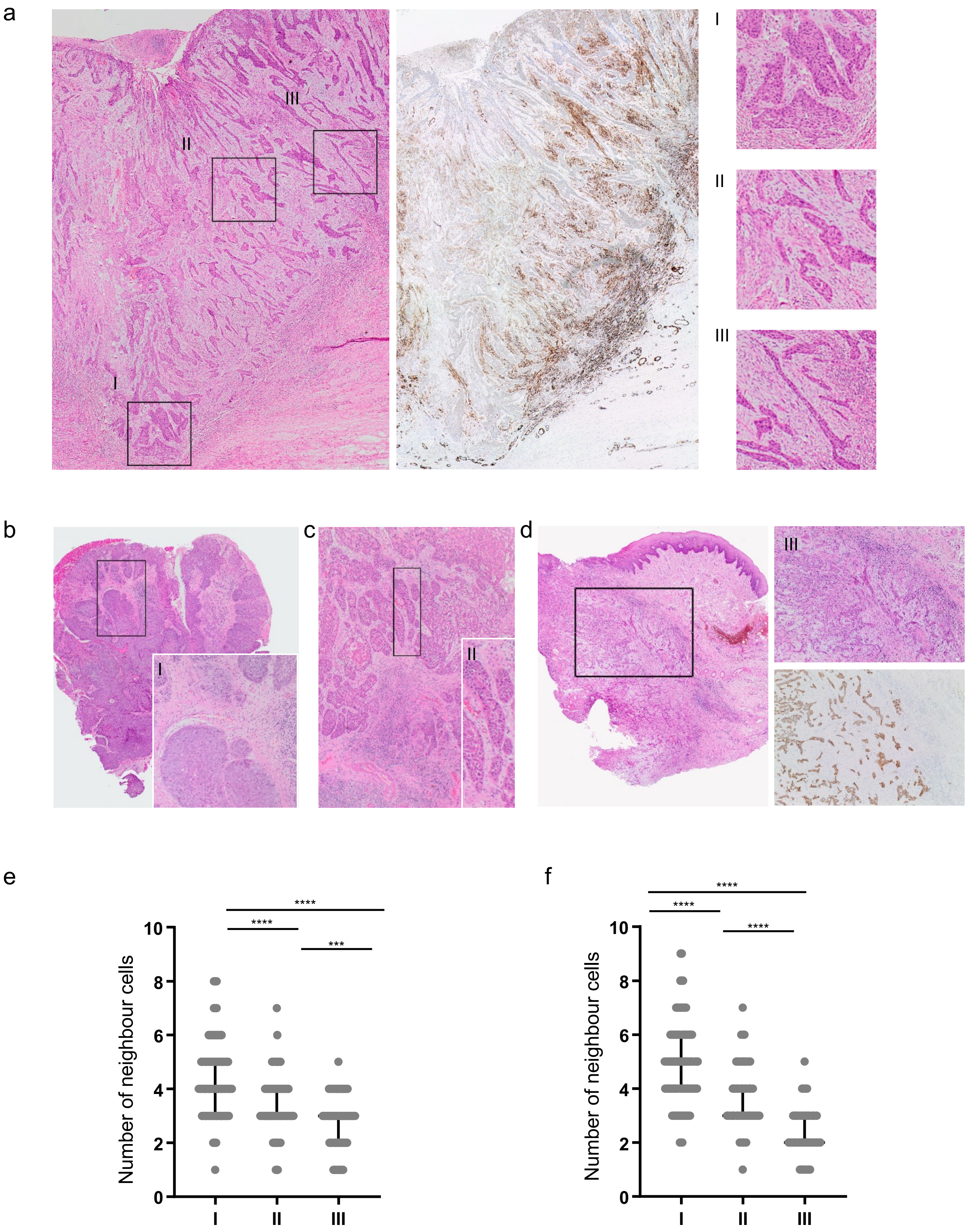

### Supplementary Figure 4

Supplementary Figure 4: Cancer cell – matrix adhesion modulates the tapering of strands

a

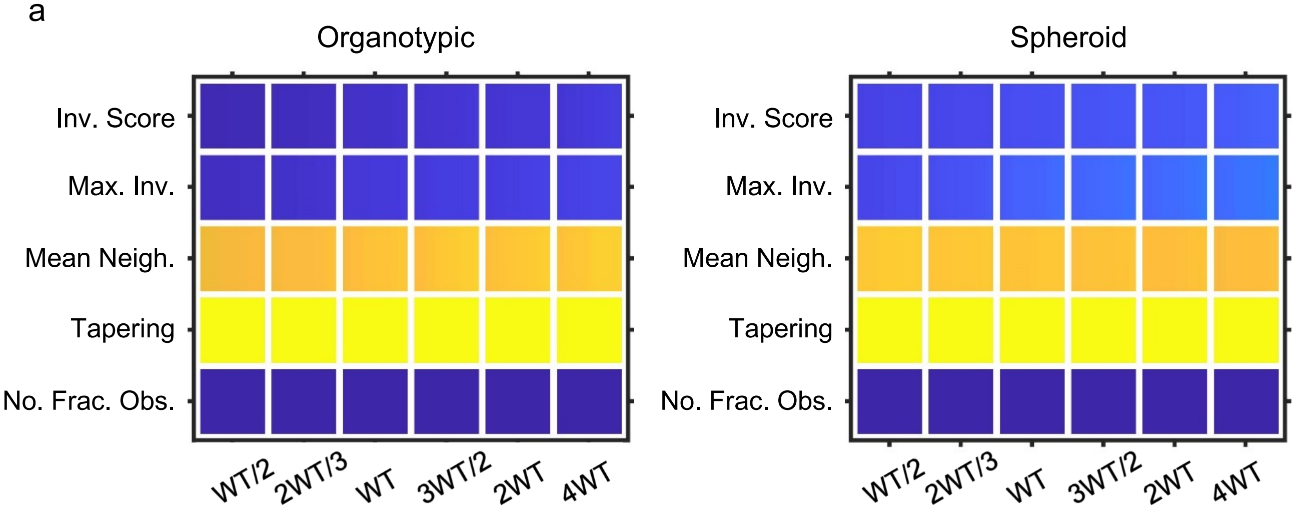

### Supplementary Figure 7

Supplementary Figure 7: Proteolysis-driven strand widening requires adherens junctions

a

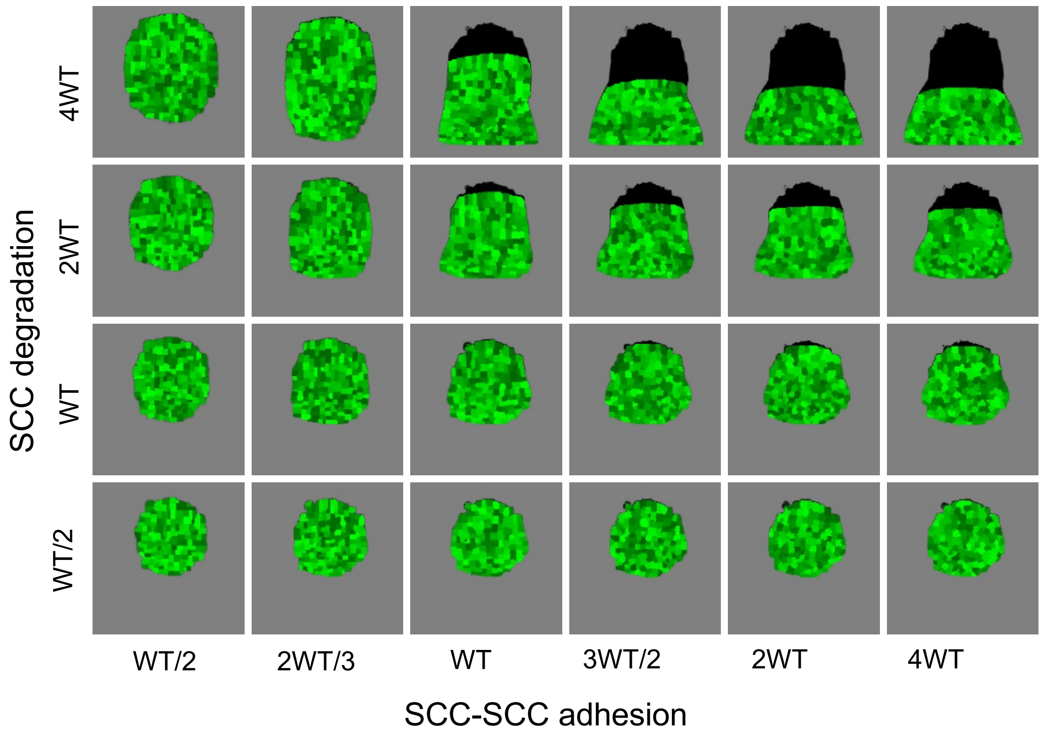


b

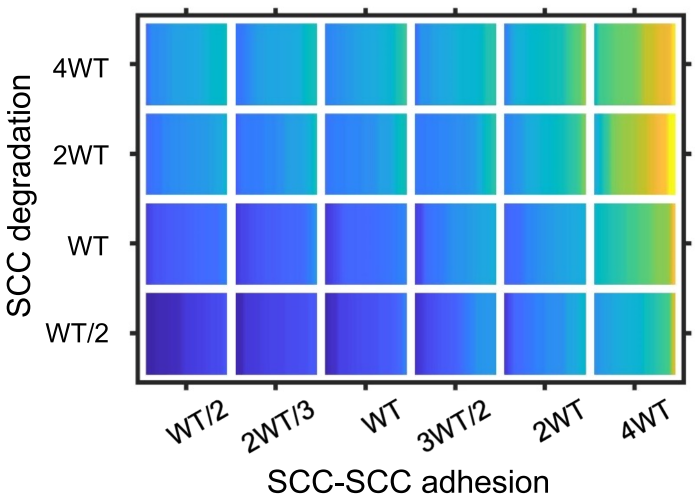
