## Supplementary Figure 2 for "Interplay of adherens junctions and matrix proteolysis determines the invasive pattern and growth of squamous cell carcinoma"

Supplementary Figure 2: Agent-based modelling recapitulates diversity of collective invasion pattern

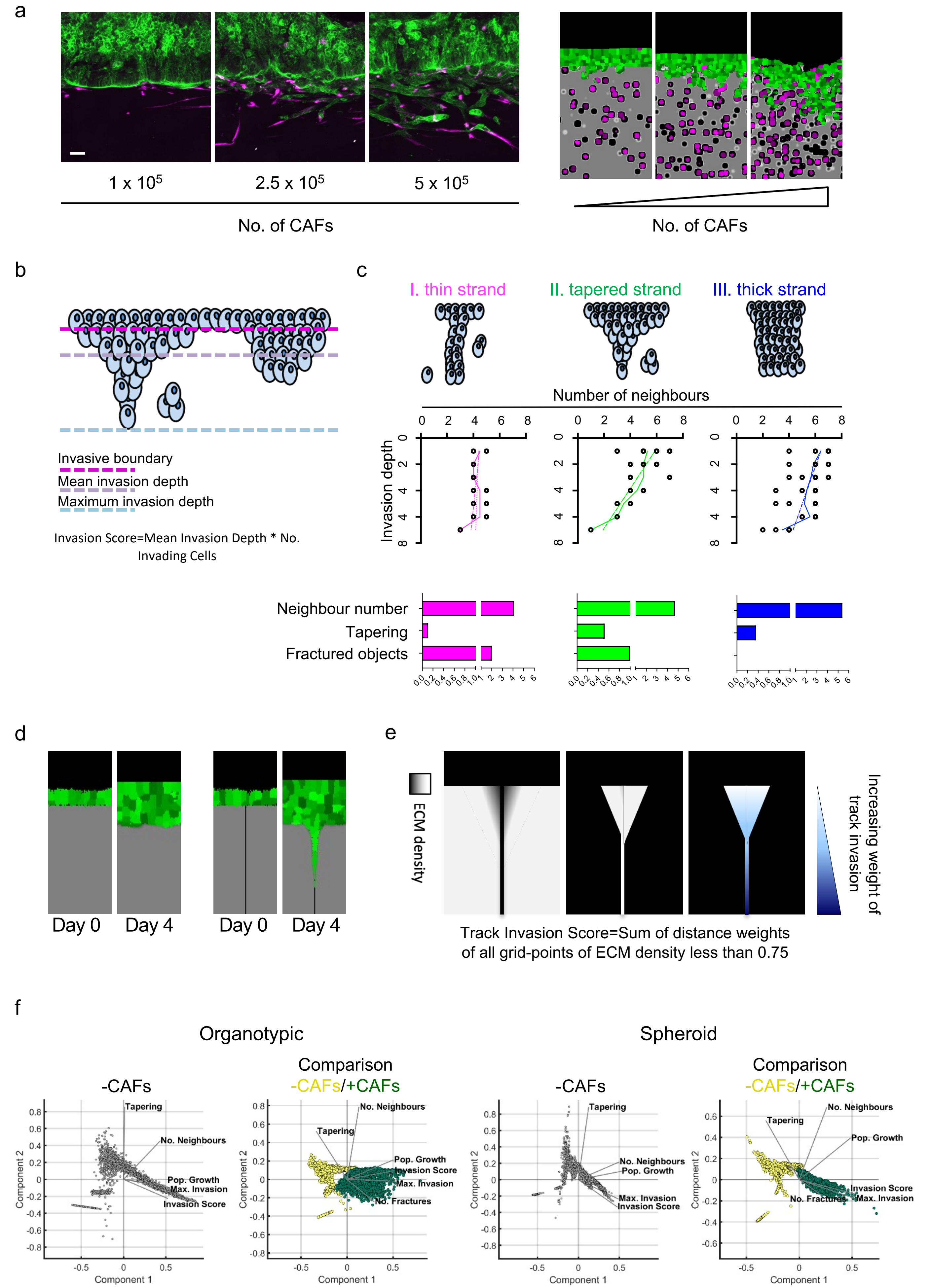
