## Supplementary Figure 3 for "Interplay of adherens junctions and matrix proteolysis determines the invasive pattern and growth of squamous cell carcinoma"

Supplementary Figure 3: Matrix proteolysis determines strand width, but not the distance invaded

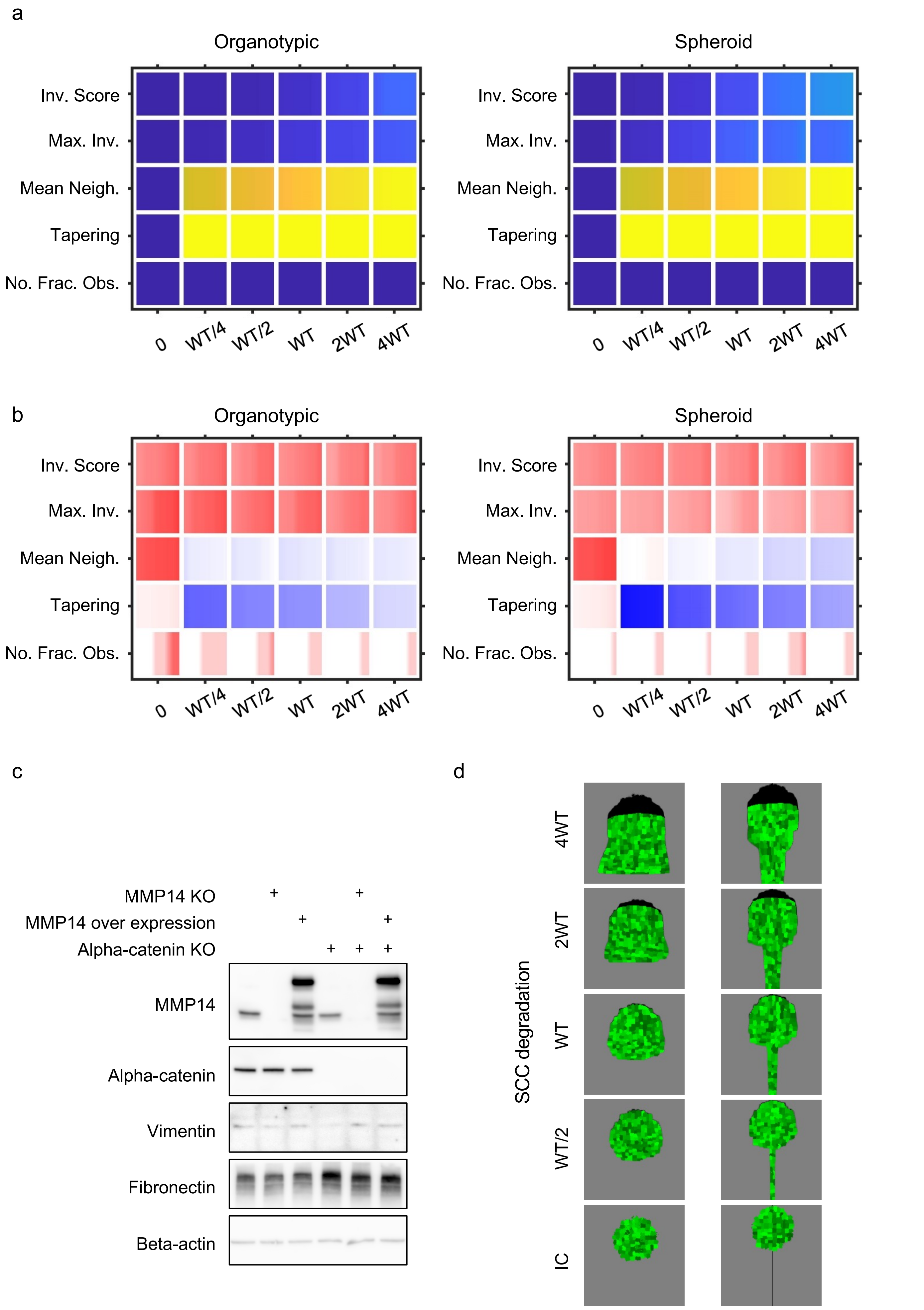
