## Supplementary Figure 5 for "Interplay of adherens junctions and matrix proteolysis determines the invasive pattern and growth of squamous cell carcinoma"

Supplementary Figure 5: Cancer cell – cancer adhesion is required for efficient invasion in response to uniform directional cues

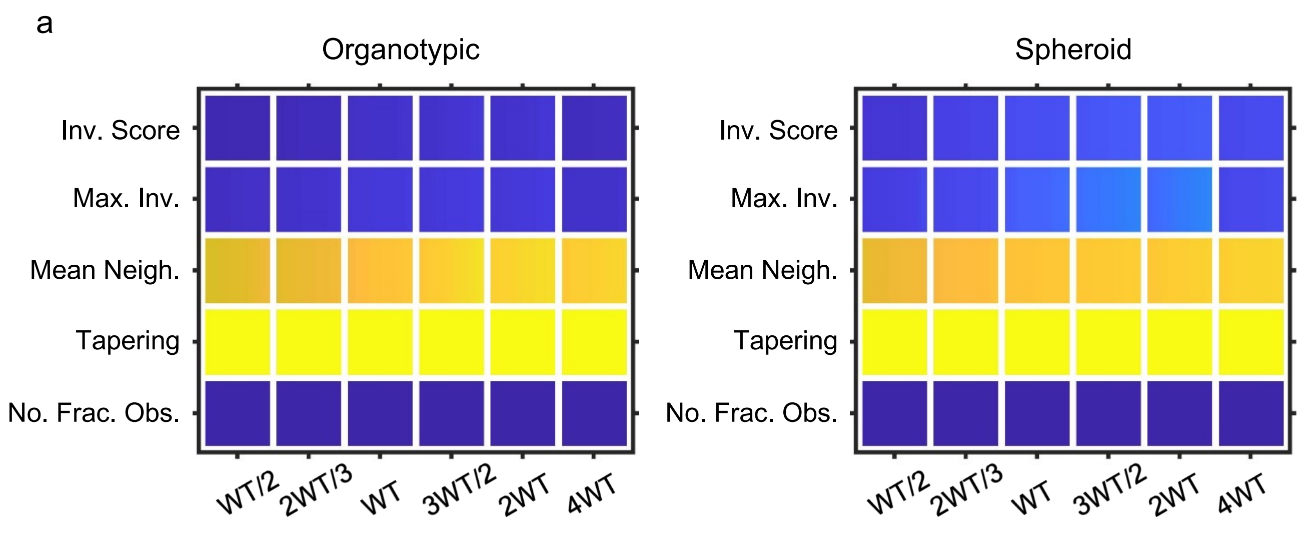


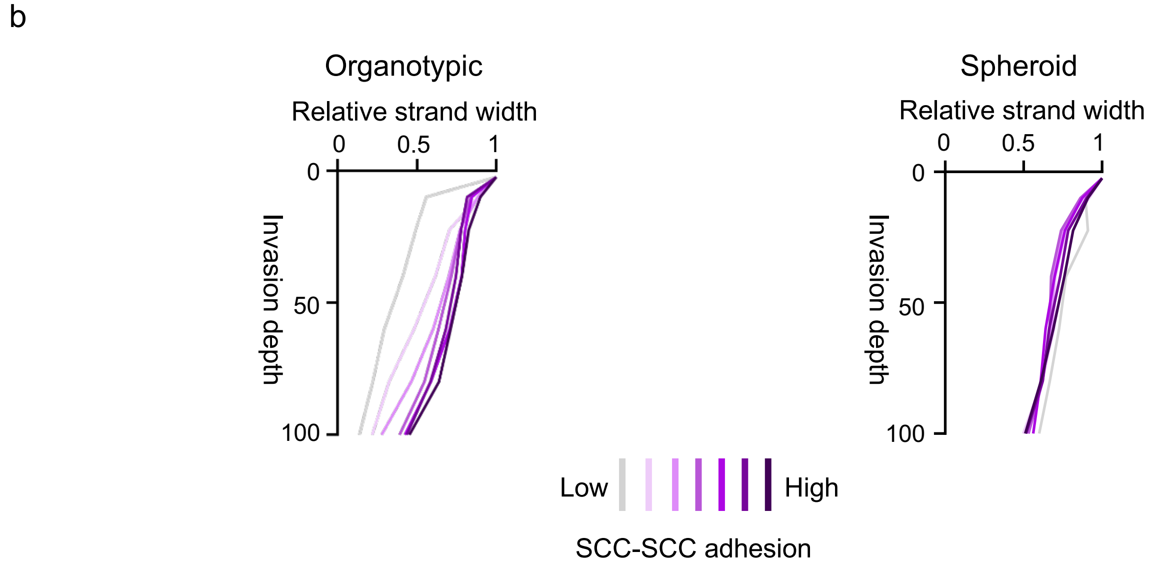


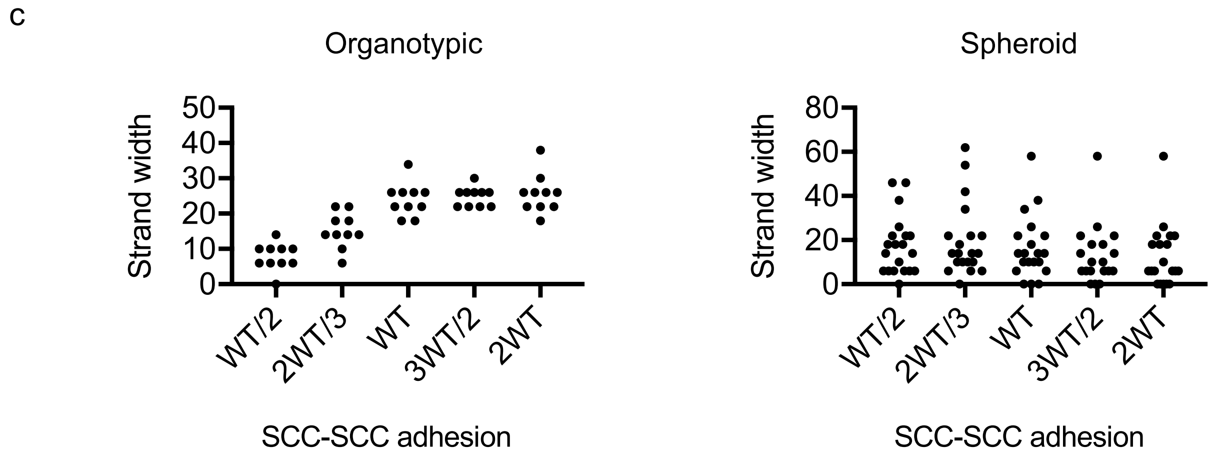
