## Supplementary Figure 6 for "Interplay of adherens junctions and matrix proteolysis determines the invasive pattern and growth of squamous cell carcinoma"

Supplementary Figure 6: Supra-cellular coordination of actomyosin organisation by cell – cell junctions enables wide invading strands

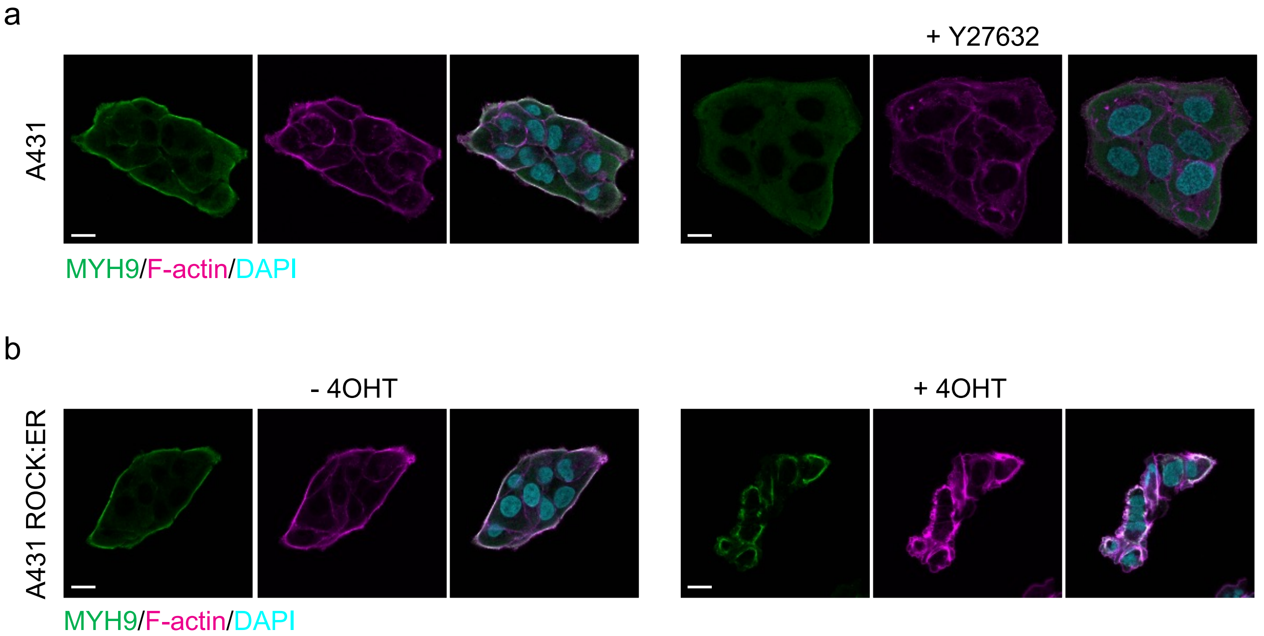
