## Supplementary Figure 8 for "Interplay of adherens junctions and matrix proteolysis determines the invasive pattern and growth of squamous cell carcinoma"

### Supplementary Figure 8: Strand widening is linked to tumour growth and metastasis

a

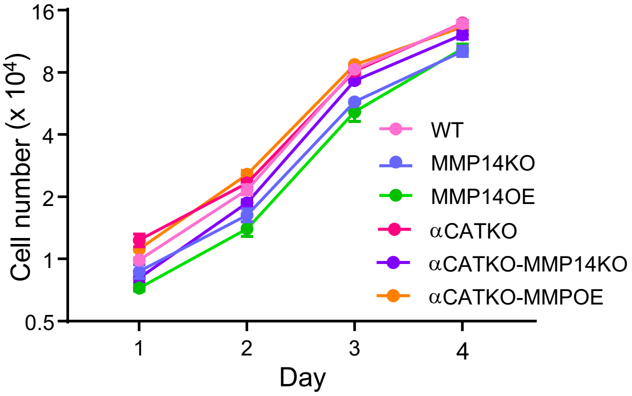


b

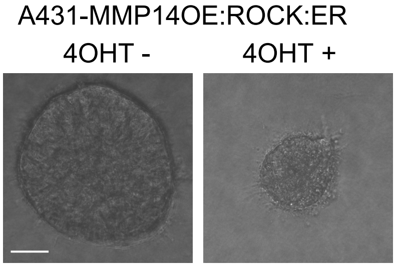


c

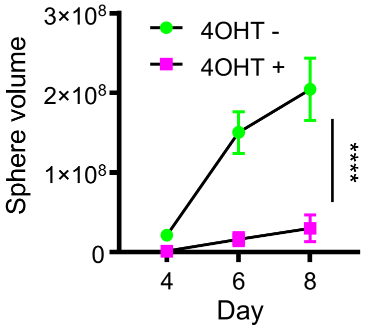


d

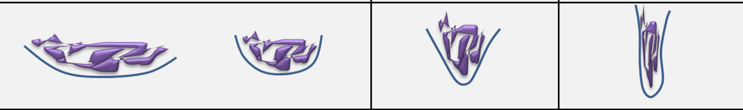

| | Pattern I | | Pattern II | Pattern III |
|---|---|---|---|---|
| Cell-cell adhesion | High | High | Medium | Low |
| Cell-matrix adhesion | < cell-cell adhesion | Intermediate | Low | > cell-cell adhesion |
| Matrix Proteolysis | High | High-medium | Low | Low |
| Directional cue | Uniform | Uniform | Radial | Radial or uniform |
| Fibroblasts | No | Maybe | Yes | Yes |
