## Supplementary Table 1 for "Interplay of adherens junctions and matrix proteolysis determines the invasive pattern and growth of squamous cell carcinoma"

### Supplementary Table 1: Key CC3D parameter values

| CC3D Parameter | CC3D Parameter value | Real world value | Comments |
|---|---|---|---|
| Monte Carlo timestep (MCS) | 1 | 30 seconds | |
| Voxel | 1 | 2 microns | |
| VCAF target volume | 800 voxels | 6400 microns$^3$ | 6500 microns$^3$ experimental measure |
| VCAF target surface | 700 voxels | 2800 microns$^2$ | 4900 microns$^2$ experimental measure |
| SCC initial target volume | 400 voxels | 3200 microns$^3$ | |
| SCC dividing volume | 800 voxels | 6400 microns$^3$ | |
| Median SCC volume in WT conditions | 550 voxels | 4400 microns$^3$ | 4500 microns$^3$ experimental measure |
| SCC surface area | 324 voxels (median) | 1296 microns$^2$ (median) | Sphere assumed for surface area<br>1700 microns$^2$ experimental measure |
| Mean time to mitosis | 8640 MCS | 3 days | |
| SCC-ECM adhesion | 10 (contact energy) | 45.53 (experimental measure) | Adhesions are normalised to SCC-ECM adhesion. They are inverted and multiplied by 10 to give contact energies. |
| SCC-SCC adhesion | 21 (contact energy) | 21.8 | |
| SCC-CAF adhesion | 35 | 25.2 | SCC-CAF adhesion was marginally reduced below experimental measure in model (contact energy would be 18 from experiment) |
| CAF-CAF adhesion | 45 | 9.3 | |
| CAF-ECM adhesion | 15 | 29.6 | |
| SCC-ECM adhesion for zero density ECM | 40 | 11.4 | |
| CAF-CAF repulsion range | 20 voxels | 40 microns | Approximately 1.5 CAF widths |
| SCC taxis energy | 13 | median speed 0.2 microns/min | 0.2 microns/min experimental measure |
| CAF taxis minimum energy | 3.5 | 0.06 micron/min | |
| CAF taxis maximum energy | 21 | 0.29 microns/min | |
| CAF median speed | 10 | 0.1 microns/min | 0.1 microns/min experimental measure |
| CAF taxis stimulation range | 30 voxels | | Approximately 2.5 CAF widths |
| CAF ECM pushing rate | 0.0140 | Corresponds to a reduced CAF speed of 0.07 microns/min through ECM | Speed due to pushing is 3 times faster than speed due to degradation. |
| CAF ECM degradation rate | 0.0012 | Corresponds to a reduced CAF speed of 0.018 microns/min through ECM | Degradation and pushing effects on speed are sub-linear. The effective median speed is between 0.07 and 0.088 microns/min. |
| SCC pushing rate | 0 | | |
| SCC degradation rate | 0.0001 | Corresponds to a reduced SCC speed of 0.009 microns/min through ECM | Effect of degradation is half for SCCs compared to CAFs. Effects are normalised to kinesis levels. |
