## Supplementary Table 2 for "Interplay of adherens junctions and matrix proteolysis determines the invasive pattern and growth of squamous cell carcinoma"

### Supplementary Table 2: Additional CC3D parameter values

| CC3D Parameter | CC3D Parameter value |
|---|---|
| **Organotypic Directionality Weights (selected such that invasive pattern mimics experimental data)** | |
| SCC random direction | 0.5 |
| SCC chemotactic direction | 0.5 |
| CAF random direction | 0.3 |
| CAF protrusion direction | 0.3 |
| CAF persistence direction | 0.4 |
| **Spheroid Directionality Weights** | |
| SCC random direction | 0.5 |
| SCC radial direction | 0.5 |
| CAF random direction | 0.1 |
| CAF protrusion direction | 0.1 |
| CAF radial direction | 0.8 |
| **Energy parameters** | |
| SCC volume energy | 1 |
| SCC surface area | 0.01 |
| CAF volume energy | 1 |
| CAF surface area | 0.25 |
| Erlang shape parameter for SCC division | 6 |
| CAF repulsion energy | 100 |
| CAF kinesis stimulation (CAF stimulation from neighbouring cells) | 0.09 |
| CAF kinesis decay rate | 1 |
| Cell ECM penetration penalty energy (Magnitude of penalty for moving into ECM of density 1. Penalty changes linearly with ECM density) | 100 |
| ECM density penalty threshold (Cells with no remodelling ability suffer a very large energy penalty for moving into ECM above this threshold density) | 0.9 |
| CAF remodelling minimum energy (Remodelling rate of CAFs determined by current kinesis levels. Minimum energy of zero means any CAF that is motile will remodel the ECM.) | 0 |
| CAF remodelling maximum energy | 21 |
| SCC remodelling CAF equivalent energy (CAF kinesis level required to match SCC motility. Normalises cell type remodelling effects) | 17 |
| Cell ECM adhesion factor (loss of adhesion and ECM density reduces) | 0.25 |
