## Supplementary Table 3 for "Interplay of adherens junctions and matrix proteolysis determines the invasive pattern and growth of squamous cell carcinoma"

### Supplementary Table 3: PCA loadings and variance explained

| **Organotypic + CAFs** | | | | | | |
|---|---|---|---|---|---|---|
| **Loadings** | **PC1** | **PC2** | | | **Var. Explained** | **Cum. Var. Explained** |
| **Maximum Invasion** | 0.48965 | -0.3021 | | **PC1** | 47.2081 | 47.2081 |
| **Invasion Score** | 0.5542 | 0.043342 | | **PC2** | 27.4539 | 74.6619 |
| **Number Neighbours** | 0.28187 | 0.61759 | | **PC3** | 14.1015 | 88.7634 |
| **Tapering** | -0.36162 | 0.455 | | **PC4** | 5.1516 | 93.915 |
| **Fractured Objects** | 0.030328 | -0.49911 | | **PC5** | 4.0917 | 98.0067 |
| **Cell Growth** | 0.49191 | 0.26326 | | **PC6** | 1.9933 | 100 |

| **Organotypic -CAFs** | | | | | | |
|---|---|---|---|---|---|---|
| **Loadings** | **PC1** | **PC2** | | | **Var. Explained** | **Cum. Var. Explained** |
| **Maximum Invasion** | 0.54135 | 0.096364 | | **PC1** | 64.0988 | 64.0988 |
| **Invasion Score** | 0.52256 | 0.22653 | | **PC2** | 25.8079 | 89.9067 |
| **Number Neighbours** | 0.43157 | -0.45305 | | **PC3** | 5.5076 | 95.4143 |
| **Tapering** | 0.01147 | -0.85397 | | **PC4** | 4.1319 | 99.5462 |
| **Cell Growth** | 0.49748 | 0.069901 | | **PC5** | 0.4538 | 100 |

| **Organotypic + CAFs -CAFs** | | | | | | |
|---|---|---|---|---|---|---|
| **Loadings** | **PC1** | **PC2** | | | **Var. Explained** | **Cum. Var. Explained** |
| **Maximum Invasion** | 0.52676 | -0.018388 | | **PC1** | 54.9694 | 54.9694 |
| **Invasion Score** | 0.50918 | 0.094024 | | **PC2** | 22.656 | 77.6254 |
| **Number Neighbours** | 0.12876 | 0.78576 | | **PC3** | 13.1608 | 90.7862 |
| **Tapering** | -0.33366 | 0.52 | | **PC4** | 5.2631 | 96.0493 |
| **Fractured Objects** | 0.29201 | -0.2427 | | **PC5** | 2.4501 | 98.4994 |
| **Cell Growth** | 0.50009 | 0.20999 | | **PC6** | 1.5006 | 100 |

| **Spheroid + CAFs** | | | | | | |
|---|---|---|---|---|---|---|
| **Loadings** | **PC1** | **PC2** | | | **Var. Explained** | **Cum. Var. Explained** |
| **Maximum Invasion** | 0.57566 | 0.15072 | | **PC1** | 42.8451 | 42.8451 |
| **Invasion Score** | 0.44128 | 0.48587 | | **PC2** | 22.3283 | 65.1734 |
| **Number Neighbours** | -0.36688 | 0.52766 | | **PC3** | 16.6975 | 81.8709 |
| **Tapering** | -0.56428 | 0.14708 | | **PC4** | 12.2754 | 94.1463 |
| **Fractured Objects** | 0.027087 | 0.59877 | | **PC5** | 3.202 | 97.3484 |
| **Cell Growth** | -0.14193 | 0.28745 | | **PC6** | 2.6516 | 100 |

| **Spheroid -CAFs** | | | | | | |
|---|---|---|---|---|---|---|
| **Loadings** | **PC1** | **PC2** | | | **Var. Explained** | **Cum. Var. Explained** |
| **Maximum Invasion** | 0.48621 | 0.23444 | | **PC1** | 70.1278 | 70.1278 |
| **Invasion Score** | 0.47407 | 0.33209 | | **PC2** | 23.3452 | 93.4729 |
| **Number Neighbours** | 0.49779 | -0.21567 | | **PC3** | 3.7844 | 97.2574 |
| **Tapering** | 0.12511 | -0.88191 | | **PC4** | 2.5016 | 99.7589 |
| **Cell Growth** | 0.5248 | -0.10238 | | **PC5** | 0.2411 | 100 |

| **Spheroid + CAFs -CAFs** | | | | | | |
|---|---|---|---|---|---|---|
| **Loadings** | **PC1** | **PC2** | | | **Var. Explained** | **Cum. Var. Explained** |
| **Maximum Invasion** | 0.53047 | 0.17858 | | **PC1** | 51.2063 | 51.2063 |
| **Invasion Score** | 0.53239 | 0.13651 | | **PC2** | 26.3236 | 77.5299 |
| **Number Neighbours** | 0.26057 | -0.68358 | | **PC3** | 15.4308 | 92.9607 |
| **Tapering** | -0.31488 | -0.55834 | | **PC4** | 4.8707 | 97.8314 |
| **Fractured Objects** | 0.18666 | 0.1297 | | **PC5** | 1.7236 | 99.555 |
| **Cell Growth** | 0.48299 | -0.39195 | | **PC6** | 0.445 | 100 |
