## Supplementary CC3D Code Walkthrough for "Interplay of adherens junctions and matrix proteolysis determines the invasive pattern and growth of squamous cell carcinoma"

### Interplay of adherens junctions and matrix proteolysis determines the invasive pattern and growth of squamous cell carcinoma: A walkthrough of developed CC3D plug-ins and steppables

Robert P. Jenkins* and Melda Tozluoglu

#### 1 Cellular Potts Model Documentation

##### 1.1 Biology of the Setup

The system is composed of two cell types, SCC cells and VCAFs. Two types of assay are simulated - the organotypic invasion assay and the spheroid assay. The extracellular matrix (ECM) surrounding the cells prevents cell movement where the ECM is dense. Matrix density surrounding a cell also determines a cell's adhesion to ECM. Both cell types can remodel the ECM to generate tracks. This remodelling involves degradation and dislocation of ECM, but not secretion. VCAFs can both degrade and dislocate ECM. SCCs have only a low level ability to degrade the matrix. VCAFs are stimulated to move and remodel the ECM by nearby cells. VCAFs also repulse neighbouring VCAFs. SCC cells have a chemotactic response, driving metastasis. This chemotactic response is uniform in the organotypic context and radial in the spheroid context. Assays are typically simulated for five days. SCC cells proliferate with a frequency of 0.33 cell divisions per day in the simulations. VCAFs do not divide during the simulations, posessing an average cell division time in excess of one week - greater than the timeframe of simulation.

##### 1.2 Implemented Functions Overview

The Cellular Potts Model (CPM) is used to model the system and CompuCell3D software is utilised for implementation. The existing functionality of the Compucell3D software is used where available. Pixel tracker plugin and Boundary Pixel tracker plugins allow the storing of pixels and boundary pixels respectively, belonging to a given cell. The Center Of Mass plugin tracks and updates the centre of mass of each cell. VolumeLocalFlex and SurfaceLocalFlex allow the setting of

*.

volume and surface area to each cell individually. This is useful for growth during cell division. Similary, AdhesionFlex allows the setting of each cell's adhesive properties individually. This is particularly useful when the density of ECM surrounding a cell determines its cell-ECM adhesion. The Neighbor tracker plugin tracks the neighbours of every cell and the size of overlap between each neighbour. It is used to generate output data for analysis. Since our simulation space is large, Global Boundary Pixel tracker tracks the boundary pixels of every cell such that only pixel copy attempts on cell boundaries are considered. It is used in conjunction with the Boundary Walker algorithm. Volume tracker is used to track changes in cellular volume. It is used frequently when determining the centroid of each cell and when evaluating energy contributions of pixel copy attempts. Multiple functions are implemented to the software to develop a model including specific needs of the biological phenomena of interest. In terms of cell motility, these functions include cell directionality and VCAF kinesis stimulation. In terms of ECM interaction they include ECM remodelling and ECM density dependent adhesion. Finally, cell division of SCCs is also included.

#### 1.3 CPM Flow and Nomenclature

In the Cellular Potts Model, the modelling environment is represented on a grid. Each point of the grid can be occupied by the medium, or a cell of given type. Each cell in the model is represented by a collection of grid points. At each step of the simulation, all grid points try to copy themselves to a neighbouring grid point. In biological terms, if grid point $(i, j, k)$ belongs to an SCC cell, and the grid point succeeds in copying itself to grid point $(i+1, j, k)$, it would be imitating the SCC cell changing morphology and protruding into the direction of new grid point - grid point $(i+1, j, k)$. Similarly, if grid point $(i+2, j, k)$ belongs to medium, and $(i+1, j, k)$ belongs to an SCC cell, the same copy event would mean SCC cell retracting a part of its body (Figure 1). Throughout this document, at each copy attempt the cell attempting to copy into a given pixel will be known as newCell and the cell currently occupying that pixel prior to the copy attempt will be known as oldCell.

The success of the copy attempt is probabilistic, dependent on the energy change in the system associated with the grid copy. Each of the biological phenomena described in Subsection 1.1 are represented by an energy contribution. Calculation steps for each one of the defined energy contributions are called 'plugins' in CompuCell3D nomenclature. If the overall energy change upon grid point copy is negative, the copy attempt will be accepted, and if the energy change is positive, the change will be accepted with a Boltzmann-weighted probability

$$P(\Delta E) = \exp(-\Delta E/\beta). \tag{1.1}$$

Here, probability $P(\Delta E)$ is inversely proportional to the change in energy, and scaled by the fluctuation amplitude of the cell, $\beta$. One time step, or Monte Carlo step (MCS), is defined as running through all grid points of the system in a random order, and making grid copy attempts. Once one time step is complete, updates in overall system and cell properties are made, such as remodelling of the ECM, or

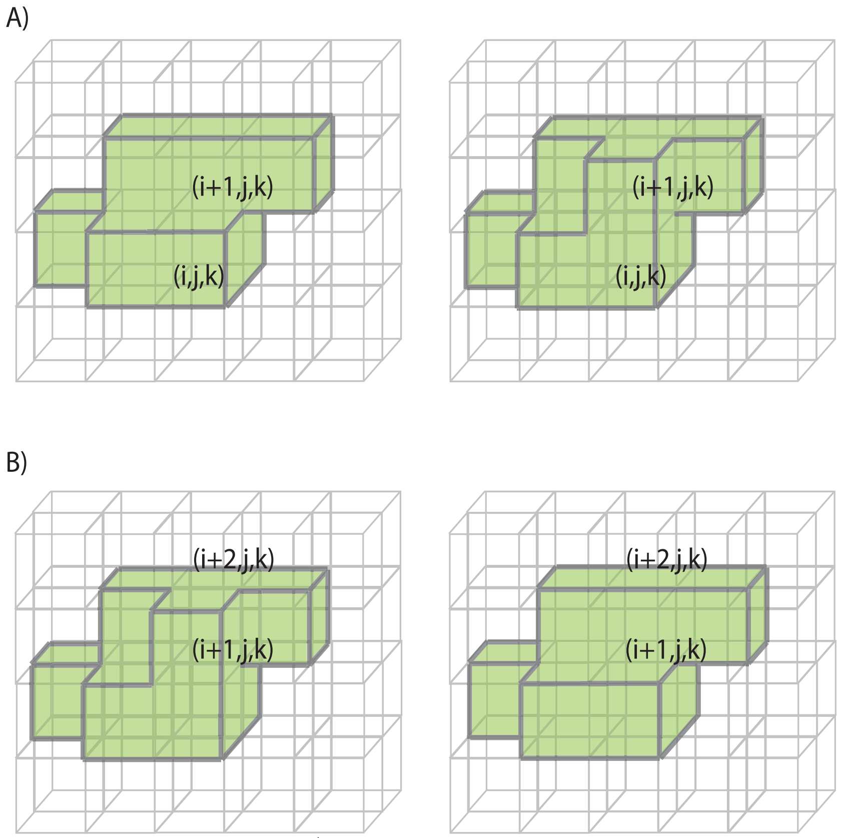


**Figure 1:** Grid copy events A) An SCC cell protruding into the medium. B) An SCC cell retracting part of its body.

a cell division. The functions executing such updates between each time step are called 'steppables' in CC3D (Figure 2). Each plugin and steppable is introduced in turn, both with the theory and the C++ code. Some plugins and steppables work in both the organotypic and spheroid contexts. In other cases a separate plugin or steppable has been written for each environment. These differences generally arise either because the organotypic environment requires periodic boundary conditions and the spheroid environment does not or that the directional cues are different in each case. In instances where the plugin or steppable differs between the two environments, the theory and code are first introduced for the organotypic context and then condensed versions are introduced for the spheroid context.

### 2 SCC cell division

#### 2.1 Methodology

Cell division is implemented identically in both the organotypic and spheroid contexts. The time period to division for each cell is a random variable, either from the exponential distribution or Erlang distribution at the discretion of the user. The probability density function (pdf) of the exponential distribution is

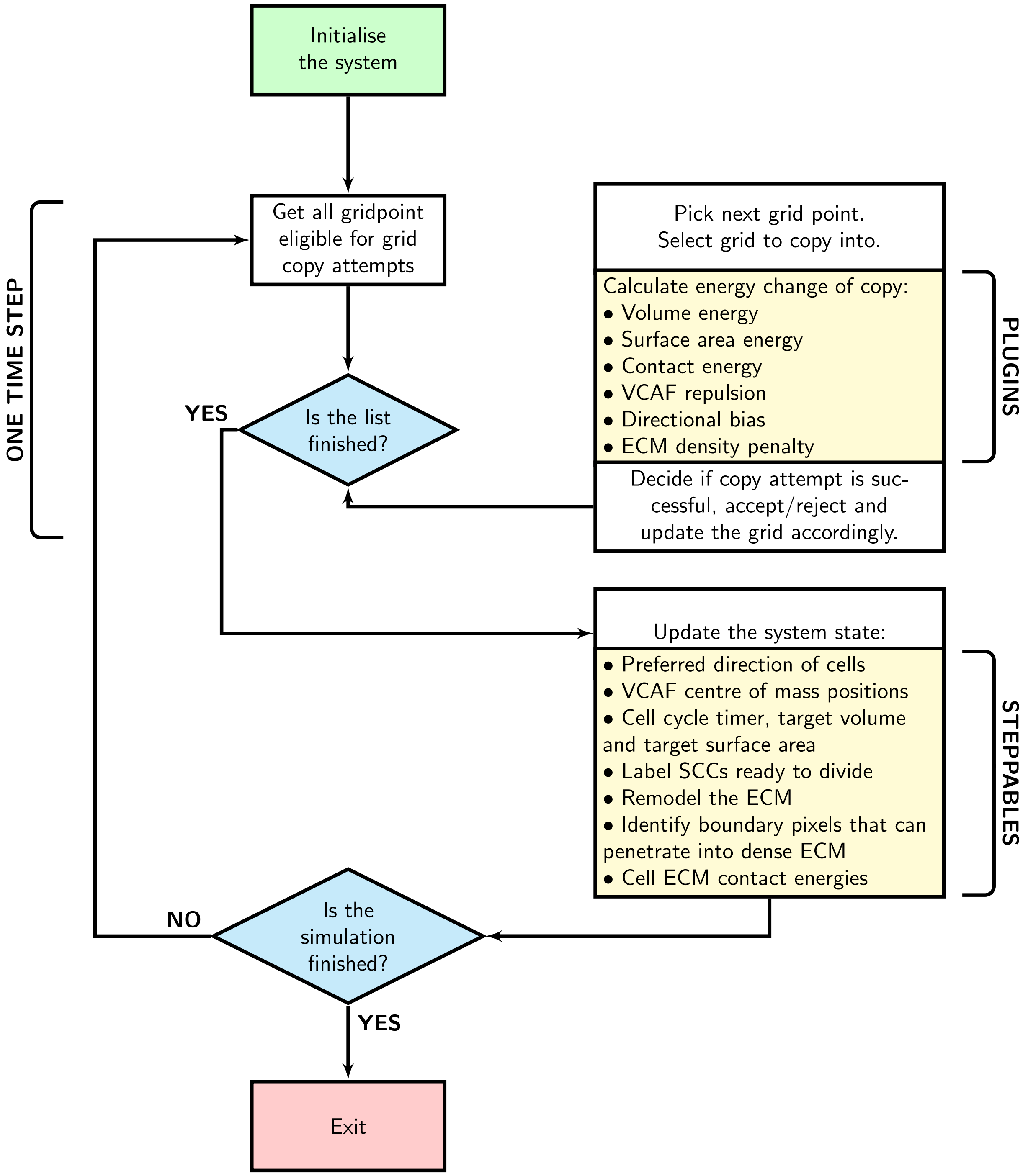


**Figure 2:** A simple flow diagram defining the flow of Cellular Potts Model implementation in Compucell3D. The plugins calculate the energy contribution of each of the defined biological phenomena to the current grid point copy attempt. After all grid points are checked for copy attempts, the system's properties are updated in steppables. Execution of all listed steppables completes one time step. This procedure is repeated until the desired number of time steps are simulated.

given by

$$p(x) = \begin{cases} \lambda e^{-\lambda x} & x \geq 0, \\ 0 & x < 0, \end{cases} \tag{2.1}$$

with mean $1/\lambda$. For a uniformly distributed random variable, $U \sim \mathcal{U}(0,1)$, an exponential random variable, $X$, can be derived

$$X = -\frac{1}{\lambda}\log(U). \tag{2.2}$$

The pdf of the Erlang distribution is given by

$$p(x) = \frac{\lambda^k x^{k-1} e^{-\lambda x}}{(k-1)!},\ x, \lambda \geq 0 \tag{2.3}$$

where $k$ is the shape parameter and $\lambda$ is the rate parameter. The mean is given by $k/\lambda$. For $k$ uniformly distributed random variables, $U_i \sim \mathcal{U}(0,1)$, an Erlang random variable, $X$, can be derived

$$X = -\frac{1}{\lambda}\log\left(\prod_{i=1}^{k} U_i\right). \tag{2.4}$$

Following this allocation of random time for cell division, the cell attempts to increase its volume linearly with time. Upon reaching a volume threshold the cell divides with a new random time for cell division being allocated to each of the daughter cells independently. The increase in volume over time could be hindered by other environmental factors such as competition for space, causing the cell to take longer to divide than the period allocated, or effectively enter a mitotically quiescent state where it cannot grow large enough to divide. Target cell volume increases linearly with time but the manner in which surface area increases can be selected by the user, either to always increase by the same ratio as the volume or to increase at a rate comparable to spheres i.e. for a cell with target volume, $V$, its target surface area, $SA$, can be calculated

$$r = \left(\frac{3V}{4\pi}\right)^{1/3}, \tag{2.5}$$

$$SA = 4\pi r^2. \tag{2.6}$$

Cell division is implemented in two separate steppables. The main steppable for use in organotypic and spheroid simulations is described in Subsubsection 2.2.2. These simulations however, require initial condition files. These initial condition files include the pif files, with the pixel locations of all cells and also the stage of cell division of every cell. The initial condition cells should all be at different stages of their cell cycle and thus they should all be different sizes. In addition, due to the random nature of cell division, the time to divide should be an independent random variable for each cell. The cell division steppable required in the generation of these initial conditions is described first, in Subsubsection 2.2.1.

##### 2.1.1 Cell division initial condition generation

In order to set up these random initial condition files for organotypic and spheroid simulations we first need to run a set of simulations to generate them. In the case of organotypic generators, SCCs are initialised resting on top of ECM, with a chemotactic response downwards and zero ECM remodelling ability. This causes cells to push downwards but they are unable to move into the ECM, keeping them localised on top of the invasive boundary. In the case of spheroids, SCCs can be set up as a sphere of cells, with no directionality imposed and in medium rather than ECM. This combination stops cells from moving radially outwards but allows the mass to grow in volume outwards. In both the organotypic and spheroid cases, fibroblasts can be initialised with the SCCs. In the initialisations, fibroblasts have zero remodelling ability and their number remains constant as they possess no ability to divide.

In either assay, SCCs are input into these initial condition simulations with identical volumes and a uniformly distributed random initial delay until they enter the cell cycle. Hence they all enter the cell cycle at a random MCS and then grow in size according to their own cell division clock. Thus, at the end of these initial condition generating simulations the cells are well mixed and of different size and cell cycle stage. By running multiple initialising simulations, we can generate initial condition files with different distributions of cells both spatially and in terms of their cell cycle stage.

#### 2.2 Code Documentation

##### 2.2.1 Cell Division Initialisation Steppable

We begin with the steppable, MitosisInitialisation, required to generate cell division initial condition files. The functions below are given in the MitosisInitialisation.cpp file. The steppable begins with an initiation function, Listing 2.2.1, that identifies and links the simulation main frame to the steppable. Plugins utilised by the steppable are loaded in this initialisation, in this case a pixel tracker plugin.

```
void MitosisInitialisation::init(Simulator *simulator, CC3DXMLElement *
    _xmlData) {
    potts = simulator->getPotts();
    sim=simulator;
    cellFieldG = (WatchableField3D<CellG *> *)potts->getCellFieldG();
    simulator->registerSteerableObject(this);
    update(_xmlData);
    bool pluginAlreadyRegisteredFlag;
    pixelTrackerPlugin=(PixelTrackerPlugin*)Simulator::pluginManager.get("
        PixelTracker",&pluginAlreadyRegisteredFlag);
    if(!pluginAlreadyRegisteredFlag)
        pixelTrackerPlugin->init(simulator);
    pixelTrackerAccessorPtr=pixelTrackerPlugin->getPixelTrackerAccessorPtr();
}
```

**Listing 2.2.1:** SCC cell division initialisation Steppable - init

The update function, Listing 2.2.2, reads the simulation inputs from the xml simulation setup and allocates them to the steppable. This includes VCAF and SCC cell types. Cell volume, volume energy and equivalents for surface area are included with default parameter fits set to experimental data. The mean time that each cell must wait prior to entering the cell division cycle is included as are input parameters relating to the cell division cycle including a multiplier to determine the size the cell grows to over the cell cycle and the size threshold at which the cell then divides. Input for how the increase in surface area occurs (at the same rate as volume or such that the cell maintains a spherical shape) also exists. The probability density used to determine length of time a cell will take to divide (exponential or Erlang) in addition to the mean time to division and the shape parameter (in the case of Erlang) are included. Finally, output text file names and the MCS where the initialisation records all of the output needed for initial conditions of future simulations are included. Many of the inputs include default values in the absence of an xml input including the default values for cell type being set to 1 for SCC and 2 for VCAF.

```
void MitosisInitialisation::update(CC3DXMLElement *_xmlData, bool _fullInitFlag){
    if(_xmlData->findElement("SCCtype")){SCCtype=_xmlData->getFirstElement("SCCtype")->getDouble();}
    else{SCCtype=1;cerr<<"SCCtype not specified, defaul t value set to: "<<SCCtype<<endl;}
    if(_xmlData->findElement("VCAFtype")){VCAFtype=_xmlData->getFirstElement("VCAFtype")->getDouble();}
    else{VCAFtype=2;cerr<<"VCAFtype not specified, defaul t value set to: "<<VCAFtype<<endl;}
    if(_xmlData->findElement("SCCVolume")){SCCVolume=_xmlData->getFirstElement("SCCVolume")->getDouble();}
    else{SCCVolume=550;cerr<<"SCCVolume not specified, defaul t value set to: "<<SCCVolume<<endl;}
    if(_xmlData->findElement("VCAFVolume")){VCAFVolume=_xmlData->getFirstElement("VCAFVolume")->getDouble();}
    else{VCAFVolume=800;cerr<<"VCAFVolume not specified, defaul t value set to: "<<VCAFVolume<<endl;}
    if(_xmlData->findElement("SCCLambdaVolume")){SCCLambdaVolume=_xmlData->getFirstElement("SCCLambdaVolume")->getDouble();}
    else{SCCLambdaVolume=1;cerr<<"SCCLambdaVolume not specified, defaul t value set to: "<<SCCLambdaVolume<<endl;}
    if(_xmlData->findElement("VCAFLambdaVolume")){VCAFLambdaVolume=_xmlData->getFirstElement("VCAFLambdaVolume")->getDouble();}
    else{VCAFLambdaVolume=1;cerr<<"VCAFLambdaVolume not specified, defaul t value set to: "<<VCAFLambdaVolume<<endl;}
    if(_xmlData->findElement("SCCSurface")){SCCSurface=_xmlData->getFirstElement("SCCSurface")->getDouble();}
    else{SCCSurface=550;cerr<<"SCCSurface not specified, defaul t value set to: "<<SCCSurface<<endl;}
    if(_xmlData->findElement("VCAFSurface")){VCAFSurface=_xmlData->getFirstElement("VCAFSurface")->getDouble();}
    else{VCAFSurface=700;cerr<<"VCAFSurface not specified, defaul t value set to: "<<VCAFSurface<<endl;}
    if(_xmlData->findElement("SCCLambdaSurface")){SCCLambdaSurface=_xmlData->getFirstElement("SCCLambdaSurface")->getDouble();}
    else{SCCLambdaSurface=1;cerr<<"SCCLambdaSurface not specified, defaul t value set to: "<<SCCLambdaSurface<<endl;}
    if(_xmlData->findElement("VCAFLambdaSurface")){VCAFLambdaSurface=_xmlData->getFirstElement("VCAFLambdaSurface")->getDouble();}
    else{VCAFLambdaSurface=1;cerr<<"VCAFLambdaSurface not specified, defaul t value set to: "<<VCAFLambdaSurface<<endl;}
    if(_xmlData->findElement("InitialTimeToMitosisSCC")){InitialTimeToMitosisSCC=_xmlData->getFirstElement("InitialTimeToMitosisSCC")->getDouble();}
    else{InitialTimeToMitosisSCC=5760;cerr<<"InitialTimeToMitosisSCC not specified, default value set to: "<<InitialTimeToMitosisSCC<<"steps"<<endl;}
    if(_xmlData->findElement("MeanTimeToMitosisCheckSCC")){MeanTimeToMitosisCheckSCC=_xmlData->getFirstElement("
```

```
            MeanTimeToMitosisCheckSCC")->getDouble();}
        else{MeanTimeToMitosisCheckSCC=5760;cerr<<"MeanTimeToMitosisCheckSCC not
            specified, default value set to: "<<MeanTimeToMitosisCheckSCC<<"steps
            "<<endl;}
        if(_xmlData->findElement("VolumeMultiplierForGrowth")){
            VolumeMultiplierForGrowth=_xmlData->getFirstElement("
            VolumeMultiplierForGrowth")->getDouble();}
        else{ VolumeMultiplierForGrowth=2;cerr<<"VolumeMultiplierForGrowth not
            specified, default value set to: "<<VolumeMultiplierForGrowth<<"steps
            "<<endl;}
        if(_xmlData->findElement("DivisionThresholdMultiplier")){
            DivisionThresholdMultiplier=_xmlData->getFirstElement("
            DivisionThresholdMultiplier")->getDouble();}
        else{ DivisionThresholdMultiplier=0.9;cerr<<"DivisionThresholdMultiplier
            not specified, default value set to: "<<DivisionThresholdMultiplier<<
            "steps"<<endl;}
        if(_xmlData->findElement("InitialCellOutputFileName")){
            InitialCellOutputFileName=_xmlData->getFirstElement("
            InitialCellOutputFileName")->getText();}
        if(_xmlData->findElement("TimeToCellDivisionFileName")){
            TimeToCellDivisionFileName=_xmlData->getFirstElement("
            TimeToCellDivisionFileName")->getText();}
        if(_xmlData->findElement("TimeToWriteCells")){TimeToWriteCells=_xmlData->
            getFirstElement("TimeToWriteCells")->getDouble();}
        else{TimeToWriteCells=5800;cerr<<"TimeToWriteCells not specified, default
             value set to: "<<TimeToWriteCells<<"steps"<<endl;}
        if(_xmlData->findElement("DivisionTimeDistribution")){
            DivisionTimeDistribution=_xmlData->getFirstElement("
            DivisionTimeDistribution")->getText();}
        else{std::string str ("Exponential");DivisionTimeDistribution=str;cerr<<"
            DivisionTimeDistribution not specified, default distribution set to:
            "<<DivisionTimeDistribution<<endl;}
        if(_xmlData->findElement("ErlangShapeParameter")){ErlangShapeParameter=
            _xmlData->getFirstElement("ErlangShapeParameter")->getDouble();}
        else{ErlangShapeParameter=1;cerr<<"ErlangShapeParameter not specified,
            default value set to: "<<ErlangShapeParameter<<endl;}
        if(_xmlData->findElement("SurfacevolumeRatio")){SurfacevolumeRatio=
            _xmlData->getFirstElement("SurfacevolumeRatio")->getText();}
        else{std::string str ("Sphere");DivisionTimeDistribution=str;cerr<<"
            DivisionTimeDistribution not specified, default distribution set to:
            "<<SurfacevolumeRatio<<endl;}
    }
```

**Listing 2.2.2:** SCC cell division initialisation Steppable - update

The 'start' function, Listing 2.2.3, is called at the beginning of the simulation, allocating variables of the steppable object. In this case there are no parameters to be allocated.

```
    void MitosisInitialisation::start(){
    }
```

**Listing 2.2.3:** SCC cell division initialisation Steppable - start

The step function is the main component of the steppable that updates the overall system and cell properties at each timestep. Variables related to probability distribution of cell division time and rate of growth of surface area are given in Listing 2.2.4.

```
    void MitosisInitialisation::step(const unsigned int currentStep){
        CellInventory *cellInventoryPtr=& potts->getCellInventory();
        CellInventory::cellInventoryIterator cInvItr;
        CellG *cell;

        const char *distributionname;
        distributionname=DivisionTimeDistribution.c_str();
        const char *Dis1 = "Exponential";
```

```
        const char *Dis2 = "Erlang";

        const char *SurfVolRatio;
        SurfVolRatio=SurfacevolumeRatio.c_str();
        const char *Ratio1 = "Sphere";
        const char *Ratio2 = "User";
```

**Listing 2.2.4:** SCC cell division initialisation Steppable - step 1

Cell cycle clocks cannot all be allocated exactly at time zero because this would not generate a truly random sample of cell cycle stages even if the length of the cell cycle was random in each case. The cells would all begin in phase and would only slowly drift out of phase over multiple cell divisions. We require them to be out of phase from each other from the beginning. To generate this random phasing, cells begin the simulation not in the cell cycle. The length of delay until permanently entering the cell cycle is itself a uniformly distributed random variable with mean value determined from the xml input, generally selected to be of the same magnitude as the cell cycle itself. The randomness of this delay causes each cell to enter the cell cycle out of phase from all others. During this delay, the cell does not grow and cannot divide. Only once the cell has completed this delay phase, will it enter the cell cycle with the potential to grow and divide over a random period of time.

In most cases, the variables in the steppable will require some form of initialisation and so the steppable functionality is different for MCS zero compared to all others (Listing 2.2.5). The random delay for each cell is generated at MCS zero. Each SCC is given an InitialConditionCell status of true and allocated a timer value between 0 and the InitialTimeToMitosis parameter, generated from the Uniform distribution. The cell's constant target volume and volume energy parameter are set from user input. The target surface area of the cell is defined either in terms of what the surface area should be if the cell is a sphere or to be a pre-defined user determined size. VCAF surface area, volume and energy parameters are set here and remain fixed for the entire simulation as VCAF division is ignored.

```
        if(currentStep==0){
            for(cInvItr=cellInventoryPtr->cellInventoryBegin() ; cInvItr !=
                cellInventoryPtr->cellInventoryEnd() ;++cInvItr ){
                cell=cellInventoryPtr->getCell(cInvItr);
                if (cell->type == SCCtype){
                    cell->InitialConditionCell=true;
                    cell->targetVolume=SCCVolume;
                    cell->lambdaVolume=SCCLambdaVolume;
                    if(strcmp(SurfVolRatio, Ratio1)==0){
                        double radius=pow(3.0*(double)cell->targetVolume
                            /4.0/3.14,1/3.0);
                        double SurfVolRatioMultiplier=1.0;
                        cell->targetSurface=SurfVolRatioMultiplier*4*3.14*pow(
                            radius,2);
                    }
                    else if(strcmp(SurfVolRatio, Ratio2)==0){
                        cell->targetSurface=SCCSurface;
                    }
                    cell->lambdaSurface=SCCLambdaSurface;
                        double t_rand=RandomGenerator::Obj().getUniformRV( 0.0,
                            1.0 );
                        cell->InitialTimeToMitosis = (int) (t_rand*
                            InitialTimeToMitosisSCC);
                        cell->OriginalTargetVolume = cell->targetVolume;
```

```
                        cell->OriginalTargetSurface = cell->targetSurface;
                    }
                else if(cell->type==VCAFtype){
                    cell->targetVolume=VCAFVolume;
                    cell->lambdaVolume=VCAFLambdaVolume;
                    cell->targetSurface=VCAFSurface;
                    cell->lambdaSurface=VCAFLambdaSurface;
                }
            }
        }
```

**Listing 2.2.5:** SCC cell division initialisation Steppable - step 2

The initial counter to cell division initiation is counted down for each SCC (Listing 2.2.6). When the counter reaches zero, the InitialConditionCell label is set to false. This cell then enters the cell division cycle of repeated growth and division. The cell is first allocated a time to cell division. In idealised conditions, the cell will grow in size over this time period and then divide. The time to mitosis is randomly generated for each cell, either from the exponential or Erlang distributions, depending on the xml input. The mean of the distribution and, in the case of the Erlang distribution, the shape parameter are xml inputs. The final distributions are derived from randomly generated uniformly distributed variables as in Equations (2.2) and (2.4).

```

        else{
            for(cInvItr=cellInventoryPtr->cellInventoryBegin() ; cInvItr !=
                cellInventoryPtr->cellInventoryEnd() ;++cInvItr ){
                cell=cellInventoryPtr->getCell(cInvItr);
                if (cell->InitialConditionCell==true){
                    cell->InitialTimeToMitosis--;
                    if (cell->InitialTimeToMitosis==0){
                        cell->InitialConditionCell=false;
                        if (cell->type == SCCtype){
                            if(strcmp(distributionname, Dis1)==0){
                                double t_rand=RandomGenerator::Obj().getUniformRV
                                    ( 0.0, 1.0 );
                                double t_rand_exp=-log(t_rand);
                                cell->TimeToMitosisCheck = (int) (t_rand_exp*
                                    MeanTimeToMitosisCheckSCC);
                            }
                            else if(strcmp(distributionname, Dis2)==0){
                                double t_rand=1.0;
                                    for( int a = 1; a < (int)ErlangShapeParameter
                                        +1; a = a + 1 ){
                                        double t_randsingle=RandomGenerator::Obj
                                            ().getUniformRV( 0.0, 1.0 );
                                        t_rand=t_rand*t_randsingle;
                                    }
                                    double t_rand_exp=-log(t_rand)/
                                        ErlangShapeParameter;
                                    cell->TimeToMitosisCheck = (int) (t_rand_exp*
                                        MeanTimeToMitosisCheckSCC);
                            }
                        }
                    }
                }
```

**Listing 2.2.6:** SCC cell division initialisation Steppable - step 3

Most of the cell growth functionality of a cell is decribed by Listing 2.2.7. For a MCS, the cell's mitosis timer is increased by 1 and its maximum volume (the volume the cell is trying to grow to in order to divide) is allocated. Based on the cell's time to mitosis (TimeToMitosisCheck), the current mitosis timer and the cell's maximum volume, its volume gradient is calculated as

$$\begin{aligned}&\text{volumegradient} = \text{originaltargetvolume}+\\ &(\text{volumemultiplierforgrowth} - 1) \times \text{originaltargetvolume} \times \frac{\text{mitosistimer}}{\text{timetomitosischeck}}. \quad (2.7)\end{aligned}$$

No cell's target volume can be lower than its initial target volume upon cell initialisation. Additionally, no cell's volume can exceed 10% of its current volume. If the cell is unable to grow due to space constraints then it may be much smaller than it should be according to its allocated time to mitosis, the size it should be at that point in time for the allocated time to mitosis and its actual current size. In these cases its target volume would be significantly higher than its actual volume. The constraint that stops a cell's target volume being more than 10% greater than the cell's current volume stops the cell from ballooning in size if the space constraints are rapidly removed. Following the calculation of the cell's current target volume, the cell's target surface area is calculated as described in 2.1, either to have the same ratio between newly initialised surface area and current target surface area as the volume would or such that the volume-surface area ratio is as in a sphere.

If a cell's volume increases beyond its division threshold then it divides (Listing 2.2.7). The cell target volume, target surface area and mitosis timer are all reset and the random time to mitosis for both daughter cells are randomly generated via either an exponential or Erlang distribution. The dividing cell is then given a true label for the variable dividethiscellinpython. This can then be sent to a python steppable, linked to the cc3d file, that divides the cell and allocates all parameters to the daughter cells.

```
else if(cell->InitialConditionCell==false){
    double cellvolumeplus=1.1*cell->volume;
    cell->mitosistimer++;
    cell->MaxCellVolume=cell->OriginalTargetVolume*
        VolumeMultiplierForGrowth;
    double volumegradient=cell->OriginalTargetVolume+(
        VolumeMultiplierForGrowth-1)*cell->OriginalTargetVolume/
        cell->TimeToMitosisCheck*cell->mitosistimer;
    cell->targetVolume=(int)(min(min(volumegradient,max(
        cellvolumeplus,double(cell->OriginalTargetVolume))),
        double(cell->MaxCellVolume)));
    if(strcmp(SurfVolRatio, Ratio1)==0){
        double radius=pow(3.0*(double)cell->targetVolume
            /4.0/3.14,1/3.0);
        double SurfVolRatioMultiplier=1.0;
        cell->targetSurface=SurfVolRatioMultiplier*4*3.14*pow(
            radius,2);
    }
    else if(strcmp(SurfVolRatio, Ratio2)==0){
        cell->targetSurface=(int)(cell->targetVolume/cell->
            OriginalTargetVolume*cell->OriginalTargetSurface);
    }
    if(cell->volume>cell->MaxCellVolume*
        DivisionThresholdMultiplier){
        cell->DivideThisCellInPython=true;
        cell->targetVolume=cell->OriginalTargetVolume;
        cell->targetSurface=cell->OriginalTargetSurface;
        cell->mitosistimer=0;
        if (cell->type == SCCtype){
            if(strcmp(distributionname, Dis1)==0){
                double t_rand=RandomGenerator::Obj().getUniformRV
                    ( 0.0, 1.0 );
```

```
                                    double t_rand_exp=-log(t_rand);
                                    cell->TimeToMitosisCheck = (int) (t_rand_exp*
                                        MeanTimeToMitosisCheckSCC);
                                    double t_rand2=RandomGenerator::Obj().
                                        getUniformRV( 0.0, 1.0 );
                                    double t_rand_exp2=-log(t_rand2);
                                    cell->DaughterCellTimer = (int) (t_rand_exp2*
                                        MeanTimeToMitosisCheckSCC);
                                }
                                else if(strcmp(distributionname, Dis2)==0){
                                    double t_rand=1.0;
                                    for( int a = 1; a < (int)ErlangShapeParameter+1;
                                        a = a + 1 ){
                                        double t_randsingle=RandomGenerator::Obj().
                                            getUniformRV( 0.0, 1.0 );
                                        t_rand=t_rand*t_randsingle;
                                    }
                                    double t_rand_exp=-log(t_rand)/
                                        ErlangShapeParameter;
                                    cell->TimeToMitosisCheck = (int) (t_rand_exp*
                                        MeanTimeToMitosisCheckSCC);

                                    double t_rand2=1.0;
                                    for( int a = 1; a < (int)ErlangShapeParameter+1;
                                        a = a + 1 ){
                                        double t_randsingle2=RandomGenerator::Obj().
                                            getUniformRV( 0.0, 1.0 );
                                        t_rand2=t_rand2*t_randsingle2;
                                    }
                                    double t_rand_exp2=-log(t_rand2)/
                                        ErlangShapeParameter;
                                    cell->DaughterCellTimer = (int)(t_rand_exp2*
                                        MeanTimeToMitosisCheckSCC);
                                }
                            }
                        }
                    }
                }
            }
```

**Listing 2.2.7:** SCC cell division initialisation Steppable - step 4

Finally, in Listing 2.2.8, when the initialisation simulation reaches a specified number of MCS, the initialisation files are printed. This includes the pif file recording the pixel locations of every cell in the simulation and a cell division output file documenting the random time to mitosis, current mitosis timer, the cell's maximum volume and the cell's current target volume and surface area. These files are then taken as initial condition files and used by the steppable described in Subsection 2.2.2.

```

        if(currentStep==TimeToWriteCells){
            const char *outputname_pif;.
            outputname_pif=InitialCellOutputFileName.c_str();
            InitialCellOutputFile.open(outputname_pif,ofstream::out);
            if (!InitialCellOutputFile.is_open()){cerr<<"could not open "<<
                InitialCellOutputFileName<<"!!"<<endl;}
                const char *outputname_time;
                outputname_time=TimeToCellDivisionFileName.c_str();
                TimeToCellDivisionFile.open(outputname_time,ofstream::out);
            if (!TimeToCellDivisionFile.is_open()){cerr<<"could not open "<<
                TimeToCellDivisionFile<<"!!"<<endl;}
            for(cInvItr=cellInventoryPtr->cellInventoryBegin() ; cInvItr !=
                cellInventoryPtr->cellInventoryEnd() ;++cInvItr ){
                cell=cellInventoryPtr->getCell(cInvItr);
                if(cell->type == SCCtype){
```

```
                    TimeToCellDivisionFile<<setw(5)<<cell->id<<setw(15)<<cell->
                        mitosistimer<<setw(15)<<cell->TimeToMitosisCheck<<setw
                        (10)<<cell->MaxCellVolume<<setw(10)<<cell->targetVolume<<
                        setw(10)<<cell->targetSurface<<endl;
                }
                set<PixelTrackerData> cellPixels=pixelTrackerAccessorPtr->get(
                    cell->extraAttribPtr)->pixelSet;
                for(set<PixelTrackerData >::iterator sitr=cellPixels.begin() ;
                    sitr != cellPixels.end() ;++sitr){
                    int xx=sitr->pixel.x;
                    int yy=sitr->pixel.y;
                    int zz=sitr->pixel.z;
                    int ct = cell->type;
                    if(ct==1){
                        InitialCellOutputFile <<setw(5)<<cell->id<<setw(7)<<"SCC"
                            <<setw(5)<< xx<<setw(5)<< xx<<setw(5)<< yy <<setw(5)
                            <<      yy<<setw(5)<<zz<<setw(5)<<zz<<endl;
                    }
                    else if(ct==2){
                        InitialCellOutputFile <<setw(5)<<cell->id<<setw(7)<<"VCAF
                            "<<setw(5)<< xx<<setw(5)<< xx<<setw(5)<< yy <<setw(5)
                            <<     yy<<setw(5)<<zz<<setw(5)<<zz<<endl;
                    }
                }
            }
            InitialCellOutputFile.close();
            TimeToCellDivisionFile.close();
        }
    }
```

**Listing 2.2.8:** SCC cell division initialisation Steppable - step 5

##### 2.2.2 Post Initialised Steppable

The post initialised steppable, MitosisPostInitialise, works very similarly to the initialisation steppable, except it is used during the simulations themselves, rather than generating the initial condition files. The functions below are given in the MitosisPostInitialise.cpp file. Due to the similarity with the initial condition generating steppable of Subsection 2.2.1, the listings are described more briefly.

The initiation function is given in Listing 2.2.9 while the update function in Listing 2.2.10 contains a similar set of parameters to Listing 2.2.2. The only significant additions are input so that the cell division initialisation text file created in Listing 2.2.8 can be read in and such that output cell division data can be printed to a text file.

```
void MitosisPostInitialise::init(Simulator *simulator, CC3DXMLElement *_xmlData)
    {

    potts = simulator->getPotts();
    sim=simulator;
    cellFieldG = (WatchableField3D<CellG *> *)potts->getCellFieldG();

    simulator->registerSteerableObject(this);
    update(_xmlData);
}
```

**Listing 2.2.9:** SCC cell division Steppable - init

```

void MitosisPostInitialise::update(CC3DXMLElement *_xmlData, bool _fullInitFlag){
    if(_xmlData->findElement("SCCtype")){SCCtype=_xmlData->getFirstElement("
        SCCtype")->getDouble();}
```

```
else{SCCtype=1;cerr<<"SCCtype not specified , defaul t value set to: "<<SCCtype<<endl;}
if(_xmlData->findElement("VCAFtype")){VCAFtype=_xmlData->getFirstElement("VCAFtype")->getDouble();}
else{VCAFtype=2;cerr<<"VCAFtype not specified , defaul t value set to: "<<VCAFtype<<endl;}
if(_xmlData->findElement("SCCVolume")){SCCVolume=_xmlData->getFirstElement("SCCVolume")->getDouble();}
else{SCCVolume=550;cerr<<"SCCVolume not specified , defaul t value set to: "<<SCCVolume<<endl;}
if(_xmlData->findElement("VCAFVolume")){VCAFVolume=_xmlData->getFirstElement("VCAFVolume")->getDouble();}
else{VCAFVolume=800;cerr<<"VCAFVolume not specified , defaul t value set to: "<<VCAFVolume<<endl;}
if(_xmlData->findElement("SCCLambdaVolume")){SCCLambdaVolume=_xmlData->getFirstElement("SCCLambdaVolume")->getDouble();}
else{SCCLambdaVolume=1;cerr<<"SCCLambdaVolume not specified , defaul t value set to: "<<SCCLambdaVolume<<endl;}
if(_xmlData->findElement("VCAFLambdaVolume")){VCAFLambdaVolume=_xmlData->getFirstElement("VCAFLambdaVolume")->getDouble();}
else{VCAFLambdaVolume=1;cerr<<"VCAFLambdaVolume not specified , defaul t value set to: "<<VCAFLambdaVolume<<endl;}
if(_xmlData->findElement("SCCSurface")){SCCSurface=_xmlData->getFirstElement("SCCSurface")->getDouble();}
else{SCCSurface=550;cerr<<"SCCSurface not specified , default value set to: "<<SCCSurface<<endl;}
if(_xmlData->findElement("VCAFSurface")){VCAFSurface=_xmlData->getFirstElement("VCAFSurface")->getDouble();}
else{VCAFSurface=700;cerr<<"VCAFSurface not specified , default value set to: "<<VCAFSurface<<endl;}
if(_xmlData->findElement("SCCLambdaSurface")){SCCLambdaSurface=_xmlData->getFirstElement("SCCLambdaSurface")->getDouble();}
else{SCCLambdaSurface=1;cerr<<"SCCLambdaSurface not specified , default value set to: "<<SCCLambdaSurface<<endl;}
if(_xmlData->findElement("VCAFLambdaSurface")){VCAFLambdaSurface=_xmlData->getFirstElement("VCAFLambdaSurface")->getDouble();}
else{VCAFLambdaSurface=1;cerr<<"VCAFLambdaSurface not specified , default value set to: "<<VCAFLambdaSurface<<endl;}
if(_xmlData->findElement("MeanTimeToMitosisCheckSCC")){MeanTimeToMitosisCheckSCC=_xmlData->getFirstElement("MeanTimeToMitosisCheckSCC")->getDouble();}
else{MeanTimeToMitosisCheckSCC=2880;cerr<<"MeanTimeToMitosisCheckSCC not specified , default value set to: "<<MeanTimeToMitosisCheckSCC<<"steps"<<endl;}
if(_xmlData->findElement("VolumeMultiplierForGrowth")){VolumeMultiplierForGrowth=_xmlData->getFirstElement("VolumeMultiplierForGrowth")->getDouble();}
else{ VolumeMultiplierForGrowth=2;cerr<<"VolumeMultiplierForGrowth not specified , default value set to: "<<VolumeMultiplierForGrowth<<"steps"<<endl;}
if(_xmlData->findElement("DivisionThresholdMultiplier")){DivisionThresholdMultiplier=_xmlData->getFirstElement("DivisionThresholdMultiplier")->getDouble();}
else{ DivisionThresholdMultiplier=0.8;cerr<<"DivisionThresholdMultiplier not specified , default value set to: "<<DivisionThresholdMultiplier<<"steps"<<endl;}
if(_xmlData->findElement("MitosisStageInputFileName")){MitosisStageInputFileName=_xmlData->getFirstElement("MitosisStageInputFileName")->getText();}
if(_xmlData->findElement("TimeOfCellDivisionFileName")){TimeOfCellDivisionFileName=_xmlData->getFirstElement("TimeOfCellDivisionFileName")->getText();}
if(_xmlData->findElement("DivisionTimeDistribution")){DivisionTimeDistribution=_xmlData->getFirstElement("DivisionTimeDistribution")->getText();}
else{std::string str ("Exponential");DivisionTimeDistribution=str;cerr<<"DivisionTimeDistribution not specified , default distribution set to: "<<DivisionTimeDistribution<<endl;}
if(_xmlData->findElement("ErlangShapeParameter")){ErlangShapeParameter=_xmlData->getFirstElement("ErlangShapeParameter")->getDouble();}
```

```
    else{ErlangShapeParameter=1;cerr<<"ErlangShapeParameter not specified ,
        default value set to: "<<ErlangShapeParameter<<endl;}
    if(_xmlData->findElement("SurfacevolumeRatio")){SurfacevolumeRatio=_xmlData->
        getFirstElement("SurfacevolumeRatio")->getText();}
    else{std::string str ("Sphere");DivisionTimeDistribution=str;cerr<<"
        DivisionTimeDistribution not specified , default distribution set to: "<<
        SurfacevolumeRatio<<endl;}
}
```

**Listing 2.2.10:** SCC cell division Steppable - update

The start function 2.2.11 is not empty in this instance - the text file is opened where cell division output data is written every time a cell divides.

```

void MitosisPostInitialise::start(){

    const char *outputname;
    outputname=TimeOfCellDivisionFileName.c_str();
    TimeOfCellDivisionFile.open(outputname,ofstream::out);
    if (!TimeOfCellDivisionFile.is_open()){cerr<<"could not open "<<
        TimeOfCellDivisionFile<<"!!"<<endl;}
}
```

**Listing 2.2.11:** SCC cell division Steppable - start

```

void MitosisPostInitialise::step(const unsigned int currentStep){

    CellInventory *cellInventoryPtr=& potts->getCellInventory();
    CellInventory::cellInventoryIterator cInvItr;
    CellG *cell;
    const char *distributionname;
    distributionname=DivisionTimeDistribution.c_str();
    const char *Dis1 = "Exponential";
    const char *Dis2 = "Erlang";

    const char *SurfVolRatio;
    SurfVolRatio=SurfacevolumeRatio.c_str();
    const char *Ratio1 = "Sphere";
    const char *Ratio2 = "User";
```

**Listing 2.2.12:** SCC cell division Steppable - step 1

The beginning of the step function 2.2.12 initialises variables as in Listing 2.2.4. For MCS zero, the cell division input file (generated by the cell division initialisation of subsection 2.2.1 and Listing 2.2.8) is read and cell properties are allocated to individual cells (Listing 2.2.13).

```
    if(currentStep==0){
        const char *inputname;
        inputname=MitosisStageInputFileName.c_str();
        MitosisStageInputFile.open(inputname,ifstream::in);
        if(!MitosisStageInputFile.is_open()){cerr<<"could not open "<<
            MitosisStageInputFile<<" !!"<<endl;}
            int counter=-1;
            for(cInvItr=cellInventoryPtr->cellInventoryBegin() ; cInvItr !=
                cellInventoryPtr->cellInventoryEnd() ;++cInvItr ){
                cell=cellInventoryPtr->getCell(cInvItr);
                if (cell->type == SCCtype){
                    cell->targetVolume=SCCVolume;
                    cell->lambdaVolume=SCCLambdaVolume;
                    if(strcmp(SurfVolRatio, Ratio1)==0){
                    double radius=pow(3.0*(double)cell->targetVolume
                        /4.0/3.14,1/3.0);
                    double SurfVolRatioMultiplier=1.0;
```

```
                    cell->targetSurface=SurfVolRatioMultiplier*4*3.14*pow(radius
                        ,2);
                }
                else if(strcmp(SurfVolRatio, Ratio2)==0){
                    cell->targetSurface=SCCSurface;
                }
                cell->lambdaSurface=SCCLambdaSurface;
                cell->OriginalTargetVolume = cell->targetVolume;
                cell->OriginalTargetSurface = cell->targetSurface;
                MitosisStageInputFile>>cell->id;
                MitosisStageInputFile>>cell->mitosistimer;
                MitosisStageInputFile>>cell->TimeToMitosisCheck;
                MitosisStageInputFile>>cell->MaxCellVolume;
                MitosisStageInputFile>>cell->targetVolume;
                MitosisStageInputFile>>cell->targetSurface;
            }
            else if(cell->type == VCAFtype){
                cell->targetVolume=VCAFVolume;
                cell->lambdaVolume=VCAFLambdaVolume;
                cell->targetSurface=VCAFSurface;
                cell->lambdaSurface=VCAFLambdaSurface;
            }
        }
        MitosisStageInputFile.close();
    }
```

**Listing 2.2.13:** SCC cell division Steppable - step 2

The key mechanisms of the cell division steppable are decribed in Listing 2.2.14. It is functionally similar to Listing 2.2.7. The only addition is that if a cell divides then its cell id, final count on the mitosis timer, allocated time to mitosis and final actual volume and target volumes are printed into the output text file (opened in Listing 2.2.11).

```
        else{
            for(cInvItr=cellInventoryPtr->cellInventoryBegin() ; cInvItr !=
                cellInventoryPtr->cellInventoryEnd() ;++cInvItr ){
                cell=cellInventoryPtr->getCell(cInvItr);
                if (cell->type == SCCtype){
                    double cellvolumeplus=1.1*cell->volume;
                    cell->mitosistimer++;
                    cell->MaxCellVolume=cell->OriginalTargetVolume*
                        VolumeMultiplierForGrowth;
                    double volumegradient=cell->OriginalTargetVolume+(
                        VolumeMultiplierForGrowth-1)*cell->OriginalTargetVolume/
                        cell->TimeToMitosisCheck*cell->mitosistimer;
                    cell->targetVolume=(int)(min(min(volumegradient,max(
                        cellvolumeplus,double(cell->OriginalTargetVolume))),
                        double(cell->MaxCellVolume)));
                    if(strcmp(SurfVolRatio, Ratio1)==0){
                    double  radius=pow(3.0*(double)cell->targetVolume
                        /4.0/3.14,1/3.0);
                        double SurfVolRatioMultiplier=1.0;
                        cell->targetSurface=SurfVolRatioMultiplier*4*3.14*pow(
                            radius,2);
                    }
                    else if(strcmp(SurfVolRatio, Ratio2)==0){
                        cell->targetSurface=(int)(cell->targetVolume/cell->
                            OriginalTargetVolume*cell->OriginalTargetSurface);
                    }
                    if(cell->volume>cell->MaxCellVolume*
                        DivisionThresholdMultiplier){
                        TimeOfCellDivisionFile<<setw(5)<<cell->id<<setw(15)<<cell
                            ->mitosistimer<<setw(15)<<cell->TimeToMitosisCheck<<
                            setw(10)<<cell->volume<<setw(10)<<cell->targetVolume
                            <<setw(10)<<endl;
                        cell->DivideThisCellInPython=true;
                        cell->targetVolume=cell->OriginalTargetVolume;
```

```
                        cell->targetSurface=cell->OriginalTargetSurface;
                        cell->mitosistimer=0;
                        if(strcmp(distributionname, Dis1)==0){
                            double t_rand=RandomGenerator::Obj().getUniformRV(
                                0.0, 1.0 );
                            double t_rand_exp=-log(t_rand);
                            cell->TimeToMitosisCheck = (int) (t_rand_exp*
                                MeanTimeToMitosisCheckSCC);
                            double t_rand2=RandomGenerator::Obj().getUniformRV(
                                0.0, 1.0 );
                            double t_rand_exp2=-log(t_rand2);
                            cell->DaughterCellTimer = (int) (t_rand_exp2*
                                MeanTimeToMitosisCheckSCC);
                        }
                        else if(strcmp(distributionname, Dis2)==0){
                            double t_rand=1.0;
                            for( int a = 1; a < (int)ErlangShapeParameter+1; a =
                                a + 1 ){
                            double t_randsingle=RandomGenerator::Obj().
                                getUniformRV( 0.0, 1.0 );
                            t_rand=t_rand*t_randsingle;
                        }
                        double t_rand_exp=-log(t_rand)/ErlangShapeParameter;
                        cell->TimeToMitosisCheck = (int) (t_rand_exp*
                            MeanTimeToMitosisCheckSCC);

                        double t_rand2=1.0;

                        for( int a = 1; a < (int)ErlangShapeParameter+1; a = a +
                            1 ){
                            double t_randsingle2=RandomGenerator::Obj().
                                getUniformRV( 0.0, 1.0 );
                            t_rand2=t_rand2*t_randsingle2;
                        }
                        double t_rand_exp2=-log(t_rand2)/ErlangShapeParameter;
                        cell->DaughterCellTimer = (int)(t_rand_exp2*
                            MeanTimeToMitosisCheckSCC);
                    }
                }
            }
        }
    }
}
```

**Listing 2.2.14:** SCC cell division Steppable - step 3

### 3 Cellular preferred directionality

#### 3.1 Methodology

Both SCC cells and VCAFs have a preferred direction of movement that is context dependent (organotypic or spheroid). Each cell type also has a kinesis level, with SCC kinesis being constant and VCAF kinesis being stimulated by neighbouring cells whilst VCAFs also repulse each other. The stimulation of kinesis and selection of preferred direction is explained in Section 4, and VCAF repulsion is explained in Section 5. This section explains how the directionality is provided, given a preferred direction and known level of kinesis stimulation for VCAFs.

The normalised direction of movement, $\hat{\mathbf{u}}$, is the unit vector in the direction of centre of mass translocation. For each pixel copy attempt, the normalised direction of movement of a cell is calculated according to equations

$$\mathbf{u} = \mathbf{CM}_{\text{after}} - \mathbf{CM}_{\text{before}} \tag{3.1}$$

$$\hat{\mathbf{u}} = \frac{\mathbf{u}}{\|\mathbf{u}\|} \tag{3.2}$$

where $\mathbf{CM}_{\text{after}}$ and $\mathbf{CM}_{\text{before}}$ give the centre of mass of the cell after and before the copy attempt respectively. This is the same for both the cell that protruded into the pixel (newCell) and the cell that retracted from the pixel (oldCell).

The change of energy for newCell and oldCell are calculated from the projection of the normalised direction of movement on to the normalised preferred direction, $\hat{\mathbf{p}}$ and the cell's kinesis energy parameter such that

$$\Delta E_i = \begin{cases} -\lambda_{\text{taxis},i} \hat{\mathbf{u}} \cdot \hat{\mathbf{p}} & \text{if Cell } i \text{ is not medium} \\ 0 & \text{otherwise} \end{cases} \tag{3.3}$$

where $i$ represents either newCell or oldCell. The contribution of directionality bias to the energy change of the pixel copy will then be calculated as

$$\Delta E = \Delta E_{\text{New}} + \Delta E_{\text{Old}}. \tag{3.4}$$

#### 3.2 Code Documentation

##### 3.2.1 Steppable

The steppables required for directionality preferred cellular direction and kinesis are explained in Subsections 4.2.1 and 4.2.2.

##### 3.2.2 Plugin

The plugin responsible for calculating the energy contribution of chemotaxis in SCC cells and kinesis in VCAFs on the current pixel copy attempt is DirectionalBiasPlugin. Functions listed below are found in the DirectionalBiasPlugin.cpp file. The initiation function of the plugin identifies and links the simulation main frame to the plugin, and initiates any necessary plugins that are utilised in the functionality of this plugin (Listing 3.2.1).

```
void DirectionalBiasPlugin::init(Simulator *simulator, CC3DXMLElement *_xmlData)
{
    potts = simulator->getPotts();
    bool pluginAlreadyRegisteredFlag;
    Plugin *plugin=Simulator::pluginManager.get("VolumeTracker",&
        pluginAlreadyRegisteredFlag);
    if(!pluginAlreadyRegisteredFlag)
plugin->init(simulator);
    potts->registerEnergyFunctionWithName(this, toString());
    xmlData=_xmlData;
    simulator->registerSteerableObject(this);
    update(_xmlData);
}
```

**Listing 3.2.1:** Cellular preferred directionality plugin - init

The update function reads the simulation inputs in the xml simulation setup file, and allocates them to the plugin. Here, as elsewhere, VCAF and SCC cell types are given as inputs (Listing 3.2.2).

```
void DirectionalBiasPlugin::update(CC3DXMLElement *_xmlData, bool _fullInitFlag)
{
    if(_xmlData->findElement("SCCtype")){SCCtype=_xmlData->getFirstElement("
        SCCtype")->getDouble();}
    else{SCCtype=1;cerr<<"SCCtype not specified, default value set to: "<<SCCtype
        <<endl;}
    if(_xmlData->findElement("VCAFtype")){VCAFtype=_xmlData->getFirstElement("
        VCAFtype")->getDouble();}
    else{VCAFtype=2;cerr<<"VCAFtype not specified, default value set to: "<<
        VCAFtype<<endl;}
}
```

**Listing 3.2.2:** Cellular preferred directionality plugin - update

The 'changeEnergy' function defines how the energy of the copy attempt will be calculated. As inputs, the function takes the current pixel address, pt, that the copy attempt is trying to change the value of, the address of the cell that is trying to copy into the pixel, newCell, and the address of the cell that is currently occupying the pixel before the copy attempt, oldCell. The function initially checks if newCell and oldCell are the same and returns the energy change as zero if they are, avoiding any unnecessary calculation steps (Listing 3.2.3).

```
double DirectionalBiasPlugin::changeEnergy(const Point3D &pt,const CellG *newCell
    ,const CellG *oldCell)
{
    double energy = 0.0;
    if (oldCell == newCell) return 0;
```

**Listing 3.2.3:** Cellular preferred directionality plugin - energy change 1

If the cells are different, the energy of the new state is calculated. If newCell is a valid cell, the centre of mass before the pixel copy attempt and after the pixel copy attempt is calculated. The newCell is the cell that is to copy itself to the pixel, and the new centre of mass of newCell is calculated including this additional pixel. Then the unit vector of cellular direction of movement is calculated, and taking the projection of cellular direction on the preferred cellular direction, the energy contribution is calculated as in Equations(3.1) - (3.4), (Listing 3.2.4).

```
    if (newCell){
        if((int)newCell->type==SCCtype ||   (int)newCell->type==VCAFtype){
            float Cxpre=newCell->xCM/newCell->volume;
            float Cypre=newCell->yCM/newCell->volume;
            float Czpre=newCell->zCM/newCell->volume;
            float Cxpost=(newCell->xCM+pt.x)/(newCell->volume+1);
            float Cypost=(newCell->yCM+pt.y)/(newCell->volume+1);
            float Czpost=(newCell->zCM+pt.z)/(newCell->volume+1);
            float dCM[3]={Cxpost-Cxpre,Cypost-Cypre,Czpost-Czpre};
            float dCMmag=pow((dCM[0]*dCM[0]+dCM[1]*dCM[1]+dCM[2]*dCM[2]), (float)
                0.5);
            dCM[0]=dCM[0]/dCMmag;dCM[1]=dCM[1]/dCMmag;dCM[2]=dCM[2]/dCMmag;
            float dotp=dCM[0]*newCell->taxisidir[0]+dCM[1]*newCell->taxisidir[1]+
                dCM[2]*newCell->taxisidir[2];
            energy=energy-newCell->lambdataxis*dotp;
        }
    }
```

**Listing 3.2.4:** Cellular preferred directionality plugin - energy change 2

In a similar manner, the energy contribution of the cellular direction of oldCell is calculated. Here, oldCell loses one pixel and the centre of mass is calculated accordingly (Listing 3.2.5).

```

    if (oldCell){
        if((int)oldCell->type==SCCtype || (int)oldCell->type==VCAFtype){
            float Cxpre=oldCell->xCM/oldCell->volume;
            float Cypre=oldCell->yCM/oldCell->volume;
            float Czpre=oldCell->zCM/oldCell->volume;
            float Cxpost=(oldCell->xCM-pt.x)/(oldCell->volume-1);
            float Cypost=(oldCell->yCM-pt.y)/(oldCell->volume-1);
            float Czpost=(oldCell->zCM-pt.z)/(oldCell->volume-1);
            float dCM[3]={Cxpost-Cxpre,Cypost-Cypre,Czpost-Czpre};
            float dCMmag=pow((dCM[0]*dCM[0]+dCM[1]*dCM[1]+dCM[2]*dCM[2]), (float)
                0.5);
           dCM[0]=dCM[0]/dCMmag;dCM[1]=dCM[1]/dCMmag;dCM[2]=dCM[2]/dCMmag;
            float dotp=dCM[0]*oldCell->taxisidir[0]+dCM[1]*oldCell->taxisidir[1]+
              dCM[2]*oldCell->taxisidir[2];
            energy=energy-oldCell->lambdataxis*dotp;
        }
    }
    return energy;
}
```

**Listing 3.2.5:** Cellular preferred directionality plugin - energy change 3

### 4 Cell directionality and kinesis determination

#### 4.1 Methodology

##### 4.1.1 Kinesis

The taxis level of SCC cells is defined at the initiation of simulation, and is not updated. However, each VCAF has a different level of kinesis depending on the environmental conditions it is subjected to. Upon stimulation by other VCAFs and SCC cells, a VCAF increases its motility between a low basal level and a maximum threshold. This stimulation does not require direct contact and stimulation level is inversely proportional to distance from neighbouring cells. The biochemical basis of this stimulation is not a subject of current study.

The kinesis parameter for each VCAF decays towards it basal value with a defined decay rate, $\eta_{kin}$, and increases towards its maximum value via stimulation. To calculate the stimulation of kinesis energy of a given VCAF, $m$, distances to the centre of masses of all other VCAFs and SCC cells are first calculated. The distance between VCAF $m$ and neighbouring cell $n$ is calculated as

$$d_{m,n} = \|\mathbf{CM}_m - \mathbf{CM}_n\| \tag{4.1}$$

where $\mathbf{CM}_m$ gives the centroid location for VCAF $m$ and $\mathbf{CM}_n$ the centroid location for neighbouring cell $n$. The stimulation effect, $\nu_{\text{kin}_{m,n}}$, for neighbouring cell, $n$, on VCAF $m$, is given by

$$\nu_{\text{kin}_{m,n}} = \begin{cases} \frac{L_{\text{kin}} - d_{m,n}}{L_{\text{kin}}} & \text{if } m \neq n \text{ and } d_{m,n} < L_{\text{kin}} \\ 0 & \text{otherwise.} \end{cases} \tag{4.2}$$

The stimulation effect of cell $n$ on VCAF $m$ decreases with distance until it reaches a cut-off distance, $L_{\text{kin}}$, whereby cells no longer stimulate neighbouring VCAFs.

The overall stimulation level is calculated as the sum of stimulation effects $\nu_{\text{kin}_{m,n}}$ from all neighbouring cells, scaled with a defined stimulation scaling factor, $\alpha_{\text{kin}}$. The kinesis of VCAF $m$ at MCS $t$, is then updated based on the current level of kinesis $\lambda_{\text{kin},m,t-1}$ at MCS $t-1$, the stimulation from neighbouring cells and the kinesis decay rate such that

$$\lambda_{\text{kin},m,t} = \lambda_{\text{kin},m,t-1} + \alpha_{\text{kin}} \sum_{n=1}^{N} \nu_{\text{kin}_{m,n}} - \eta_{\text{kin}} \tag{4.3}$$

where $N$ is the total number of cells in the simulation. The kinesis level for any VCAF is capped at defined minimum and maximum rates such that

$$\lambda_{\text{kin},m,t} = \begin{cases} \lambda_{\text{kin,min}} & \text{if } \lambda_{\text{kin},m,t} < \lambda_{\text{kin,min}}, \\ \lambda_{\text{kin,max}} & \text{if } \lambda_{\text{kin},m,t} > \lambda_{\text{kin,max}}. \end{cases} \tag{4.4}$$

##### 4.1.2 Directionality

The preferred direction of SCCs and VCAFs differs between the organotypic and spheroid environments and in each case is dynamic. For cell $m$ and MCS $t$, the unit preferred direction is given by $\hat{\mathbf{p}}_{m,t}$.

**Organotypic context** The preferred direction update of VCAFs is dependent on four directional components. Firstly, persistence of preferred directionality of cell $m$ is incorporated by including a weighted component of the preferred direction at MCS $t-1$ given by $\hat{\mathbf{p}}_{m,t-1}$. Secondly, a constant chemotactic response vector $\hat{\mathbf{c}}$ is included. Thirdly, a random vector, $\hat{\mathbf{r}}_{m,t}$ is included to introduce random noise into the cell's direction. For a cell, $m$ this random direction is updated at each MCS, $t$, by randomly generating components between -1 and 1 from the uniform distribution and then normalising. Finally, a velocity component, $\hat{\mathbf{v}}_{m,t}$, composed of the velocity of the cell at the previous timestep is included such that

$$\hat{\mathbf{v}}_{m,t} = \frac{\mathbf{CM}_{m,t-1} - \mathbf{CM}_{m,t-2}}{\|\mathbf{CM}_{m,t-1} - \mathbf{CM}_{m,t-2}\|}. \tag{4.5}$$

Here, $\mathbf{CM}_{m,t}$ gives the centroid location of cell $m$ at time $t$. Weightings for the cell's previous preferred directionality and previous velocity mimic persistence and polarity. VCAFs are also repulsed by other VCAFs, influencing their direction. Hence, in conjuction with VCAF-VCAF repulsion (Section 5), memory in preferred direction and previous velocity aid VCAFs in turning away from other VCAFs. The VCAF's direction is then given by

$$\mathbf{p}_{m,t} = (1 - w_c - w_r - w_v)\hat{\mathbf{p}}_{m,t-1} + w_v\hat{\mathbf{v}}_{m,t} + w_r\hat{\mathbf{r}}_{m,t} + w_c\hat{\mathbf{c}}, \tag{4.6}$$

$$\hat{\mathbf{p}}_{m,t} = \frac{\mathbf{p}_t}{\|\mathbf{p}_t\|}, \tag{4.7}$$

where $0 \leq w_c$, $w_r$, $w_v \leq 1$ are the weightings for chemotactic response, random directionality and persistent velocity respectively and $w_c + w_r + w_v = 1$.

The preferred direction update of SCC cells is derived similarly but is composed of only two of the above components: constant chemotactic response and a random directionality that is updated at every timestep. The derivation of preferred directionality for SCCs is thus equivalent to (4.6) where $w_v = 0$ and $w_c + w_r = 1$.

**Spheroid context** In the spheroid context, in the case of VCAFs, the constant chemotactic response vector $\hat{\mathbf{c}}$ is replaced by a radial chemotactic response vector $\hat{\omega}$, calculated similarly to velocity persistence (Equation (4.5)), as

$$\hat{\omega}_{m,t} = \frac{\mathbf{CM}_{m,t} - \mathbf{CM}_{S,t}}{\|\mathbf{CM}_{m,t} - \mathbf{CM}_{S,t}\|}. \tag{4.8}$$

Here, $\mathbf{CM}_{m,t}$ again gives the centroid location of cell $m$ at time $t$ whilst $\mathbf{CM}_{S,t}$ gives the centroid location of the whole spheroid. That is, the radial chemotactic direction is determined by a cell's current location relative to the centre of the spheroid. Equation (4.6) is then given by

$$\mathbf{p}_{m,t} = (1 - w_\omega - w_r - w_v)\hat{\mathbf{p}}_{m,t-1} + w_v \hat{\mathbf{v}}_{m,t} + w_r \hat{\mathbf{r}}_{m,t} + w_\omega \hat{\omega}, \tag{4.9}$$

with equivalent parameter constraints as described above.

In the case of SCCs, the directionality is derived as in the organotypic context, with a uniform chemotactic response and a random direction. However, in the spheroid context the SCCs also have a radial directionality component derived as in Equation (4.8).

#### 4.2 Code Documentation

##### 4.2.1 Organotypic Directionality and Kinesis Steppable

The steppable responsible for updating the kinesis energy parameter and the preferred direction of VCAFs in the organotypic context is OrganotypicKinesisDirectionality with the functions below being given in the OrganotypicKinesisDirectionality.cpp file. The initiation function of the steppable identifies and links the simulation main frame to the steppable (Listing 4.2.1).

```
void OrganotypicKinesisDirectionality::init(Simulator *simulator, CC3DXMLElement
    *_xmlData) {
    potts = simulator->getPotts();
    sim=simulator;
    cellFieldG = (WatchableField3D<CellG *> *)potts->getCellFieldG();
    simulator->registerSteerableObject(this);
    update(_xmlData);
}
```

**Listing 4.2.1:** VCAF kinesis stimulation Steppable - init

The update function reads the input in the simulation setup xml file, and allocates them to the steppable. Here, SCC and VCAF cell types, taxis energy parameter of SCC cells, minimum and maximum values of kinesis energy parameter for VCAFs, chemotaxis direction of both cell types, the decay rate of VCAF kinesis stimulation, scaling factor for VCAF kinesis stimulation, relevant weights of random noise, chemotaxis and previous velocity contributions to each cell type's orientation update, and the cut-off distance for VCAF kinesis stimulation are given as inputs to this function (Listing 4.2.2).

```

void OrganotypicKinesisDirectionality::update(CC3DXMLElement *_xmlData, bool _fullInitFlag){

    if(_xmlData->findElement("SCCtype")){SCCtype=_xmlData->getFirstElement("SCCtype")->getDouble();}
    else{SCCtype=1;cerr<<"SCCtype not specified, default value set to: "<<SCCtype<<endl;}
    if(_xmlData->findElement("VCAFtype")){VCAFtype=_xmlData->getFirstElement("VCAFtype")->getDouble();}
    else{VCAFtype=2;cerr<<"VCAFtype not specified, default value set to: "<<VCAFtype<<endl;}
    if(_xmlData->findElement("SCCLambdaTaxis"))
SCClambdataxis=_xmlData->getFirstElement("SCCLambdaTaxis")->getDouble();
    if(_xmlData->findElement("VCAFLambdaTaxis_min"))
VCAFlambdataxis_min=_xmlData->getFirstElement("VCAFLambdaTaxis_min")->getDouble();
    if(_xmlData->findElement("VCAFLambdaTaxis_max"))
VCAFlambdataxis_max=_xmlData->getFirstElement("VCAFLambdaTaxis_max")->getDouble();
    if(_xmlData->findElement("SCCchemotaxisdir_x"))
SCCchemotaxisdir[0]=_xmlData->getFirstElement("SCCchemotaxisdir_x")->getDouble();
    if(_xmlData->findElement("SCCchemotaxisdir_y"))
SCCchemotaxisdir[1]=_xmlData->getFirstElement("SCCchemotaxisdir_y")->getDouble();
    if(_xmlData->findElement("SCCchemotaxisdir_z"))
SCCchemotaxisdir[2]=_xmlData->getFirstElement("SCCchemotaxisdir_z")->getDouble();
    if(_xmlData->findElement("VCAFchemotaxisdir_x")){
VCAFchemotaxisdir[0]=_xmlData->getFirstElement("VCAFchemotaxisdir_x")->getDouble();
}
    else{VCAFchemotaxisdir[0]=0.0;cerr<<"VCAFchemotaxisdir_x not specified, default value set to: "<<VCAFchemotaxisdir[0]<<"."<<endl;
}
    if(_xmlData->findElement("VCAFchemotaxisdir_y")){
        VCAFchemotaxisdir[1]=_xmlData->getFirstElement("VCAFchemotaxisdir_y")->getDouble();
}
    else{VCAFchemotaxisdir[1]=0.0;cerr<<"VCAFchemotaxisdir_y not specified, default value set to: "<<VCAFchemotaxisdir[1]<<"."<<endl;
}
    if(_xmlData->findElement("VCAFchemotaxisdir_z")){
VCAFchemotaxisdir[2]=_xmlData->getFirstElement("VCAFchemotaxisdir_z")->getDouble();
}
    else{VCAFchemotaxisdir[2]=-1.0;cerr<<"VCAFchemotaxisdir_z not specified, default value set to: "<<VCAFchemotaxisdir[2]<<"."<<endl;
}
    if(_xmlData->findElement("LambdaDecayRate"))
rateoflambdadecay=_xmlData->getFirstElement("LambdaDecayRate")->getDouble();
    if(_xmlData->findElement("RepulsionScalingFactor"))
repulsionscalefactor=_xmlData->getFirstElement("RepulsionScalingFactor")->getDouble();
    if(_xmlData->findElement("Weight_RandomNoise"))
wrand=_xmlData->getFirstElement("Weight_RandomNoise")->getDouble();
    if(_xmlData->findElement("Weight_PreviousVelocity"))
wprev=_xmlData->getFirstElement("Weight_PreviousVelocity")->getDouble();
    if(_xmlData->findElement("Weight_Chemotactic")){
wchem=_xmlData->getFirstElement("Weight_Chemotactic")->getDouble();
}
    else{wchem=0.0;cerr<<"Weight_Chemotactic not specified, default value set to: "<<wchem<<"."<<endl;
}
    if(_xmlData->findElement("Weight_RandomSCCNoise"))
wsccrand=_xmlData->getFirstElement("Weight_RandomSCCNoise")->getDouble();
```

```
    if(_xmlData->findElement("ThresholdDiameter"))
        thres_diameter=_xmlData->getFirstElement("ThresholdDiameter")->getDouble
            ();
        thres2=thres_diameter*thres_diameter;
    }

}
```

**Listing 4.2.2:** Organotypic kinesis and directionality steppable - update

There are no parameters to be allocated in the 'start' function of this steppable. The 'step' function carries out the necessary modifications described above. At the initial step, the cells have their kinesis energy parameters and initial preferred directions allocated. For SCC cells, the kinesis energy term is constant whilst for VCAFs it is initially assigned to be at the minimum value set in the xml input (Listings 4.2.3 and 4.2.4). For both cell types, the direction is allocated in terms of chemotactic and random directions which is then normalised. The previous centre of mass is updated as the current centroid location for each cell type.

```
void OrganotypicKinesisDirectionality::step(const unsigned int currentStep){
    CellInventory *cellInventoryPtr=& potts->getCellInventory();
    CellInventory::cellInventoryIterator cInvItr;
    CellG *cell;
    double mean=0.0;
    unsigned int cellCounter=0;
    for(cInvItr=cellInventoryPtr->cellInventoryBegin() ; cInvItr !=
        cellInventoryPtr->cellInventoryEnd() ;++cInvItr ){
        cell=cellInventoryPtr->getCell(cInvItr);
        if(currentStep==0){
            if((int)cell->type==SCCtype){
                cell->lambdataxis=SCClambdataxis;
                double mag=SCCchemotaxisdir[0]*SCCchemotaxisdir[0]+
                    SCCchemotaxisdir[1]*SCCchemotaxisdir[1]+SCCchemotaxisdir[2]*
                    SCCchemotaxisdir[2];
                mag=pow(mag,0.5);
                float SCCchemotaxisnorm[3];
                SCCchemotaxisnorm[0]=SCCchemotaxisdir[0]/mag;
                SCCchemotaxisnorm[1]=SCCchemotaxisdir[1]/mag;
                SCCchemotaxisnorm[2]=SCCchemotaxisdir[2]/mag;
                float r[3];
                r[0]=rand()%2000;r[0]=r[0]/1000.0;r[0]=r[0]-1;
                r[1]=rand()%2000;r[1]=r[1]/1000.0;r[1]=r[1]-1;
                r[2]=rand()%2000;r[2]=r[2]/1000.0;r[2]=r[2]-1;
                float rmag=pow((r[0]*r[0] + r[1]*r[1] + r[2]*r[2]),(float) 0.5);
                r[0]=r[0]/rmag;r[1]=r[1]/rmag;r[2]=r[2]/rmag;
                float wsccpre=1.0-wsccrand;
                cell->taxisidir[0]=wsccpre*SCCchemotaxisnorm[0]+wsccrand*r[0];
                cell->taxisidir[1]=wsccpre*SCCchemotaxisnorm[1]+wsccrand*r[1];
                cell->taxisidir[2]=wsccpre*SCCchemotaxisnorm[2]+wsccrand*r[2];
                double taxisdirmag = cell->taxisidir[0]*cell->taxisidir[0]+cell->
                    taxisidir[1]*cell->taxisidir[1]+cell->taxisidir[2]*cell->
                    taxisidir[2];
                taxisdirmag=pow(taxisdirmag,(double) 0.5);
                cell->taxisidir[0]=cell->taxisidir[0]/taxisdirmag;
                cell->taxisidir[1]=cell->taxisidir[1]/taxisdirmag;
                cell->taxisidir[2]=cell->taxisidir[2]/taxisdirmag;

                cell->previousCM[0]=cell->xCOM;
                cell->previousCM[1]=cell->yCOM;
                cell->previousCM[2]=cell->zCOM;
            }
```

**Listing 4.2.3:** Organotypic kinesis and directionality steppable - step 1

```
    else if((int)cell->type==VCAFtype){
        cell->lambdataxis=VCAFlambdataxis_min;

        double mag=VCAFchemotaxisdir[0]*VCAFchemotaxisdir[0]+VCAFchemotaxisdir
            [1]*VCAFchemotaxisdir[1]+VCAFchemotaxisdir[2]*VCAFchemotaxisdir[2];
        mag=pow(mag,0.5);
        float VCAFchemotaxisnorm[3];
        VCAFchemotaxisnorm[0]=VCAFchemotaxisdir[0]/mag;
        VCAFchemotaxisnorm[1]=VCAFchemotaxisdir[1]/mag;
        VCAFchemotaxisnorm[2]=VCAFchemotaxisdir[2]/mag;

        float r[3];
        r[0]=rand()%2000;r[0]=r[0]/1000.0;r[0]=r[0]-1;
        r[1]=rand()%2000;r[1]=r[1]/1000.0;r[1]=r[1]-1;
        r[2]=rand()%2000;r[2]=r[2]/1000.0;r[2]=r[2]-1;
        float rmag=pow((r[0]*r[0] + r[1]*r[1] + r[2]*r[2]),(float) 0.5);
        r[0]=r[0]/rmag;r[1]=r[1]/rmag;r[2]=r[2]/rmag;
        float wrandinitial=1.0-wchem;
        cell->taxisidir[0]=wchem*VCAFchemotaxisnorm[0]+wrandinitial*r[0];
        cell->taxisidir[1]=wchem*VCAFchemotaxisnorm[1]+wrandinitial*r[1];
        cell->taxisidir[2]=wchem*VCAFchemotaxisnorm[2]+wrandinitial*r[2];

        double taxisdirmag = cell->taxisidir[0]*cell->taxisidir[0]+cell->
            taxisidir[1]*cell->taxisidir[1]+cell->taxisidir[2]*cell->taxisidir
            [2];
        taxisdirmag=pow(taxisdirmag,(double) 0.5);
        cell->taxisidir[0]=cell->taxisidir[0]/taxisdirmag;
        cell->taxisidir[1]=cell->taxisidir[1]/taxisdirmag;
        cell->taxisidir[2]=cell->taxisidir[2]/taxisdirmag;

        cell->previousCM[0]=cell->xCOM;
        cell->previousCM[1]=cell->yCOM;
        cell->previousCM[2]=cell->zCOM;
        }
}
```

**Listing 4.2.4:** Organotypic kinesis and directionality steppable - step 2

For future timesteps the direction of each SCC cell is updated by generating a new random direction and normalising (4.2.5). If the SCC has just divided then its just_divided_taxis_label is set to true and the cell's previous centre of mass is updated and the label updated to false.

```
else if((int)cell->type==SCCtype){
    cell->lambdataxis=SCClambdataxis;
    double mag=SCCchemotaxisdir[0]*SCCchemotaxisdir[0]+SCCchemotaxisdir[1]*
        SCCchemotaxisdir[1]+SCCchemotaxisdir[2]*SCCchemotaxisdir[2];
    mag=pow(mag,0.5);
    float SCCchemotaxisnorm[3];
    SCCchemotaxisnorm[0]=SCCchemotaxisdir[0]/mag;
    SCCchemotaxisnorm[1]=SCCchemotaxisdir[1]/mag;
    SCCchemotaxisnorm[2]=SCCchemotaxisdir[2]/mag;

    float r[3];
    r[0]=rand()%2000;r[0]=r[0]/1000.0;r[0]=r[0]-1;
    r[1]=rand()%2000;r[1]=r[1]/1000.0;r[1]=r[1]-1;
    r[2]=rand()%2000;r[2]=r[2]/1000.0;r[2]=r[2]-1;
    float rmag=pow((r[0]*r[0] + r[1]*r[1] + r[2]*r[2]),(float) 0.5);
    r[0]=r[0]/rmag;r[1]=r[1]/rmag;r[2]=r[2]/rmag;

    float wsccpre=1.0-wsccrand;
    cell->taxisidir[0]=wsccpre*SCCchemotaxisnorm[0]+wsccrand*r[0];
    cell->taxisidir[1]=wsccpre*SCCchemotaxisnorm[1]+wsccrand*r[1];
    cell->taxisidir[2]=wsccpre*SCCchemotaxisnorm[2]+wsccrand*r[2];

```

```
    double taxisdirmag = cell->taxisidir[0]*cell->taxisidir[0]+cell->taxisidir
        [1]*cell->taxisidir[1]+cell->taxisidir[2]*cell->taxisidir[2];
    taxisdirmag=pow(taxisdirmag,(double) 0.5);
    cell->taxisidir[0]=cell->taxisidir[0]/taxisdirmag;
    cell->taxisidir[1]=cell->taxisidir[1]/taxisdirmag;
    cell->taxisidir[2]=cell->taxisidir[2]/taxisdirmag;

    if (cell->just_divided_taxis_label == true ){
        cell->previousCM[0]=cell->xCOM;
        cell->previousCM[1]=cell->yCOM;
        cell->previousCM[2]=cell->zCOM;
        cell->just_divided_taxis_label=false;
    }

}
```

**Listing 4.2.5:** Organotypic kinesis and directionality steppable - step 3

In the following steps, all VCAFs are scanned to update the energy parameter and direction with the constraints of current environment. Initially, the kinesis energy parameter is decayed (Listing 4.2.6).

```

else if((int)cell->type==VCAFtype){
    cell->lambdataxis=cell->lambdataxis-rateoflambdadecay;
```

**Listing 4.2.6:** Organotypic kinesis and directionality steppable - step 4

Then all the cells of the simulation are scanned to check for their distances to the current VCAF, and the cumulative, distance dependent stimulation (or repulsion as the cells stimulate the VCAF to move away from themselves) parameter is calculated (Listing 4.2.7).

```

    double repulsion=0.0;
    CellInventory::cellInventoryIterator cInvItr2;
    CellG *cell2;
    for(cInvItr2=cellInventoryPtr->cellInventoryBegin() ; cInvItr2 !=
        cellInventoryPtr->cellInventoryEnd() ;++cInvItr2 ){
        cell2=cellInventoryPtr->getCell(cInvItr2);
        if (((int)cell2->type==SCCtype || (int)cell2->type==VCAFtype) && cell2->
            id!=cell->id){
            float x=cell2->xCOM, y=cell2->yCOM, z=cell2->zCOM;
            float dx=(cell->xCOM-x),dy=(cell->yCOM-y),dz=(cell->zCOM-z);
            float d2=dx*dx+dy*dy+dz*dz;
            if(d2<thres2){
                float d=pow(d2,(float)0.5);
                repulsion=repulsion+(-(1.0/thres_diameter)*d+1);
            }
        }
    }
```

**Listing 4.2.7:** Organotypic kinesis and directionality steppable - step 5

Once all cells are checked for stimulating the current VCAF, the kinesis energy parameter is updated via the cumulative stimulation parameter and the scaling factor. The level of kinesis is capped within the minimum and maximum limits defined in the xml input file (Listing 4.2.8).

```

cell->lambdataxis=cell->lambdataxis+repulsionscalefactor*repulsion;
if(cell->lambdataxis<VCAFlambdataxis_min){
    cell->lambdataxis=VCAFlambdataxis_min;
}
else if(cell->lambdataxis>VCAFlambdataxis_max){
```

```
        cell->lambdataxis=VCAFlambdataxis_max;
}
```

**Listing 4.2.8:** Organotypic kinesis and directionality steppable - step 6

The preferred direction update for VCAFs is then initiated. The chemotactic vector is normalised. A random vector is generated and normalised for the stochastic noise. The unit direction velocity is calculated from the translocation of the cell centre of mass, the previous centre of mass recorded in each cell is updated here. Finally, the preferred direction is updated with the weighting defined in the xml input file, and normalised (Listing 4.2.9).

```

    double mag=VCAFchemotaxisdir[0]*VCAFchemotaxisdir[0]+VCAFchemotaxisdir[1]*
        VCAFchemotaxisdir[1]+VCAFchemotaxisdir[2]*VCAFchemotaxisdir[2];
    mag=pow(mag,0.5);
    float VCAFchemotaxisnorm[3];
    VCAFchemotaxisnorm[0]=VCAFchemotaxisdir[0]/mag;
    VCAFchemotaxisnorm[1]=VCAFchemotaxisdir[1]/mag;
    VCAFchemotaxisnorm[2]=VCAFchemotaxisdir[2]/mag;

    float r[3];
    r[0]=rand()%2000;r[0]=r[0]/1000.0;r[0]=r[0]-1;
    r[1]=rand()%2000;r[1]=r[1]/1000.0;r[1]=r[1]-1;
    r[2]=rand()%2000;r[2]=r[2]/1000.0;r[2]=r[2]-1;
    float rmag=pow((r[0]*r[0] + r[1]*r[1] + r[2]*r[2]),(float) 0.5);
    r[0]=r[0]/rmag;r[1]=r[1]/rmag;r[2]=r[2]/rmag;

    float vprev[3]={cell->xCOM-cell->previousCM[0],cell->yCOM-cell->previousCM
        [1],cell->zCOM-cell->previousCM[2]};
    float vprevmag=pow((vprev[0]*vprev[0] + vprev[1]*vprev[1] + vprev[2]*vprev
        [2]),(float)0.5);
    if(vprevmag==0){
        vprev[0]=0.0;vprev[1]=0.0;vprev[2]=0.0;
    }
    else{
        vprev[0]=vprev[0]/vprevmag;vprev[1]=vprev[1]/vprevmag;vprev[2]=vprev[2]/
            vprevmag;
    }
    cell->previousCM[0]=cell->xCOM;cell->previousCM[1]=cell->yCOM;cell->
        previousCM[2]=cell->zCOM;
    float wpre=1.0-wrand-wprev-wchem;

    cell->taxisidir[0]=wpre*cell->taxisidir[0]+wrand*r[0]+wprev*vprev[0]+wchem*
        VCAFchemotaxisnorm[0];
    cell->taxisidir[1]=wpre*cell->taxisidir[1]+wrand*r[1]+wprev*vprev[1]+wchem*
        VCAFchemotaxisnorm[1];
    cell->taxisidir[2]=wpre*cell->taxisidir[2]+wrand*r[2]+wprev*vprev[2]+wchem*
        VCAFchemotaxisnorm[2];
    double taxisdirmag = cell->taxisidir[0]*cell->taxisidir[0]+cell->taxisidir
        [1]*cell->taxisidir[1]+cell->taxisidir[2]*cell->taxisidir[2];
    taxisdirmag=pow(taxisdirmag,(double) 0.5);
    cell->taxisidir[0]=cell->taxisidir[0]/taxisdirmag;
    cell->taxisidir[1]=cell->taxisidir[1]/taxisdirmag;
    cell->taxisidir[2]=cell->taxisidir[2]/taxisdirmag;
    }
  }
}
```

**Listing 4.2.9:** Organotypic kinesis and directionality steppable - step 7

##### 4.2.2 Spheroid Directionality and Kinesis Steppable

The steppable for the spheroid context is similar but accounts for the slightly different preferred directions described in Section 4.1.2. In particular, the directionality in the spheroid context includes a radial component that requires tracking of the spheroid centroid (Equation (4.8)). The tracking of this spheroid centroid is carried out by the SpheroidCentroid (Subsection 4.2.3).

Due to the similarity between organotypic and spheroid contexts, the overall code does not significantly differ to that presented in Subsection 4.2.1 and as such, we present a reduced illustration of the spheroid code. The function responsible in the spheroid context is SpheroidKinesisDirectionality with the functions below being given in the SpheroidKinesisDirectionality.cpp file.

In the update function 4.2.10 we present only input variables that do not appear in the organotypic context. These include the centroid locations of the spheroid at time zero (needed for radial directionality), directionality weightings to account for differing directionality components and the SCC uniform directionality vector.

```
void SpheroidKinesisDirectionality::update(CC3DXMLElement *_xmlData, bool
    _fullInitFlag){

    if(_xmlData->findElement("SpheroidCentroid_x"))
SpheroidCentroidVars[0]=_xmlData->getFirstElement("SpheroidCentroid_x")->
    getDouble();
    if(_xmlData->findElement("SpheroidCentroid_y"))
SpheroidCentroidVars[1]=_xmlData->getFirstElement("SpheroidCentroid_y")->
    getDouble();
    if(_xmlData->findElement("SpheroidCentroid_z"))
SpheroidCentroidVars[2]=_xmlData->getFirstElement("SpheroidCentroid_z")->
    getDouble();
    if(_xmlData->findElement("Weight_RadialVelocityVCAF")){
wchem=_xmlData->getFirstElement("Weight_RadialVelocityVCAF")->getDouble();
}
    else{
wchem=0.0;cerr<<"Weight_RadialVelocityVCAF not specified, default value set to: "
    <<wchem<<"."<<endl;
}
    if(_xmlData->findElement("Weight_UniformVelocitySCC")){
wsccuniform=_xmlData->getFirstElement("Weight_UniformVelocitySCC")->getDouble();
}
    else{
wsccuniform=0.0;cerr<<"Weight_UniformVelocitySCC not specified, default value set
     to: "<<wsccuniform<<"."<<endl;
}
    if(_xmlData->findElement("Weight_RandomSCCNoise")){
wsccrand=_xmlData->getFirstElement("Weight_RandomSCCNoise")->getDouble();
}
    else{
wsccrand=0.0;cerr<<"Weight_RandomSCCNoise not specified, default value set to: "
    <<wsccrand<<"."<<endl;
}
    if(_xmlData->findElement("SCCuniformtaxisdir_x"))
SCCuniformtaxisdir[0]=_xmlData->getFirstElement("SCCuniformtaxisdir_x")->
    getDouble();
    else{
SCCuniformtaxisdir[0]=0.0;cerr<<"SCCuniformtaxisdir_x not specified, default
    value set to: "<<SCCuniformtaxisdir[0]<<"."<<endl;
}
    if(_xmlData->findElement("SCCuniformtaxisdir_y"))
SCCuniformtaxisdir[1]=_xmlData->getFirstElement("SCCuniformtaxisdir_y")->
    getDouble();
    else{
SCCuniformtaxisdir[1]=0.0;cerr<<"SCCuniformtaxisdir_y not specified, default
    value set to: "<<SCCuniformtaxisdir[1]<<"."<<endl;
}
    if(_xmlData->findElement("SCCuniformtaxisdir_z"))
SCCuniformtaxisdir[2]=_xmlData->getFirstElement("SCCuniformtaxisdir_z")->
    getDouble();
```

```
    else{
SCCuniformtaxisdir[2]=0.0;cerr<<"SCCuniformtaxisdir_z not specified, default
    value set to: "<<SCCuniformtaxisdir[2]<<"."<<endl;
    }
}
```

**Listing 4.2.10:** Spheroid kinesis and directionality steppable - update

Differences in the step function again reflect the different preferred directionality of cells between spheroid and organotypic contexts. Kinesis in the spheroid context is as in the organotypic context and as such ignored here. The initial directionality of each SCC at time zero is defined in 4.2.11 in terms of a random component, a uniform directional component and a radial directional component. The initial directionality of each VCAF is defined in 4.2.12 in terms of a random direction and a radial direction.

```

void SpheroidKinesisDirectionality::step(const unsigned int currentStep){
    CellInventory *cellInventoryPtr=& potts->getCellInventory();
    CellInventory::cellInventoryIterator cInvItr;
    CellG *cell;
    double mean=0.0;
    unsigned int cellCounter=0;
    for(cInvItr=cellInventoryPtr->cellInventoryBegin() ; cInvItr !=
        cellInventoryPtr->cellInventoryEnd() ;++cInvItr ){
    cell=cellInventoryPtr->getCell(cInvItr);
    if(currentStep==0){
        if((int)cell->type==SCCtype){
            cell->lambdataxis=SCClambdataxis;
            cell->previousCM[0]=cell->xCOM;
            cell->previousCM[1]=cell->yCOM;
            cell->previousCM[2]=cell->zCOM;
            float SCC_Spheroidsradialdir[3]={
                cell->previousCM[0]-SpheroidCentroidVars[0],
                cell->previousCM[1]-SpheroidCentroidVars[1],
                cell->previousCM[2]-SpheroidCentroidVars[2]
            };
            float SCC_Spheroidsradialdirmag=pow((SCC_Spheroidsradialdir[0]*
                SCC_Spheroidsradialdir[0]+ SCC_Spheroidsradialdir[1]*
                SCC_Spheroidsradialdir[1]+SCC_Spheroidsradialdir[2]*
                SCC_Spheroidsradialdir[2]),(float)0.5);
            SCC_Spheroidsradialdir[0]=SCC_Spheroidsradialdir[0]/
                SCC_Spheroidsradialdirmag;
            SCC_Spheroidsradialdir[1]=SCC_Spheroidsradialdir[1]/
                SCC_Spheroidsradialdirmag;
            SCC_Spheroidsradialdir[2]=SCC_Spheroidsradialdir[2]/
                SCC_Spheroidsradialdirmag;
            double mag=SCCuniformtaxisdir[0]*SCCuniformtaxisdir[0]+
                SCCuniformtaxisdir[1]*SCCuniformtaxisdir[1]+SCCuniformtaxisdir
                [2]*SCCuniformtaxisdir[2];
            mag=pow(mag,0.5);
            float SCCuniformtaxisnorm[3];
            if(mag==0){SCCuniformtaxisnorm[0]=0.0;SCCuniformtaxisnorm[1]=0.0;
                SCCuniformtaxisnorm[2]=0.0;
            }
            else{
                SCCuniformtaxisnorm[0]=SCCuniformtaxisdir[0]/mag;
                SCCuniformtaxisnorm[1]=SCCuniformtaxisdir[1]/mag;
                SCCuniformtaxisnorm[2]=SCCuniformtaxisdir[2]/mag;
            }
            float r[3];
            r[0]=rand()%2000;r[0]=r[0]/1000.0;r[0]=r[0]-1;
            r[1]=rand()%2000;r[1]=r[1]/1000.0;r[1]=r[1]-1;
            r[2]=rand()%2000;r[2]=r[2]/1000.0;r[2]=r[2]-1;
            float rmag=pow((r[0]*r[0] + r[1]*r[1] + r[2]*r[2]),(float) 0.5);
            r[0]=r[0]/rmag;r[1]=r[1]/rmag;r[2]=r[2]/rmag;
            float wsccpre=1.0-wsccrand-wsccuniform;
```

```
            cell->taxisidir[0]=wsccpre*SCC_Spheroidsradialdir[0]+wsccrand*r[0]+
                wsccuniform*SCCuniformtaxisnorm[0];
            cell->taxisidir[1]=wsccpre*SCC_Spheroidsradialdir[1]+wsccrand*r[1]+
                wsccuniform*SCCuniformtaxisnorm[1];
            cell->taxisidir[2]=wsccpre*SCC_Spheroidsradialdir[2]+wsccrand*r[2]+
                wsccuniform*SCCuniformtaxisnorm[2];
}
```

**Listing 4.2.11:** Spheroid kinesis and directionality steppable - step 1

```
        else if((int)cell->type==VCAFtype){
            cell->lambdataxis=VCAFlambdataxis_min;
            cell->previousCM[0]=cell->xCOM;
            cell->previousCM[1]=cell->yCOM;
            cell->previousCM[2]=cell->zCOM;
            float VCAF_Spheroidsradialdir[3]={
                cell->previousCM[0]-SpheroidCentroidVars[0],
                cell->previousCM[1]-SpheroidCentroidVars[1],
                cell->previousCM[2]-SpheroidCentroidVars[2]
            };
            float VCAF_Spheroidsradialdirmag=pow((VCAF_Spheroidsradialdir[0]*
                VCAF_Spheroidsradialdir[0] + VCAF_Spheroidsradialdir[1]*
                VCAF_Spheroidsradialdir[1] + VCAF_Spheroidsradialdir[2]*
                VCAF_Spheroidsradialdir[2]),(float)0.5);
            VCAF_Spheroidsradialdir[0]=VCAF_Spheroidsradialdir[0]/
                VCAF_Spheroidsradialdirmag;
            VCAF_Spheroidsradialdir[1]=VCAF_Spheroidsradialdir[1]/
                VCAF_Spheroidsradialdirmag;
            VCAF_Spheroidsradialdir[2]=VCAF_Spheroidsradialdir[2]/
                VCAF_Spheroidsradialdirmag;
            float r[3];
            r[0]=rand()%2000;r[0]=r[0]/1000.0;r[0]=r[0]-1;
            r[1]=rand()%2000;r[1]=r[1]/1000.0;r[1]=r[1]-1;
            r[2]=rand()%2000;r[2]=r[2]/1000.0;r[2]=r[2]-1;
            float rmag=pow((r[0]*r[0] + r[1]*r[1] + r[2]*r[2]),(float) 0.5);
            r[0]=r[0]/rmag;r[1]=r[1]/rmag;r[2]=r[2]/rmag;
            float wrandinitial=1.0-wchem;
            cell->taxisidir[0]=wchem*VCAF_Spheroidsradialdir[0]+wrandinitial*r
                [0];
            cell->taxisidir[1]=wchem*VCAF_Spheroidsradialdir[1]+wrandinitial*r
                [1];
            cell->taxisidir[2]=wchem*VCAF_Spheroidsradialdir[2]+wrandinitial*r
                [2];
            double taxisdirmag = cell->taxisidir[0]*cell->taxisidir[0]+cell->
                taxisidir[1]*cell->taxisidir[1]+cell->taxisidir[2]*cell->
                taxisidir[2];
            taxisdirmag=pow(taxisdirmag,(double) 0.5);
            cell->taxisidir[0]=cell->taxisidir[0]/taxisdirmag;
            cell->taxisidir[1]=cell->taxisidir[1]/taxisdirmag;
            cell->taxisidir[2]=cell->taxisidir[2]/taxisdirmag;
        }
    }
```

**Listing 4.2.12:** Spheroid kinesis and directionality steppable - step 2

For future timesteps the direction of each SCC cell is updated as in 4.2.13 to also take account of radial direction. Similarly, the directionality of each VCAF is updated in 4.2.14 to account for radial directionality. The division status of each SCC and the kinesis levels of each VCAF are as in the organotypic context and thus removed from these illustrations.

```
        else if((int)cell->type==SCCtype){
            cell->lambdataxis=SCClambdataxis;
            float SCC_Spheroidsradialdir[3]={cell->previousCM[0]-cell->
                SpheroidCentroidX, cell->previousCM[1]-cell->SpheroidCentroidY,
                cell->previousCM[2]-cell->SpheroidCentroidZ};
```

```
        float SCC_Spheroidsradialdirmag=pow((SCC_Spheroidsradialdir[0]*
            SCC_Spheroidsradialdir[0] + SCC_Spheroidsradialdir[1]*
            SCC_Spheroidsradialdir[1] + SCC_Spheroidsradialdir[2]*
            SCC_Spheroidsradialdir[2]),(float)0.5);
        SCC_Spheroidsradialdir[0]=SCC_Spheroidsradialdir[0]/
            SCC_Spheroidsradialdirmag;SCC_Spheroidsradialdir[1]=
            SCC_Spheroidsradialdir[1]/SCC_Spheroidsradialdirmag;
            SCC_Spheroidsradialdir[2]=SCC_Spheroidsradialdir[2]/
            SCC_Spheroidsradialdirmag;
        cell->previousCM[0]=cell->xCOM; cell->previousCM[1]=cell->yCOM; cell->
            previousCM[2]=cell->zCOM;
        double mag=SCCuniformtaxisdir[0]*SCCuniformtaxisdir[0]+
            SCCuniformtaxisdir[1]*SCCuniformtaxisdir[1]+SCCuniformtaxisdir
            [2]*SCCuniformtaxisdir[2];
        mag=pow(mag,0.5);
        float SCCuniformtaxisnorm[3];
        if(mag==0){
            SCCuniformtaxisnorm[0]=0.0;SCCuniformtaxisnorm[1]=0.0;
                SCCuniformtaxisnorm[2]=0.0;
        }
        else{
            SCCuniformtaxisnorm[0]=SCCuniformtaxisdir[0]/mag;
            SCCuniformtaxisnorm[1]=SCCuniformtaxisdir[1]/mag;
            SCCuniformtaxisnorm[2]=SCCuniformtaxisdir[2]/mag;
        }
        float r[3];
        r[0]=rand()%2000;r[0]=r[0]/1000.0;r[0]=r[0]-1;
        r[1]=rand()%2000;r[1]=r[1]/1000.0;r[1]=r[1]-1;
        r[2]=rand()%2000;r[2]=r[2]/1000.0;r[2]=r[2]-1;
        float rmag=pow((r[0]*r[0] + r[1]*r[1] + r[2]*r[2]),(float) 0.5);
        r[0]=r[0]/rmag;r[1]=r[1]/rmag;r[2]=r[2]/rmag;
        float wsccpre=1.0-wsccrand-wsccuniform;
        cell->taxisidir[0]=wsccpre*SCC_Spheroidsradialdir[0]+wsccrand*r[0]+
            wsccuniform*SCCuniformtaxisnorm[0];
        cell->taxisidir[1]=wsccpre*SCC_Spheroidsradialdir[1]+wsccrand*r[1]+
            wsccuniform*SCCuniformtaxisnorm[1];
        cell->taxisidir[2]=wsccpre*SCC_Spheroidsradialdir[2]+wsccrand*r[2]+
            wsccuniform*SCCuniformtaxisnorm[2];
        double taxisdirmag = cell->taxisidir[0]*cell->taxisidir[0]+cell->
            taxisidir[1]*cell->taxisidir[1]+cell->taxisidir[2]*cell->
            taxisidir[2];
        taxisdirmag=pow(taxisdirmag,(double) 0.5);
        cell->taxisidir[0]=cell->taxisidir[0]/taxisdirmag;
        cell->taxisidir[1]=cell->taxisidir[1]/taxisdirmag;
        cell->taxisidir[2]=cell->taxisidir[2]/taxisdirmag;
    }
```

**Listing 4.2.13:** Spheroid kinesis and directionality steppable - step 3

```
    else if((int)cell->type==VCAFtype){
        float VCAF_Spheroidsradialdir[3]={cell->previousCM[0]-cell->
            SpheroidCentroidX,cell->previousCM[1]-cell->SpheroidCentroidY,
            cell->previousCM[2]-cell->SpheroidCentroidZ};
        float VCAF_Spheroidsradialdirmag=pow((VCAF_Spheroidsradialdir[0]*
            VCAF_Spheroidsradialdir[0] + VCAF_Spheroidsradialdir[1]*
            VCAF_Spheroidsradialdir[1] + VCAF_Spheroidsradialdir[2]*
            VCAF_Spheroidsradialdir[2]),(float)0.5);
        VCAF_Spheroidsradialdir[0]=VCAF_Spheroidsradialdir[0]/
            VCAF_Spheroidsradialdirmag;
        VCAF_Spheroidsradialdir[1]=VCAF_Spheroidsradialdir[1]/
            VCAF_Spheroidsradialdirmag;
        VCAF_Spheroidsradialdir[2]=VCAF_Spheroidsradialdir[2]/
            VCAF_Spheroidsradialdirmag;
        float r[3];
        r[0]=rand()%2000;r[0]=r[0]/1000.0;r[0]=r[0]-1;
        r[1]=rand()%2000;r[1]=r[1]/1000.0;r[1]=r[1]-1;
        r[2]=rand()%2000;r[2]=r[2]/1000.0;r[2]=r[2]-1;
        float rmag=pow((r[0]*r[0] + r[1]*r[1] + r[2]*r[2]),(float) 0.5);
        r[0]=r[0]/rmag;r[1]=r[1]/rmag;r[2]=r[2]/rmag;
```

```
            float vprev[3]={cell->xCOM-cell->previousCM[0], cell->yCOM-cell->
                previousCM[1], cell->zCOM-cell->previousCM[2]};
            float vprevmag=pow((vprev[0]*vprev[0] + vprev[1]*vprev[1] + vprev[2]*
                vprev[2]),(float)0.5);
            if(vprevmag==0){
                vprev[0]=0.0;vprev[1]=0.0;vprev[2]=0.0;
            }
            else{
                vprev[0]=vprev[0]/vprevmag;vprev[1]=vprev[1]/vprevmag;vprev[2]=
                    vprev[2]/vprevmag;
            }
            cell->previousCM[0]=cell->xCOM;cell->previousCM[1]=cell->yCOM;cell->
                previousCM[2]=cell->zCOM;
            float wpre=1.0-wrand-wprev-wchem;
            cell->taxisidir[0]=wpre*cell->taxisidir[0]+wrand*r[0]+wprev*vprev[0]+
                wchem*VCAF_Spheroidsradialdir[0];
            cell->taxisidir[1]=wpre*cell->taxisidir[1]+wrand*r[1]+wprev*vprev[1]+
                wchem*VCAF_Spheroidsradialdir[1];
            cell->taxisidir[2]=wpre*cell->taxisidir[2]+wrand*r[2]+wprev*vprev[2]+
                wchem*VCAF_Spheroidsradialdir[2];
            double taxisdirmag = cell->taxisidir[0]*cell->taxisidir[0]+cell->
                taxisidir[1]*cell->taxisidir[1]+cell->taxisidir[2]*cell->
                taxisidir[2];
            taxisdirmag=pow(taxisdirmag,(double) 0.5);
            cell->taxisidir[0]=cell->taxisidir[0]/taxisdirmag;
            cell->taxisidir[1]=cell->taxisidir[1]/taxisdirmag;
            cell->taxisidir[2]=cell->taxisidir[2]/taxisdirmag;
        }
    }
}
```

**Listing 4.2.14:** Spheroid kinesis and directionality steppable - step 4

##### 4.2.3 Spheroid Centroid Steppable

The steppable that records the centroid location of the spheroid in the spheroid context is SpheroidCentroid with functions below being given in the SpheroidCentroid.cpp file. The initiation function of the steppable is given by Listing 4.2.15 as usual linking the simulation main frame to the steppable.

```

void SpheroidCentroid::init(Simulator *simulator, CC3DXMLElement *_xmlData) {
    potts = simulator->getPotts();
    sim=simulator;
    cellFieldG = (WatchableField3D<CellG *> *)potts->getCellFieldG();
    simulator->registerSteerableObject(this);
    update(_xmlData);
    bool pluginAlreadyRegisteredFlag;
    pixelTrackerPlugin=(PixelTrackerPlugin*)Simulator::pluginManager.get("
        PixelTracker",&pluginAlreadyRegisteredFlag); //this will load
        VolumeTracker plugin if it is not already loaded
    if(!pluginAlreadyRegisteredFlag)
        pixelTrackerPlugin->init(simulator);
        pixelTrackerAccessorPtr=pixelTrackerPlugin->getPixelTrackerAccessorPtr();
    }
}
```

**Listing 4.2.15:** Spheroid centroid Steppable - init

The update function (Listing 4.2.16) reads input in the simulation setup xml file, and allocates them to the steppable. Here, SCC and VCAF cell types are included. Additionally, the centroid location of the spheroid in $x$, $y$ and $z$ coordinates at initialisation is included (this can be calculated from the pif file prior to input).

```
void SpheroidCentroid::update(CC3DXMLElement *_xmlData, bool _fullInitFlag){
    if(_xmlData->findElement("SCCtype")){SCCtype=_xmlData->getFirstElement("SCCtype")->getDouble();}
    else{SCCtype=1;cerr<<"SCCtype not specified, default value set to: "<<SCCtype<<endl;}
    if(_xmlData->findElement("VCAFtype")){VCAFtype=_xmlData->getFirstElement("VCAFtype")->getDouble();}
    else{VCAFtype=2;cerr<<"VCAFtype not specified, default value set to: "<<VCAFtype<<endl;}
    if(_xmlData->findElement("SpheroidCentroid_x"))
SpheroidCentroidVar[0]=_xmlData->getFirstElement("SpheroidCentroid_x")->getDouble();
    if(_xmlData->findElement("SpheroidCentroid_y"))
SpheroidCentroidVar[1]=_xmlData->getFirstElement("SpheroidCentroid_y")->getDouble();
    if(_xmlData->findElement("SpheroidCentroid_z"))
SpheroidCentroidVar[2]=_xmlData->getFirstElement("SpheroidCentroid_z")->getDouble();
}
```

**Listing 4.2.16:** Spheroid centroid Steppable - update

There are no parameters to be allocated in the 'start' function of this steppable. The 'step' function carries out the necessary modifications described above. At the initial step, the centroid location of the spheroid, defined in Listing 4.2.16, is allocated to each cell (see Listing 4.2.17).

```
void SpheroidCentroid::step(const unsigned int currentStep){
    CellInventory *cellInventoryPtr=& potts->getCellInventory();
    CellInventory::cellInventoryIterator cInvItr;
    CellG *cell;
    if(currentStep==0){
        for(cInvItr=cellInventoryPtr->cellInventoryBegin() ; cInvItr !=
            cellInventoryPtr->cellInventoryEnd() ;++cInvItr ){
            cell=cellInventoryPtr->getCell(cInvItr);
            cell->SpheroidCentroidX=SpheroidCentroidVar[0];
            cell->SpheroidCentroidY=SpheroidCentroidVar[1];
            cell->SpheroidCentroidZ=SpheroidCentroidVar[2];
        }
    }
```

**Listing 4.2.17:** Spheroid centroid Steppable - step 1

For future timesteps the centroid location of the spheroid is calculated and the updated location allocated to each cell (Listing 4.2.18).

```
    vector <int> xcoord,ycoord,zcoord;
    for(cInvItr=cellInventoryPtr->cellInventoryBegin() ; cInvItr !=
        cellInventoryPtr->cellInventoryEnd() ;++cInvItr ){
        cell=cellInventoryPtr->getCell(cInvItr);
        if((int)cell->type==SCCtype){
            set<PixelTrackerData> cellPixels=pixelTrackerAccessorPtr->get(cell->
                extraAttribPtr)->pixelSet;
            for(set<PixelTrackerData>::iterator sitr=cellPixels.begin() ; sitr !=
                cellPixels.end() ;++sitr){
                int xx=sitr->pixel.x;int yy=sitr->pixel.y;int zz=sitr->pixel.z;
                xcoord.push_back(xx);ycoord.push_back(yy);zcoord.push_back(zz);
            }
        }
    }
    float centroidx=float(std::accumulate(xcoord.begin(),xcoord.end(),0.0))/float
        (xcoord.size());
    float centroidy=float(std::accumulate(ycoord.begin(),ycoord.end(),0.0))/float
        (ycoord.size());
```

```
    float centroidz=float(std::accumulate(zcoord.begin(),zcoord.end(),0.0))/float
        (zcoord.size());
    for(cInvItr=cellInventoryPtr->cellInventoryBegin() ; cInvItr !=
        cellInventoryPtr->cellInventoryEnd() ;++cInvItr ){
        cell=cellInventoryPtr->getCell(cInvItr);
        cell->SpheroidCentroidX=int(round(centroidx));
        cell->SpheroidCentroidY=int(round(centroidy));
        cell->SpheroidCentroidZ=int(round(centroidz));
    }
}
```

**Listing 4.2.18:** Spheroid centroid Steppable - step 2

##### 4.2.4 Plugin

The plugins utilising the functionality of VCAF kinesis stimulation are given in Section 3.2.2 and Section 5.2.

### 5 VCAF-VCAF repulsion

#### 5.1 Methodology

VCAFs are stimulated to move by both SCC cells and other VCAFs. SCC cells induce only kinesis without imposing a specific direction whilst VCAFS induce both kinesis and repulsion (see Section 4). The reulsive effect of neighbouring VCAFs on an individual VCAF is modelled via a VCAF repulsion energy term. For a VCAF, $m$, the centre of mass distance to a neighbouring VCAF, $n$, is calculated before and after the pixel copy attempt as

$$d_{\text{before}_{m,n}} = \|\mathbf{CM}_{\text{before}_m} - \mathbf{CM}_n\|, \tag{5.1}$$

$$d_{\text{after}_{m,n}} = \|\mathbf{CM}_{\text{after}_m} - \mathbf{CM}_n\|. \tag{5.2}$$

Distance dependent repulsion, $\rho$, is then calculated for each of these distances before and after the copy attempt such that

$$\rho_{\text{before}_{m,n}} = \begin{cases} \frac{L_{\text{rep}} - d_{\text{before}_{m,n}}}{L_{\text{rep}}} & \text{if } m \neq n \text{ and } d_{\text{before}_{m,n}} < L_{\text{rep}} \\ 0 & \text{otherwise} \end{cases}, \tag{5.3}$$

$$\rho_{\text{after}_{m,n}} = \begin{cases} \frac{L_{\text{rep}} - d_{\text{after}_{m,n}}}{L_{\text{rep}}} & \text{if } m \neq n \text{ and } d_{\text{after}_{m,n}} < L_{\text{rep}} \\ 0 & \text{otherwise} \end{cases}. \tag{5.4}$$

Here, the repulsion effect is zero beyond the defined cut-off distance, $L_{\text{rep}}$ and between 0 and 1 within this cut-off. Consider a single VCAF, $m$, in a system with $N_{VCAF}$ VCAFs. If $m$ is the new cell protruding into the pixel copy attempt then the energy contribution of repulsion on this new cell is

$$\Delta E_{new} = \alpha_\rho \sum_{n=1}^{N} \left( \rho_{\text{after}_{m,n}} - \rho_{\text{before}_{m,n}} \right). \tag{5.5}$$

If $m$ is the old cell retracting from the pixel copy attempt then the energy contribution of repulsion on this old cell is

$$\Delta E_{old} = \alpha_\rho \sum_{n=1}^{N} \left(\rho_{\text{after}_{m,n}} - \rho_{\text{before}_{m,n}}\right). \tag{5.6}$$

In each case, $\alpha_\rho$ gives the repulsion scaling factor. The larger the value, the more pulsive VCAFs are. The overall energy contribution of VCAF repulsion for this pixel copy is then calculated as

$$\Delta E = \Delta E_{\text{New}} + \Delta E_{\text{Old}}. \tag{5.7}$$

#### 5.2 Code Documentation

The plugin VCAFRepulsionPlugin (Subsection 5.2.2) is responsible for calculating the energy contributions of VCAF-VCAF repulsion. In order to calculate these contributions, the centre of mass positions of all VCAFs are recorded in the VCAFPositionListing steppable (Subsection 5.2.1).

##### 5.2.1 VCAF Position Listing Steppable

For scanning through VCAFs, the centre of mass positions of all VCAFs are recorded in a separate list, and are updated in each time step via the VCAFPositionListing steppable. The functions described below are given in the VCAFPositionListing.cpp file. The initiation function of the steppable identifies and links the simulation main frame to the steppable (Listing 5.2.5).

```
void VCAFPositionListing::init(Simulator *simulator, CC3DXMLElement *_xmlData) {
    potts = simulator->getPotts();
    sim=simulator;
    cellFieldG = (WatchableField3D<CellG *> *)potts->getCellFieldG();
    simulator->registerSteerableObject(this);
    update(_xmlData);
}
```

**Listing 5.2.1:** VCAF position listing steppable - init

The update function reads VCAF cell type as input from the simulation setup xml file (Listing 5.2.2).

```
void VCAFPositionListing::update(CC3DXMLElement *_xmlData, bool _fullInitFlag){
    if(_xmlData->findElement("VCAFtype")) VCAFtype=_xmlData->getFirstElement("
        VCAFtype")->getDouble();
}
```

**Listing 5.2.2:** VCAF position listing steppable - update

There are no variables that can be allocated before initialisation of the cells, hence the 'start' function is empty. The 'step' function updates the VCAF position list. At the initial step, a linked list of positions and id numbers of all VCAFs in the simulation is generated. For ease of access, the root (the first member) of the linked list is recorded by each VCAF in the simulation (Listing 5.2.3).

```
void VCAFPositionListing::step(const unsigned int currentStep){


    CellInventory *cellInventoryPtr=& potts->getCellInventory();
    CellInventory::cellInventoryIterator cInvItr;
    CellG *cell;
    if(currentStep==0){
        VCAFListroot = new VCAFpositionnode;
        VCAFpositionnode *conductor;
        conductor = VCAFListroot;
        bool initiatedfirstcell=false;
        for(cInvItr=cellInventoryPtr->cellInventoryBegin() ; cInvItr !=
            cellInventoryPtr->cellInventoryEnd() ;++cInvItr ){
            cell=cellInventoryPtr->getCell(cInvItr);
            if((int)cell->type==VCAFtype){
                if(!initiatedfirstcell){
                    initiatedfirstcell=true;
                }
            else{
                conductor->next = new VCAFpositionnode;
                conductor = conductor->next;
            }
            cell->VCAFListroot=VCAFListroot;
            conductor->id=cell->id;
            conductor->pos[0]=cell->xCM/cell->volume;
            conductor->pos[1]=cell->yCM/cell->volume;
            conductor->pos[2]=cell->zCM/cell->volume;
            }
        }
    }
```

**Listing 5.2.3:** VCAF position listing steppable - step 1

At each of the following steps, all VCAFs are scanned and the linked list of positions is updated accordingly (Listing 5.2.4).

```
    else{
        for(cInvItr=cellInventoryPtr->cellInventoryBegin() ; cInvItr !=
            cellInventoryPtr->cellInventoryEnd() ;++cInvItr ){
            cell=cellInventoryPtr->getCell(cInvItr);
            if((int)cell->type==VCAFtype){
                VCAFpositionnode *conductor = VCAFListroot;
                while ( conductor!= NULL ){
                    if(conductor->id==cell->id){
                        conductor->pos[0]=cell->xCM/cell->volume;
                        conductor->pos[1]=cell->yCM/cell->volume;
                        conductor->pos[2]=cell->zCM/cell->volume;
                    }
                    conductor = conductor->next;
                }
            }
        }
    }
```

**Listing 5.2.4:** VCAF position listing steppable - step 2

##### 5.2.2 VCAF-VCAF Repulsion Plugin

The plugin responsible for calculating the energy contribution of VCAF-VCAF repulsion on the current pixel copy attempt is VCAFRepulsionPlugin. The functions described below are given in the VCAFRepulsionPlugin.cpp file.

The initiation function of the plugin identifies and links the simulation main frame to the plugin, and initiates any necessary plugins that are utilised in the functionality of this plugin (Listing 5.2.5).

```
void VCAFRepulsionPlugin::init(Simulator *simulator, CC3DXMLElement *_xmlData)
{
    potts = simulator->getPotts();
    bool pluginAlreadyRegisteredFlag;
    Plugin *plugin=Simulator::pluginManager.get("VolumeTracker",&
        pluginAlreadyRegisteredFlag);
    if(!pluginAlreadyRegisteredFlag)
        plugin->init(simulator);
    potts->registerEnergyFunctionWithName(this, toString());
    xmlData=_xmlData;

    simulator->registerSteerableObject(this);
    update(_xmlData);
}
```

**Listing 5.2.5:** VCAF-VCAF repulsion Plugin - init

The update function reads the simulation inputs in the simulation setup xml file, and allocates them to the plugin. Here, the VCAF cell type, VCAF-VCAF repulsion energy parameter and the threshold diameter for repulsion are given (Listing 5.2.6).

```
void VCAFRepulsionPlugin::update(CC3DXMLElement *_xmlData, bool _fullInitFlag)
{
    if(_xmlData->findElement("VCAFtype"))
        VCAFtype=_xmlData->getFirstElement("VCAFtype")->getDouble();
    if(_xmlData->findElement("LambdaVCAFrepulsion"))
        lambdaVCAFrepulsion=_xmlData->getFirstElement("LambdaVCAFrepulsion")->
            getDouble();
    if(_xmlData->findElement("ThresholdDiameter"))
        thres_diameter=_xmlData->getFirstElement("ThresholdDiameter")->getDouble
            ();
    thres2=thres_diameter*thres_diameter;
}
```

**Listing 5.2.6:** VCAF-VCAF repulsion Plugin - update

The 'changeEnergy' function defines how the energy of the copy attempt will be calculated. The function initially checks if newCell (the one the copy attempt is attempting to change the pixel to) and oldCell (the one the copy attempt is attempting to change the pixel from) are the same, and returns the energy change as zero if they are (Listing 5.2.7).

```
double VCAFRepulsionPlugin::changeEnergy(const Point3D &pt,const CellG *newCell,
    const CellG *oldCell)
{
    double energy = 0.0;
    if (oldCell == newCell) return 0;
```

**Listing 5.2.7:** VCAF-VCAF repulsion Plugin - change energy 1

If oldCell and newCell are not the same, and if newCell is a VCAF, the repulsion energy contribution of newCell in the pixel copy attempt is calculated. The centre of mass of newCell before and after the pixel copy are calculated. The linked list of VCAF centre of mass positions is scanned and the distances to all

VCAFs other than newCell itself are calculated for both pre-pixel copy and post-pixel copy centre of mass. The change in repulsion as a function of centre of mass distances (refer to Equations (5.1) to (5.6)), is obtained as the difference between two states (Listing 5.2.8).

```
    if (newCell){
        if((int)newCell->type==VCAFtype){
            float repulsion=0.0;
            float CMpre[3];
            float CMpos[3];
            CMpre[0]=newCell->xCM/newCell->volume;
            CMpre[1]=newCell->yCM/newCell->volume;
            CMpre[2]=newCell->zCM/newCell->volume;
            CMpos[0]=(newCell->xCM+pt.x)/(newCell->volume+1);
            CMpos[1]=(newCell->yCM+pt.y)/(newCell->volume+1);
            CMpos[2]=(newCell->zCM+pt.z)/(newCell->volume+1);
            VCAFpositionnode *conductor = newCell->VCAFListroot;
            while ( conductor!= NULL){
                if(conductor->id!=newCell->id){
                    float dxpre=conductor->pos[0]-CMpre[0];
                    float dypre=conductor->pos[1]-CMpre[1];
                    float dzpre=conductor->pos[2]-CMpre[2];
                    if(dxpre<thres_diameter && dypre<thres_diameter && dzpre<
                        thres_diameter){
                        float d2pre=dxpre*dxpre+dypre*dypre+dzpre*dzpre;
                        if(d2pre<thres2){
                            float dxpos=conductor->pos[0]-CMpos[0];
                            float dypos=conductor->pos[1]-CMpos[1];
                            float dzpos=conductor->pos[2]-CMpos[2];
                            float d2pos=dxpos*dxpos+dypos*dypos+dzpos*dzpos;
                            float dpre=pow(d2pre,(float)0.5);
                            float dpos=pow(d2pos,(float)0.5);
                            float repulsionpre=(-(1.0/thres_diameter)*dpre+1);
                            float repulsionpos=(-(1.0/thres_diameter)*dpos+1);
                            repulsion=repulsion+(repulsionpos-repulsionpre);
                        }
                    }
                }
                conductor = conductor->next;
                }
            energy=energy+repulsion*lambdaVCAFrepulsion;
        }
    }
```

**Listing 5.2.8:** VCAF-VCAF repulsion Plugin - change energy 2

If oldCell is a VCAF, the energy contribution from oldCell is calculated in a similar manner to newCell. The energy contribution of VCAF-VCAF repulsion to the overall energy change of the pixel copy attempt is the sum of contributions from newCell and oldCell (refer to Equation (5.7)), (Listing 5.2.9).

```

if (oldCell){
    if((int)oldCell->type==VCAFtype){
        float repulsion=0.0;
        float CMpre[3];
        float CMpos[3];
        CMpre[0]=oldCell->xCM/oldCell->volume;
        CMpre[1]=oldCell->yCM/oldCell->volume;
        CMpre[2]=oldCell->zCM/oldCell->volume;
        CMpos[0]=(oldCell->xCM-pt.x)/(oldCell->volume-1);
        CMpos[1]=(oldCell->yCM-pt.y)/(oldCell->volume-1);
        CMpos[2]=(oldCell->zCM-pt.z)/(oldCell->volume-1);
        VCAFpositionnode *conductor = oldCell->VCAFListroot;
        while ( conductor!= NULL){
            if(conductor->id!=oldCell->id){
```

```
                    float dxpre=conductor->pos[0]-CMpre[0];
                    float dypre=conductor->pos[1]-CMpre[1];
                    float dzpre=conductor->pos[2]-CMpre[2];
                    if(dxpre<thres_diameter && dypre<thres_diameter && dzpre<
                        thres_diameter){
                        float d2pre=dxpre*dxpre+dypre*dypre+dzpre*dzpre;
                        if(d2pre<thres2){
                            float dxpos=conductor->pos[0]-CMpos[0];
                            float dypos=conductor->pos[1]-CMpos[1];
                            float dzpos=conductor->pos[2]-CMpos[2];
                            float d2pos=dxpos*dxpos+dypos*dypos+dzpos*dzpos;
                            float dpre=pow(d2pre,(float)0.5);
                            float dpos=pow(d2pos,(float)0.5);
                            float repulsionpre=(-(1.0/thres_diameter)*dpre+1);
                            float repulsionpos=(-(1.0/thres_diameter)*dpos+1);
                            repulsion=repulsion+(repulsionpos-repulsionpre);
                        }
                    }
                }
                conductor = conductor->next;
            }
            energy=energy+repulsion*lambdaVCAFrepulsion;
        }
    }
return energy;
}
```

**Listing 5.2.9:** VCAF-VCAF repulsion Plugin - change energy 3

### 6 ECM Remodellling

#### 6.1 Methodology

Matrix density is heterogeneous in the simulation whereby neither SCC cells, nor VCAFS can penetrate into dense regions of the ECM. This is introduced in the model by an energy term linking the concentration of ECM in the pixel subject to the pixel copy attempt to the energy cost of the copy attempt. If newCell is a valid cell (not medium), then this will bring an energy cost, as the cell is penetrating into ECM, while a valid oldCell will generate an energy benefit, as oldCell is retracting from the pixel such that

$$\Delta E_{\text{New}} = \lambda_{\text{ECM,NewCell}} c_{\text{ECM}}(p,t), \tag{6.1}$$

$$\Delta E_{\text{Old}} = \lambda_{\text{ECM,OldCell}} c_{\text{ECM}}(p,t), \tag{6.2}$$

where $c_{\text{ECM}}(p,t)$ gives the ECM concentration at pixel, $p$, and time, $t$. The energy contribution of ECM penetration to the overall energy change of the pixel copy will be the sum of energy changes from newCell and oldCell such that

$$\Delta E = \Delta E_{\text{New}} + \Delta E_{\text{Old}}. \tag{6.3}$$

ECM concentration at any pixel of the simulation grid, can be remodelled by cells via degradation and dislocation of ECM fibres. For a VCAF, $m$, at MCS $t$,the rates of degradation and pushing of the fibres depends on the current kinesis stimulation level such that

$$k_{\text{deg},m,t} = \alpha_{\text{deg,VCAF}} \frac{\lambda_{\text{kin},m,t} - \lambda_{\text{kin,min}}}{\lambda_{\text{kin,max}} - \lambda_{\text{kin,min}}} \tag{6.4}$$

$$k_{\text{push},m,t} = \alpha_{\text{push,VCAF}} \frac{\lambda_{\text{kin},m,t} - \lambda_{\text{kin,min}}}{\lambda_{\text{kin,max}} - \lambda_{\text{kin,min}}}. \tag{6.5}$$

At the basal level of kinesis, defined by $\lambda_{\text{kin,min}}$, there is no remodelling, and the rates are increased linearly with increasing kinesis stimulation, towards maximum degradation, $\alpha_{\text{deg,VCAF}}$, and pushing, $\alpha_{\text{push,VCAF}}$. For SCCs, the kinesis level remains constant. To maintain consistency between VCAF and SCC remodelling amid different levels of kinesis, we set $\lambda_{\text{kin,SCC}}$ to be the kinesis level at which VCAFs would travel at the same speed, on average, as SCCs. This allows equivalence between the maximum degradation and pushing rates between SCCs and VCAFs. This results in SCC degradation and pushing rates

$$k_{\text{deg},m,t} = \alpha_{\text{deg,SCC}} \frac{\lambda_{\text{kin,SCC}} - \lambda_{\text{kin,min}}}{\lambda_{\text{kin,max}} - \lambda_{\text{kin,min}}} \tag{6.6}$$

$$k_{\text{push},m,t} = \alpha_{\text{push,SCC}} \frac{\lambda_{\text{kin,SCC}} - \lambda_{\text{kin,min}}}{\lambda_{\text{kin,max}} - \lambda_{\text{kin,min}}}. \tag{6.7}$$

The concentration, $c_{ECM}(p, t+1)$ of ECM at a given pixel, $p$ and time point, $t+1$, is dependent on the concentrations in the previous time step $t$ and the rates of degradation and pushing in both the current pixel, $p$, and all its neighbours $q \in N_{\text{Neig},p}$. Firstly, the process of degradation occurs such that

$$c_{\text{ECM}}(p, t+1) = \max(0, c_{\text{ECM}}(p, t) - k_{\text{deg},m_p,t}) \tag{6.8}$$

where $k_{\text{deg},m_p,t}$ refers to the degradation rate of the cell, $m$, occupying pixel, $p$ and time, $t$. The maximum ensures that the concentration does not reduce below zero. Secondly, the effect of pushing away from and towards that pixel is resolved whereby

$$c_{\text{ECM}}(p, t+1) = \max\left(0, (1 - k_{\text{push},m_p,t})c_{\text{ECM}}(p, t+1) + \sum_{q=1}^{N_{\text{Neigh},p}} \frac{k_{\text{push},n_q,t}c_{\text{ECM}}(q, t)}{N_{\text{Neigh},q}}\right), \tag{6.9}$$

where $n_{q,t}$ refers to the pushing kinesis of the cell occupying neighbouring pixel, $q$ at time $t$.

#### 6.2 Code Documentation

The steppables responsible for updating the ECM concentration of the simulation environment differ between the organotypic and spheroid contexts due to the organotypic context requiring periodic boundary conditions and also possessing a defined region of zero concentration ECM (above the tumour).

##### 6.2.1 Organotypic ECM Concentration Update Steppable

The steppable responsible for updating the ECM concentration of the simulation environment in the organotypic context is OrganotypicECMConcentrationUpdate. The functions described below are given in the OrganotypicECMConcentrationUpdate.cpp file. The initiation function of the steppable identifies and links the simulation main frame to the steppable (Listing 6.2.1).

```
void OrganotypicECMConcentrationUpdate::init(Simulator *simulator, CC3DXMLElement
     *_xmlData) {

    potts = simulator->getPotts();
    sim=simulator;
    cellFieldG = (WatchableField3D<CellG *> *)potts->getCellFieldG();

    simulator->registerSteerableObject(this);
    update(_xmlData);
```

**Listing 6.2.1:** Organotypic ECM concentration update Steppable - init 1

Multiple plugins pre-defined in the Compucell3D package are utilised in this steppable, such as the plugin to track the internal pixels, and boundary pixels of all cells. These plugins are introduced to the steppable in the initiation function (Listing 6.2.2).

```
bool pluginAlreadyRegisteredFlag;
Plugin *plugin=Simulator::pluginManager.get("VolumeTracker",&
     pluginAlreadyRegisteredFlag);
cerr<<"GOT HERE BEFORE CALLING INIT"<<endl;
if(!pluginAlreadyRegisteredFlag)
plugin->init(simulator);

Plugin *pluginCOM=Simulator::pluginManager.get("CenterOfMass",&
     pluginAlreadyRegisteredFlag);
cerr<<"GOT HERE BEFORE CALLING INIT"<<endl;
if(!pluginAlreadyRegisteredFlag)
pluginCOM->init(simulator);

pixelTrackerPlugin=(PixelTrackerPlugin*)Simulator::pluginManager.get("
     PixelTracker",&pluginAlreadyRegisteredFlag);
if(!pluginAlreadyRegisteredFlag)
pixelTrackerPlugin->init(simulator);

boundaryPixelTrackerPlugin=(BoundaryPixelTrackerPlugin*)Simulator::pluginManager.
     get("BoundaryPixelTracker",&pluginAlreadyRegisteredFlag);
if(!pluginAlreadyRegisteredFlag)
boundaryPixelTrackerPlugin->init(simulator);

pixelTrackerAccessorPtr=pixelTrackerPlugin->getPixelTrackerAccessorPtr();
boundaryPixelTrackerAccessorPtr=boundaryPixelTrackerPlugin->
     getBoundaryPixelTrackerAccessorPtr();
}
```

**Listing 6.2.2:** Organotypic ECM concentration update Steppable - init 2

The update function, Listing 6.2.3, reads the input in the simulation setup xml file and allocates them to the steppable. This includes VCAF and SCC cell types, the dimensions of the simulated system and parameters concerning SCC and VCAF kinesis, degradation, pushing values of each cell-type. For VCAFs, input kinesis is in terms of the maximum and minimum values described by Equation (4.4). Since SCCs have fixed kinesis, the remodelling kinesis value is input such that the remodelling ability of an SCC is equivalent to a VCAF's travelling at the same speed (see Equations (6.6) and (6.7)). Each cell type's degradation and pushing rates are included as is the energy penalty scaling factor for cells moving into denser matrix (see Equations (6.1) to (6.3)). The desired ECM concentration in pixels occupied by cells is input in the xml file but pixels occupied by cells initialised above the tissue are always set to an ECM density of zero. This tissue boundary is a user input. An input filename to read in intial conditions for matrix density as well as output filename and print period for printing ECM density are included. The input ECM file can provide homogeneous or heterogeneous ECM concentrations as input. Heterogeneous inputs could include ECM concentrations that were lower or higher over specific defined regions.

```

void OrganotypicECMConcentrationUpdate::update(CC3DXMLElement *_xmlData, bool _fullInitFlag){

    if(_xmlData->findElement("VCAFtype"))
VCAFtype=_xmlData->getFirstElement("VCAFtype")->getDouble();
    if(_xmlData->findElement("SCCtype")){SCCtype=_xmlData->getFirstElement("SCCtype")->getDouble();}
    else{SCCtype=1;cerr<<"SCCtype not specified, defaul t value set to: "<<SCCtype<<endl;}
    if(_xmlData->findElement("dimx")){dimx=_xmlData->getFirstElement("dimx")->getDouble();}
    if(_xmlData->findElement("dimy")){dimy=_xmlData->getFirstElement("dimy")->getDouble();}
    if(_xmlData->findElement("dimz")){dimz=_xmlData->getFirstElement("dimz")->getDouble();}
    if(_xmlData->findElement("SCCLambdaTaxis_VCAFequiv")){lambdaSCC_VCAFequiv=_xmlData->getFirstElement("SCCLambdaTaxis_VCAFequiv")->getDouble();}
    if(_xmlData->findElement("VCAFLambdaTaxis_min")){lambdamin=_xmlData->getFirstElement("VCAFLambdaTaxis_min")->getDouble();}
    if(_xmlData->findElement("VCAFLambdaTaxis_max")){lambdamax=_xmlData->getFirstElement("VCAFLambdaTaxis_max")->getDouble();}
    if(_xmlData->findElement("ECMDegradationRate_max_vcaf")){ecmDegradationRate_max_vcaf=_xmlData->getFirstElement("ECMDegradationRate_max_vcaf")->getDouble();}
    if(_xmlData->findElement("ECMPushingRate_max_vcaf")){ecmPushingRate_max_vcaf=_xmlData->getFirstElement("ECMPushingRate_max_vcaf")->getDouble();}
    if(_xmlData->findElement("ECMDegradationRate_max_scc")){ecmDegradationRate_max_scc=_xmlData->getFirstElement("ECMDegradationRate_max_scc")->getDouble();}
    if(_xmlData->findElement("ECMPushingRate_max_scc")){ecmPushingRate_max_scc=_xmlData->getFirstElement("ECMPushingRate_max_scc")->getDouble();}
    if(_xmlData->findElement("VCAFLambdaECMpenetration")){VCAFLambdaECMpenetration=_xmlData->getFirstElement("VCAFLambdaECMpenetration")->getDouble();}
    if(_xmlData->findElement("SCCLambdaECMpenetration")){SCCLambdaECMpenetration=_xmlData->getFirstElement("SCCLambdaECMpenetration")->getDouble();}
    if(_xmlData->findElement("ECMOutputFileName")){ECMOutputFileName=_xmlData->getFirstElement("ECMOutputFileName")->getText();}
    if(_xmlData->findElement("ECMOutputSavePeriod")){ECMOutputSavePeriod=_xmlData->getFirstElement("ECMOutputSavePeriod")->getDouble();}
    if(_xmlData->findElement("ECMInputFileName")){ECMInputFileName=_xmlData->getFirstElement("ECMInputFileName")->getText();}
    if(_xmlData->findElement("ECMSpaceBoundary")){SpaceBoundary=_xmlData->getFirstElement("ECMSpaceBoundary")->getDouble();}
    else{SpaceBoundary=10000;cerr<<"ECMSpaceBoundary not specified, default value set to: "<<SpaceBoundary<<endl;}
    if(_xmlData->findElement("CellECMConcentration")){CellECMConc=_xmlData->getFirstElement("CellECMConcentration")->getDouble();}
    else{CellECMConc=0.0;cerr<<"CellECMConcentration not specified, default value set to: "<<CellECMConc<<endl;}
}
```

**Listing 6.2.3:** Organotypic ECM concentration update Steppable - update

The 'start' function, Listing 6.2.4, is called at the beginning of the simulation, allocating the variables of the steppable object. The ECM concentration field is defined, together with a time hash map, recording the last time step each grid point has been modified. The time hash map is implemented to increase speed of execution, and provides a faster way of checking if the grid point has already been recorded for update. This avoids revisiting the same ECM location more than once. The ECM concentration field and time map are allocated dynamically but are first input for concentration equal to one and timestep zero respectively. The ECM input file is then opened and ECM concentration updated according to this input file to allow for heterogeneous ECM concentration.

```
void OrganotypicECMConcentrationUpdate::start(){
    const int x=dimx;
    const int y=dimy;
    const int z=dimz;
    ECMconcfield = new float**[x];
    for(int i = 0; i < x; i++){
        ECMconcfield[i]= new float*[y];
        for (int j = 0; j< y; j++){
            ECMconcfield[i][j] = new float[z];
        }
    }
    pixelupdatestep = new int**[x];
    for(int i = 0; i < x; i++){
        pixelupdatestep[i]= new int*[y];
        for (int j = 0; j< y; j++){
            pixelupdatestep[i][j] = new int[z];
        }
    }
    for (int i = 0; i< x; i++){
    for (int j = 0; j< y; j++){
    for (int k = 0; k< z; k++){
        ECMconcfield[i][j][k]=1.0;
        pixelupdatestep[i][j][k]=0;
    }}}

    const char *inputname;
    inputname=ECMInputFileName.c_str();
    ECMInputFile.open(inputname,ifstream::in);
    if(!ECMInputFile.is_open()){cerr<<"could not open "<<ECMInputFileName<<" !!"
        <<endl;}
    int xcoord, ycoord, zcoord;
    float concentration;
    while( !ECMInputFile.eof() ) {
        ECMInputFile>>xcoord;
        ECMInputFile>>ycoord;
        ECMInputFile>>zcoord;
        ECMInputFile>>concentration;
        ECMconcfield[xcoord][ycoord][zcoord]=concentration;
        pixelupdatestep[xcoord][ycoord][zcoord]=0;
    }
    ECMInputFile.close();
}
```

**Listing 6.2.4:** Organotypic ECM concentration update Steppable - start

The 'step' function, Listing 6.2.5, carries out the necessary modifications, executed at the end of grid copy attempts. At the initial step, the cells have their ECM penetration energy parameters, degradation and pushing rates allocated according to their types, and are given the address of ECM concentration field, for ease of access during the copy attempt energy calculations. ECM density in pixel locations where cells are initialised is set to a user determined value, typically zero. However, the ECM density of cell pixels initialised above the ECM boundary are always set to zero, to reflect the fact that above the tissue is empty space. The flag 'steppablesinitialised' is also switched to the value of 1 in each cell. The energy calculations are carried out before the call of steppables (see Figure 2). This flag will be used in the ECMPenetrationPlugin (Section 6.2.3), to ensure the energy calculations are not carried out before the initial call (step 0) of this steppable. Otherwise ECMPenetrationPlugin would run prior to the ECM field address being allocated to each cell, causing a segmentation fault and crash of the simulation.

```
void OrganotypicECMConcentrationUpdate::step(const unsigned int currentStep){
const int ECMSavePeriod=ECMOutputSavePeriod;

    CellInventory *cellInventoryPtr=& potts->getCellInventory();
    CellInventory::cellInventoryIterator cInvItr;
    CellG *cell;
    if(currentStep==0){
        for(cInvItr=cellInventoryPtr->cellInventoryBegin() ; cInvItr !=
            cellInventoryPtr->cellInventoryEnd() ;++cInvItr ){
            cell=cellInventoryPtr->getCell(cInvItr);
            cell->ECMfieldbyCell=ECMconcfield;
            if((int)cell->type==VCAFtype){
                cell->lambdaECMPenetration=VCAFLambdaECMpenetration;
                cell->kdeg=ecmDegradationRate_max_vcaf;
                cell->wpush=ecmPushingRate_max_vcaf;
            }
            else if((int)cell->type==SCCtype){
                cell->lambdaECMPenetration=SCCLambdaECMpenetration;
                cell->kdeg=ecmDegradationRate_max_scc;
                cell->wpush=ecmPushingRate_max_scc;
            }
            set<PixelTrackerData> cellPixels=pixelTrackerAccessorPtr->get(cell->
                extraAttribPtr)->pixelSet;
            for(set<PixelTrackerData>::iterator sitr=cellPixels.begin() ; sitr !=
                 cellPixels.end() ;++sitr){
                int xx=sitr->pixel.x;int yy=sitr->pixel.y;int zz=sitr->pixel.z;
                if(zz<=SpaceBoundary){
                    ECMconcfield[xx][yy][zz]=0.0;
                }
                else{
                    ECMconcfield[xx][yy][zz]=CellECMConc;
                }
            }
            cell->steppablesinitialised=1;
        }
    }
```

**Listing 6.2.5:** Organotypic ECM concentration update Steppable - step 1

In the following steps, Listing 6.2.6, cells that have just divided are assigned their ECM concentration field and ECM penetration parameter. The cell's remodelling ability is updated depending on cell type and the current ECM degradation and pushing rates calculated from their kinesis stimulation (Equations (6.4) and (6.5)).

```
else{
    for(cInvItr=cellInventoryPtr->cellInventoryBegin() ; cInvItr !=
        cellInventoryPtr->cellInventoryEnd() ;++cInvItr ){
    cell=cellInventoryPtr->getCell(cInvItr);
```

```
    if (cell->just_divided == true){
        cell->ECMfieldbyCell=ECMconcfield;
        cell->lambdaECMPenetration=SCCLambdaECMpenetration;
        cell->steppablesinitialised=1;
        cell->just_divided=false;
    }
    vector <int> xcoord,ycoord,zcoord;
    vector <float> kdeglist,wpushlist;
    for(cInvItr=cellInventoryPtr->cellInventoryBegin() ; cInvItr !=
        cellInventoryPtr->cellInventoryEnd() ;++cInvItr ){
        cell=cellInventoryPtr->getCell(cInvItr);
        float weightlambda;
        float currkdeg;
        float currwpush;
        if((int)cell->type==VCAFtype){
            weightlambda=(cell->lambdataxis-lambdamin)/(lambdamax-lambdamin);
        }
        else if((int)cell->type==SCCtype){
            weightlambda=(lambdaSCC_VCAFequiv-lambdamin)/(lambdamax-lambdamin);
        }
        currkdeg=cell->kdeg*weightlambda;
        currwpush=cell->wpush*weightlambda;
```

**Listing 6.2.6:** Organotypic ECM concentration update Steppable - step 2

The cells are scanned to extract the grid points subjected to ECM update. All pixels within one pixel neighbourhood of cell boundaries, and pixels occupied by cells, are recorded into a coordinates list, together with the current degradation and pushing rates of the cell occupying the pixel. Once a pixel is recorded in this list, the time hash map is updated to indicate this pixel was recorded for current time, and before adding any pixel to the list, the time hash map is checked to avoid duplication (Listing 6.2.7).

```

    set<BoundaryPixelTrackerData> cellBoundaryPixels=
        boundaryPixelTrackerAccessorPtr->get(cell->extraAttribPtr)->pixelSet;
    for(set<BoundaryPixelTrackerData>::iterator sitr=cellBoundaryPixels.begin() ;
         sitr != cellBoundaryPixels.end() ;++sitr){
    int xx=sitr->pixel.x;int yy=sitr->pixel.y;int zz=sitr->pixel.z;
    for(int i=-1;i<2;i++){
        for(int j=-1;j<2;j++){
            for(int k=-1;k<2;k++){
                int currz=zz+k;
                if(currz>0 && currz<dimz){
                    int currx=xx+i;int curry=yy+j;
                    if(currx>=dimx){currx=currx-dimx;}else if(currx<0){currx=
                        currx+dimx;}
                    if(curry>=dimy){curry=curry-dimy;}else if(curry<0){curry=
                        curry+dimy;}
                    if(pixelupdatestep[currx][curry][currz]!=currentStep){
                        xcoord.push_back(currx);ycoord.push_back(curry);zcoord.
                            push_back(currz);
                        kdeglist.push_back(currkdeg);wpushlist.push_back(
                            currwpush);
                        pixelupdatestep[currx][curry][currz]=currentStep;
                    }
                }
            }
        }
    }
}
    set<PixelTrackerData> cellPixels=pixelTrackerAccessorPtr->get(cell->
        extraAttribPtr)->pixelSet;
    for(set<PixelTrackerData>::iterator sitr=cellPixels.begin() ; sitr !=
        cellPixels.end() ;++sitr){
        int xx=sitr->pixel.x;int yy=sitr->pixel.y;int zz=sitr->pixel.z;
        if(pixelupdatestep[xx][yy][zz]!=currentStep){
```

```
                xcoord.push_back(xx);ycoord.push_back(yy);zcoord.push_back(zz);
                kdeglist.push_back(currkdeg);wpushlist.push_back(currwpush);
                pixelupdatestep[xx][yy][zz]=currentStep;
            }
        }
}
```

**Listing 6.2.7:** Organotypic ECM concentration update Steppable - step 3

Once all pixels to be updated are recorded, the list is randomly shuffled to avoid introducing any bias to the evolution of the simulation from the order of the cell list (Listing 6.2.8).

```

    const int N=xcoord.size();
    int randomarray[N];
    for (int i=0;i<N;i++){randomarray[i]=i;}
    for (int i=0; i<(N-1); i++){
        int r=i + (rand() % (N-i));
        int temp = randomarray[i];
        randomarray[i] = randomarray[r];
        randomarray[r] = temp;
    }
```

**Listing 6.2.8:** Organotypic ECM concentration update Steppable - step 4

The list is iterated over in this random order, starting with degradation. In the case that the rate of degradation reduces ECM concentration below zero, the concentration is reset to zero. Then the concentration to be pushed to the neighbouring cells is calculated, and distributed to all neighbours within a one pixel neighbourhood (refer to Equations (6.4) and (6.5)). The concentration increase in this pixel due to ECM being pushed into it by the neighbours (second term in Equation (6.5)) will be accounted for in the update of the relevant neighbour (Listing 6.2.9).

```

    vector<int>::iterator intItr;
    vector<float>::iterator floatItr;
    for (int m=0;m<N;m++){
        intItr=xcoord.begin();intItr=intItr+randomarray[m];int xx=(*intItr);
        intItr=ycoord.begin();
        intItr=intItr+randomarray[m];
        int yy=(*intItr);
        intItr=zcoord.begin();
        intItr=intItr+randomarray[m];
        int zz=(*intItr);
        floatItr=kdeglist.begin();
        floatItr=floatItr+randomarray[m];
        float currkdeg=(*floatItr);
        floatItr=wpushlist.begin();
        floatItr=floatItr+randomarray[m];
        float currwpush=(*floatItr);
        ECMconcfield[xx][yy][zz]=ECMconcfield[xx][yy][zz]-currkdeg;
        if(ECMconcfield[xx][yy][zz]<0.0){ECMconcfield[xx][yy][zz]=0.0;}
        float pushedvalue=ECMconcfield[xx][yy][zz]*currwpush/26.0;
    for(int i=-1;i<2;i++){
    for(int j=-1;j<2;j++){
    for(int k=-1;k<2;k++){
        if (i!=0 || j!=0 || k!=0){
            int currz=zz+k;
            if(currz>0 && currz<dimz){
                int currx=xx+i;int curry=yy+j;
                if(currx>=dimx){currx=currx-dimx;}else if(currx<0){currx=currx+
                    dimx;}
```

```
                if(curry>=dimy){curry=curry-dimy;} else if(curry<0){curry=curry+dimy;}
                ECMconcfield[currx][curry][currz]=ECMconcfield[currx][curry][currz]+pushedvalue;
            }
        }
    }}}
    ECMconcfield[xx][yy][zz]=ECMconcfield[xx][yy][zz]*(1-currwpush);
    }
}
```

**Listing 6.2.9:** Organotypic ECM concentration update Steppable - step 5

An output text file, Listing 6.2.10, recording the ECM density at each location is generated, if the current time step corresponds to the user input save period.

```

if(currentStep%ECMSavePeriod==0){
    int timestepwrite=currentStep;
    const char *outputname;
    string paddedtsstring, ecmoutputname_var, filetype;
    stringstream paddedts;
    paddedts <<setw(5) << setfill('0') <<timestepwrite;
    paddedtsstring=paddedts.str();
    ecmoutputname_var=ECMOutputFileName+paddedtsstring+filetype;
    outputname=ecmoutputname_var.c_str();
    ECMOutputFile.open(outputname,ofstream::trunc);
    if (!ECMOutputFile.is_open()){cerr<<"could not open "<<ecmoutputname_var<<"!! "<<endl;}
        ECMOutput();
        ECMOutputFile.close();
    }
}
```

**Listing 6.2.10:** Organotypic ECM concentration update Steppable - step 6

The function generating ECM density output for a specified timepoint is given by Listing 6.2.11. Only values not equal to one are output to the text file.

```

void OrganotypicECMConcentrationUpdate::ECMOutput(){
    const int x=dimx;
    const int y=dimy;
    const int z=dimz;
    for (int i = 0; i< x; i++){
        for (int j = 0; j< y; j++){
            for (int k = 0; k< z; k++){
                if (ECMconcfield[i][j][k]!=1.0){
                    if(ECMconcfield[i][j][k]!=0.0){
                        ECMOutputFile<<setw(5) <<i<<" "<<setw(5) <<j<<" "<<setw(5) <<k<<"    " <<ECMconcfield[i][j][k]<<endl;
                    }
                    else if(ECMconcfield[i][j][k]==0.0){
                        ECMOutputFile<<setw(5) <<i<<" "<<setw(5) <<j<<" "<<setw(5) <<k<<"    " <<showpoint<<setprecision(2)<<0.0<<endl;
                    }
                }
            }
        }
    }
}
```

**Listing 6.2.11:** Organotypic ECM concentration update Steppable - step 7

##### 6.2.2 Spheroid ECM Concentration Update Steppable

The spheroid steppable, SpheroidECMConcentrationUpdate is a stripped down version of the organotypic case. The spheroid context does not include periodic boundary conditions and has no empty space above the tissue to account for. Due to the similarities, we present only the instances of code that are markedly different from the organotypic case. The functions described below are given in the SpheroidECMConcentrationUpdate.cpp file.

In the step function, at MCS, 0, the only significant difference to the organotypic steppable (Listing 6.2.5) is that there is no region space devoid of ECM in the spheroid assay. Only the ECM density in pixels occupied by cells within the ECM needs to be considered.This can be seen in line 12 of Listing 6.2.12.

```
1
void SpheroidECMConcentrationUpdate::step(const unsigned int currentStep){
3       const int ECMSavePeriod=ECMOutputSavePeriod;
        CellInventory *cellInventoryPtr=& potts->getCellInventory();
5       CellInventory::cellInventoryIterator cInvItr;
        CellG *cell;
7       if(currentStep==0){
            for(cInvItr=cellInventoryPtr->cellInventoryBegin() ; cInvItr !=
                cellInventoryPtr->cellInventoryEnd() ;++cInvItr ){
9               set<PixelTrackerData> cellPixels=pixelTrackerAccessorPtr->get(cell->
                    extraAttribPtr)->pixelSet;
                for(set<PixelTrackerData>::iterator sitr=cellPixels.begin() ; sitr !=
                     cellPixels.end() ;++sitr){
11                  int xx=sitr->pixel.x;int yy=sitr->pixel.y;int zz=sitr->pixel.z;
                    ECMconcfield[xx][yy][zz]=CellECMConc;
13              }
            }
15      }
```

**Listing 6.2.12:** Spheroid ECM concentration update Steppable - step 1

The only other differences concern the lack of periodic boundary conditions in the $x$ and $y$ dimensions in the spheroid context. Compare Listing 6.2.13 to Listing 6.2.7 (lines 11 and 12 removed) and Listing 6.2.14 to Listing 6.2.9 (lines 20 and 21 removed).

```

    else{
        for(cInvItr=cellInventoryPtr->cellInventoryBegin() ; cInvItr !=
            cellInventoryPtr->cellInventoryEnd() ;++cInvItr ){
            set<BoundaryPixelTrackerData> cellBoundaryPixels=
                boundaryPixelTrackerAccessorPtr->get(cell->extraAttribPtr)->
                pixelSet;
            for(set<BoundaryPixelTrackerData>::iterator sitr=cellBoundaryPixels.
                begin() ; sitr != cellBoundaryPixels.end() ;++sitr){
                int xx=sitr->pixel.x;int yy=sitr->pixel.y;int zz=sitr->pixel.z;
                for(int i=-1;i<2;i++){
                    for(int j=-1;j<2;j++){
                        for(int k=-1;k<2;k++){
                            int currz=zz+k;
                            int currx=xx+i;
                            int curry=yy+j;
                            if(currz>0 && currz<dimz && curry>0 && curry<dimy &&
                                currx>0 && currx<dimx){
                                if(pixelupdatestep[currx][curry][currz]!=
                                    currentStep){
                                    xcoord.push_back(currx);ycoord.push_back(
                                        curry);zcoord.push_back(currz);
                                    kdeglist.push_back(currkdeg);wpushlist.
                                        push_back(currwpush);
```

```
                                        pixelupdatestep[currx][curry][currz]=
                                            currentStep;
                                    }
                                }
                            }
                        }
                    }
                }
            }
```

Listing 6.2.13: Spheroid ECM concentration update Steppable - step 3

```
            vector<int>::iterator intItr;
            vector<float>::iterator floatItr;
            for (int m=0;m<N;m++){
                intItr=xcoord.begin();intItr=intItr+randomarray[m];int xx=(*intItr);
                intItr=ycoord.begin();intItr=intItr+randomarray[m];int yy=(*intItr);
                intItr=zcoord.begin();intItr=intItr+randomarray[m];int zz=(*intItr);
                floatItr=kdeglist.begin();floatItr=floatItr+randomarray[m];float
                    currkdeg=(*floatItr);
                floatItr=wpushlist.begin();floatItr=floatItr+randomarray[m];float
                    currwpush=(*floatItr);
                if(ECMconcfield[xx][yy][zz]<0){
                    cout<<"current concentration read: "<<ECMconcfield[xx][yy][zz]<<
                        endl;
                }
                ECMconcfield[xx][yy][zz]=ECMconcfield[xx][yy][zz]-currkdeg;if(
                    ECMconcfield[xx][yy][zz]<0.0){ECMconcfield[xx][yy][zz]=0.0;}
                float pushedvalue=ECMconcfield[xx][yy][zz]*currwpush/26.0;
                for(int i=-1;i<2;i++){
                    for(int j=-1;j<2;j++){
                        for(int k=-1;k<2;k++){
                            if (i!=0 || j!=0 || k!=0){
                                int currz=zz+k;
                                int currx=xx+i;
                                int curry=yy+j;
                                if(currz>0 && currz<dimz && curry>0 && curry<dimy &&
                                    currx>0 && currx<dimx ){
                                    ECMconcfield[currx][curry][currz]=ECMconcfield[
                                        currx][curry][currz]+pushedvalue;
                                }
                            }
                        }
                    }
                }
                ECMconcfield[xx][yy][zz]=ECMconcfield[xx][yy][zz]*(1-currwpush);
            }
        }
}
```

Listing 6.2.14: Spheroid ECM concentration update Steppable - step 5

##### 6.2.3 ECM Penetration Plugin

The plugin responsible for calculating the energy contribution of ECM penetration to the current pixel copy attempt is 'ECMPenetrationPlugin'. The functions described below are given in the 'ECMPenetrationPlugin.cpp' file.

```

void ECMPenetrationPlugin::init(Simulator *simulator, CC3DXMLElement *_xmlData)
{
    potts = simulator->getPotts();
    bool pluginAlreadyRegisteredFlag;
    Plugin *plugin=Simulator::pluginManager.get("VolumeTracker",&
        pluginAlreadyRegisteredFlag);
```

```
    if(!pluginAlreadyRegisteredFlag)
        plugin->init(simulator);
    potts->registerEnergyFunctionWithName(this, toString());
    xmlData=_xmlData;

    simulator->registerSteerableObject(this);
    update(_xmlData);
}
```

**Listing 6.2.15:** ECM penetration Plugin - init

Listing 6.2.16 reads the simulation inputs although most of the input parameters are set in the ECM remodelling steppable (Section 6.2.1 and 6.2.2). In rare circumstances of high pressure buildup, single cells can be forced entirely into high density ECM despite the high penalty for doing so. Once surrounded by the high density ECM on all sides, the penalty for moving within it it any given direction disappears. Therefore, incorporated within this plugin is a mechanism that makes the energy penalty of moving into high density matrix so large that it becomes impossible, overriding the standard matrix penalty. Only matrix below a certain density can all pixels within a cell move into. If the the matrix density is above this density then only boundary pixels of that cell can move into the pixel and only if the degradation and pushing rates are above a defined level. The input for this plugin is related to defining this mechanism. ECMDensityThreshold sets the ECM density beyond which cells suffer an extreme energy penalty for attempting to penetrate. Pixels on the boundary of that cell receive the standard energy penalty and are still able to penetrate a pixel above this density if either the cell's degradation or pushing rates fall above MinimumDegThreshold or MinimumPushThreshold respectively. The determination of the boundary thickness and record of each cell's boundary pixels is made by the plugins OrganotypicCellBoundaryECMPenetration and SpheroidCellBoundaryECMPenetration (see Sections 6.2.4 and 6.2.5 ).

```

void ECMPenetrationPlugin::update(CC3DXMLElement *_xmlData, bool _fullInitFlag)
{
if(_xmlData->findElement("ECMDensityThreshold")){ecmDensityThreshold=_xmlData->
    getFirstElement("ECMDensityThreshold")->getDouble();}
else{ecmDensityThreshold=0.9;cerr<<"ECMDensityThreshold not specified, default
    value set to: "<<ecmDensityThreshold<<endl;}
if(_xmlData->findElement("MinimumDegThreshold")){minDeg=_xmlData->getFirstElement
    ("MinimumDegThreshold")->getDouble();}
else{minDeg=0.0001;cerr<<"MinimumDegThreshold not specified, default value set to
    : "<<minDeg<<endl;}
if(_xmlData->findElement("MinimumPushThreshold")){minPush=_xmlData->
    getFirstElement("MinimumPushThreshold")->getDouble();}
else{minPush=0.001;cerr<<"MinimumPushThreshold not specified, default value set
    to: "<<minPush<<endl;}
}
```

**Listing 6.2.16:** ECM penetration Plugin - update

The 'changeEnergy' function defines how the energy of the copy attempt will be calculated. The function, Listing 6.2.17, initially checks if newCell and oldCell are the same, and returns energy change as zero if they are, avoiding any unnecessary calculation steps.

```
double ECMPenetrationPlugin::changeEnergy(const Point3D &pt,const CellG *newCell,
    const CellG *oldCell)
{
    double energy = 0.0;
    if (oldCell == newCell) return 0;
```

**Listing 6.2.17:** ECM penetration Plugin - change energy 1

If newCell has just divided then it will be missing a pointer to ECMfieldbyCell. Initially, Listing 6.2.18, oldCell is checked to see if its pointer is correctly set up. If it is, then newCell is assigned oldCell's pointer.

```

    CellG *cellforECMfield;
    CellInventory *cellInventoryPtr=& potts->getCellInventory();
    bool foundPointerToECMField = false;
    if (newCell && newCell->steppablesinitialised==1){
        if (newCell->just_divided){
            if(oldCell && oldCell->steppablesinitialised==1 && oldCell->
                just_divided==false){
                CellG *cell2;
                for(CellInventory::cellInventoryIterator cInvItr2=
                    cellInventoryPtr->cellInventoryBegin() ; cInvItr2 !=
                    cellInventoryPtr->cellInventoryEnd() ;++cInvItr2 ){
                    cell2=cellInventoryPtr->getCell(cInvItr2);
                    if (cell2->id == newCell->id){
                        cell2->ECMfieldbyCell=oldCell->ECMfieldbyCell;
                        cell2->just_divided=false;
                        foundPointerToECMField = true;
                        break;
                    }
                }
            }
```

**Listing 6.2.18:** ECM penetration Plugin - change energy 2

If oldCell's pointer is not correctly set up, then we search for a cell where it is. We then assign newCell this cell's pointer, Listing 6.2.19.

```

    else{
        for(CellInventory::cellInventoryIterator cInvItr2=cellInventoryPtr->
            cellInventoryBegin() ; cInvItr2 !=cellInventoryPtr->cellInventoryEnd
            () ;++cInvItr2 ){
            cellforECMfield=cellInventoryPtr->getCell(cInvItr2);
            if (cellforECMfield->just_divided==false){
                CellG *cell2;
                for(CellInventory::cellInventoryIterator cInvItr2=
                    cellInventoryPtr->cellInventoryBegin() ; cInvItr2 !=
                    cellInventoryPtr->cellInventoryEnd() ;++cInvItr2 ){
                    cell2=cellInventoryPtr->getCell(cInvItr2);
                    if (cell2->id == newCell->id){
                        cell2->ECMfieldbyCell=cellforECMfield->ECMfieldbyCell;
                        cell2->just_divided=false;
                        foundPointerToECMField = true;
                        break;
                    }
                }
                break;
            }
        }
    }
    if (!foundPointerToECMField){cerr<<"COULD NOT FIND A POINTER TO ECM FIELD!!!
        WILL CRASH!!!"<<endl;}
}
```

**Listing 6.2.19:** ECM penetration Plugin - change energy 3

The energy penalty of ECM Penetration from newCell is then calculated, Listing 6.2.20. Pixels on the cell boundary suffer only the standard penalty for ECM penetration if the cell has a degradation and/or pushing level greater than the user input threshold levels regardless of ECM density. The standard energy contribution of ECM is given by the ECM concentration at that pixel multiplied by the ECM Penetration value, $\lambda_{\text{ECM,NewCell}}$, (Equation (6.1)), for that cell (Listing 6.2.20,line 12). That is, boundary pixels for cells of a given remodelling ability are always able to penetrate into ECM of any density. Allowing such boundary pixels to penetrate into dense matrix represents the effect of small protrusions into dense matrix as part of the overall remodelling effect. Failing to allow the cell's boundary to penetrate into dense matrix would render the cell unable to remodel dense matrix at all. The list of pixels on the cell boundary is allocated in the OrganotypicCellBoundaryECMPenetration and SpheroidCellBoundaryECMPenetration plugins (Sections 6.2.4 and 6.2.5). Boundary pixels of cells that have remodelling ability below the threshold and pixels from the interior of any cell, regardless of remodelling ability penetrate the ECM with the standard energy penalty only if the ECM is below a threshold value, given in the xml file. Pixel copy attempts into ECM above this threshold for all pixels of cells with low remodelling capability or interior pixels of cells with high remodelling ability comes at the cost of an energy penalty of $10^{10}$. This makes it almost impossible for the interior of cells to penetrate dense matrix but allows the exterior to potentially remodel dense matrix.

```
 2      bool inNewCellBoundaryList=false;
        int flipx=pt.x;int flipy=pt.y;int flipz=pt.z;
 4      int nx = newCell->canremodel_x.size();
        for (int ii=0;ii<nx;ii++){
 6          if(newCell->canremodel_x[ii]==flipx && newCell->canremodel_y[ii]==flipy
                && newCell->canremodel_z[ii]==flipz){
                inNewCellBoundaryList=true;
 8              break;
            }
10      }

12      if (((newCell->kdeg>minDeg)||(newCell->wpush>minPush))&&(
            inNewCellBoundaryList==true)){
            energy=energy+newCell->ECMfieldbyCell[pt.x][pt.y][pt.z]*newCell->
                lambdaECMPenetration;
14      }
        else{
16          if (newCell->ECMfieldbyCell[pt.x][pt.y][pt.z]<ecmDensityThreshold){
                energy=energy+newCell->ECMfieldbyCell[pt.x][pt.y][pt.z]*newCell->
                    lambdaECMPenetration;
18          }
            else{
20              energy=energy+1e10;
            }
22      }
    }
```

**Listing 6.2.20:** ECM penetration Plugin - change energy 4

Exactly the same processes and energy calculations are also carried out for oldCell in Listing 6.2.21.

```
if (oldCell && oldCell->steppablesinitialised==1){
    if (oldCell->just_divided){
```

```
        if (newCell && newCell->steppablesinitialised==1 && newCell->just_divided==false){
            CellG *cell2;
            for(CellInventory::cellInventoryIterator cInvItr2=cellInventoryPtr->cellInventoryBegin() ; cInvItr2 !=cellInventoryPtr->cellInventoryEnd() ;++cInvItr2 ){
                cell2=cellInventoryPtr->getCell(cInvItr2);
                if (cell2->id == oldCell->id){
                    cell2->ECMfieldbyCell=newCell->ECMfieldbyCell;
                    cell2->just_divided=false;
                    foundPointerToECMField = true;
                    break;
                }
            }
        }
    else{
        for(CellInventory::cellInventoryIterator cInvItr2=cellInventoryPtr->cellInventoryBegin() ; cInvItr2 !=cellInventoryPtr->cellInventoryEnd() ;++cInvItr2 ){
            cellforECMfield=cellInventoryPtr->getCell(cInvItr2);
            if (cellforECMfield->just_divided==false){
                CellG *cell2;
                for(CellInventory::cellInventoryIterator cInvItr2=cellInventoryPtr->cellInventoryBegin() ; cInvItr2 !=cellInventoryPtr->cellInventoryEnd() ;++cInvItr2 ){
                    cell2=cellInventoryPtr->getCell(cInvItr2);
                    if (cell2->id == oldCell->id){
                        cell2->ECMfieldbyCell=cellforECMfield->ECMfieldbyCell;
                        cell2->just_divided=false;
                        foundPointerToECMField = true;
                        break;
                    }
                }
                break;
            }
        }
    }
    if (!foundPointerToECMField){cerr<<"COULD NOT FIND A POINTER TO ECM FIELD!!! WILL CRASH!!!"<<endl;}
    }


    bool inOldCellBoundaryList=false;
    int flipx=pt.x;int flipy=pt.y;int flipz=pt.z;
    int nx = oldCell->canremodel_x.size();
    for (int ii=0;ii<nx;ii++){
        if(oldCell->canremodel_x[ii]==flipx && oldCell->canremodel_y[ii]==flipy && oldCell->canremodel_z[ii]==flipz){
            inOldCellBoundaryList=true;
            break;
        }
    }

    if (((oldCell->kdeg>minDeg)||(oldCell->wpush>minPush))&&(inOldCellBoundaryList==true)){
        energy=energy-oldCell->ECMfieldbyCell[pt.x][pt.y][pt.z]*oldCell->lambdaECMPenetration;
    }
    else{
        if (oldCell->ECMfieldbyCell[pt.x][pt.y][pt.z]<ecmDensityThreshold){
            energy=energy-oldCell->ECMfieldbyCell[pt.x][pt.y][pt.z]*oldCell->lambdaECMPenetration;
        }
        else{
            energy=energy-1e10;
        }
    }
}
return energy;
}
```

**Listing 6.2.21:** ECM penetration Plugin - change energy 5

##### 6.2.4 Organotypic Cell Boundary ECM Penetration Plugin

These plugins simply record the pixels on each cell's boundary for use in the ECM Penetration Plugin 6.2.3. Only pixels within the boundary of each cell can potentially move into dense matrix.

The organotypic and spheroid contexts marginally differ as the organotypic context has periodic boundary conditions in $x$ and $y$ whilst the spheroid context does not. We begin with the organotypic context given by OrganotypicCellBoundaryECMPenetration and only briefly illustrate how the spheroid context differs. All functions below, starting with the initialisation (Listing 6.2.22) are given in the OrganotypicCellBoundaryECMPenetration.cpp file.

```
void OrganotypicCellBoundaryECMPenetration::init(Simulator *simulator,
    CC3DXMLElement *_xmlData) {

    potts = simulator->getPotts();
    sim=simulator;
    cellFieldG = (WatchableField3D<CellG *> *)potts->getCellFieldG();

    simulator->registerSteerableObject(this);
    update(_xmlData);
    bool pluginAlreadyRegisteredFlag;

    pixelTrackerPlugin=(PixelTrackerPlugin*)Simulator::pluginManager.get("
        PixelTracker",&pluginAlreadyRegisteredFlag);
    if(!pluginAlreadyRegisteredFlag)
        pixelTrackerPlugin->init(simulator);

    boundaryPixelTrackerPlugin=(BoundaryPixelTrackerPlugin*)Simulator::
        pluginManager.get("BoundaryPixelTracker",&pluginAlreadyRegisteredFlag);
    if(!pluginAlreadyRegisteredFlag)
    boundaryPixelTrackerPlugin->init(simulator);

    pixelTrackerAccessorPtr=pixelTrackerPlugin->getPixelTrackerAccessorPtr();
    boundaryPixelTrackerAccessorPtr=boundaryPixelTrackerPlugin->
        getBoundaryPixelTrackerAccessorPtr();

}
```

**Listing 6.2.22:** Organotypic Cell Boundary ECM Penetration Plugin - init

As input parameters, the steppable, Listing 6.2.23, takes cell type, simulation dimensions and NeighbourOrder. NeighbourOrder determines the width of the boundary to record. The larger it is, the more of the external cell is able to penetrate into dense matrix.

```
void OrganotypicCellBoundaryECMPenetration::update(CC3DXMLElement *_xmlData, bool
     _fullInitFlag){
    if(_xmlData->findElement("SCCtype")){SCCtype=_xmlData->getFirstElement("
        SCCtype")->getDouble();}
    else{SCCtype=1;cerr<<"SCCtype not specified, default value set to: "<<SCCtype
        <<endl;}
    if(_xmlData->findElement("VCAFtype")){VCAFtype=_xmlData->getFirstElement("
        VCAFtype")->getDouble();}
    else{VCAFtype=2;cerr<<"VCAFtype not specified, default value set to: "<<
        VCAFtype<<endl;}
```

```
    if(_xmlData->findElement("NeighbourOrder")){neighbourOrder=_xmlData->getFirstElement("NeighbourOrder")->getDouble();}
    else{neighbourOrder=1;cerr<<"NeighbourOrder not specified , default value set to: "<<neighbourOrder<<endl;}
    if(_xmlData->findElement("dimx")){dimx=_xmlData->getFirstElement("dimx")->getDouble();}
    if(_xmlData->findElement("dimy")){dimy=_xmlData->getFirstElement("dimy")->getDouble();}
    if(_xmlData->findElement("dimz")){dimz=_xmlData->getFirstElement("dimz")->getDouble();}
}
```

**Listing 6.2.23:** Organotypic Cell Boundary ECM Penetration Plugin - update

Four matrices are set up in Listing 6.2.24. cellIndexMatrix is set up to track the pixel location of all cells, timePointMatrix to quickly ascertain the locations of pixels in only the current timestep and cellBoundaryMatrix and timePointBoundaryMatrix which work similarly but solely for the cell boundary.

```
void OrganotypicCellBoundaryECMPenetration::start(){

    const int x=dimx;
    const int y=dimy;
    const int z=dimz;
    cellIndexMatrix = new int**[x];
    timePointMatrix  = new int**[x];
    cellBoundaryMatrix = new int**[x];
    timePointBoundaryMatrix = new int**[x];

    for(int i = 0; i < x; i++){
        cellIndexMatrix[i]= new int*[y];
        timePointMatrix[i]= new int*[y];
        cellBoundaryMatrix[i] = new int*[y];
        timePointBoundaryMatrix[i] = new int*[y];
        for (int j = 0; j< y; j++){
            cellIndexMatrix[i][j] = new int[z];
            timePointMatrix[i][j] = new int[z];
            cellBoundaryMatrix[i][j] = new int[z];
            timePointBoundaryMatrix[i][j] = new int[z];
        }
    }
    for (int i = 0; i< x; i++){
    for (int j = 0; j< y; j++){
    for (int k = 0; k< z; k++){
        cellIndexMatrix[i][j][k]=-1;
        timePointMatrix[i][j][k]=0;
        cellBoundaryMatrix[i][j][k] = -1;
        timePointBoundaryMatrix[i][j][k] = 0;
    }}}
}
```

**Listing 6.2.24:** Organotypic Cell Boundary ECM Penetration Plugin - start

In Listing 6.2.25 the cell ids are recorded for all cell pixel locations in cellIndexMatrix and the current timestep recorded in the corresponding pixel location in timePointMatrix.

```
void OrganotypicCellBoundaryECMPenetration::step(const unsigned int currentStep){
    CellInventory *cellInventoryPtr=& potts->getCellInventory();
    CellInventory::cellInventoryIterator cInvItr;
    CellG *cell;
    int neighbourOrder1=max(neighbourOrder-1,0);

    for(cInvItr=cellInventoryPtr->cellInventoryBegin() ; cInvItr !=
        cellInventoryPtr->cellInventoryEnd() ;++cInvItr ){
```

```
        cell=cellInventoryPtr->getCell(cInvItr);
        cell->canremodel_x.clear();
        cell->canremodel_y.clear();
        cell->canremodel_z.clear();
        set<PixelTrackerData> cellPixels=pixelTrackerAccessorPtr->get(cell->
            extraAttribPtr)->pixelSet;

        for(set<PixelTrackerData>::iterator sitr=cellPixels.begin() ; sitr !=
            cellPixels.end() ;++sitr){
            int xx=sitr->pixel.x;
            int yy=sitr->pixel.y;
            int zz=sitr->pixel.z;

            cellIndexMatrix[xx][yy][zz]=cell->id;
            timePointMatrix[xx][yy][zz]=currentStep;
        }
    }
```

**Listing 6.2.25:** Organotypic Cell Boundary ECM Penetration Plugin - step 1

Listing 6.2.26 goes through each boundary pixel for each cell. It is then determined if each of the neighbours of that pixel is part of the current cell or not. If it is then it is added to the set of canremodel vectors for that cell. The cellBoundaryMatrix and timePointBoundaryMatrix make sure that the same boundary pixel is not added multiple times to these vectors. Pixels in the canremodel vectors for each cell are the only ones able to move into matrix beyond the high density threshold.

```

    for(cInvItr=cellInventoryPtr->cellInventoryBegin() ; cInvItr !=
        cellInventoryPtr->cellInventoryEnd() ;++cInvItr ){
        cell=cellInventoryPtr->getCell(cInvItr);
        set<BoundaryPixelTrackerData>           cellBoundaryPixels=
            boundaryPixelTrackerAccessorPtr->get(cell->extraAttribPtr)->pixelSet;
        for(set<BoundaryPixelTrackerData>::iterator sitr=cellBoundaryPixels.begin
            () ; sitr != cellBoundaryPixels.end() ;++sitr){
            int xx=sitr->pixel.x;
            int yy=sitr->pixel.y;
            int zz=sitr->pixel.z;
            for(int i=-neighbourOrder1;i<neighbourOrder1+1;i++){
                for(int j=-neighbourOrder1;j<neighbourOrder1+1;j++){
                    for(int k=-neighbourOrder1;k<neighbourOrder1+1;k++){
                        int currz=zz+k;
                        if(currz>=0 && currz<dimz){
                                int currx=xx+i;int curry=yy+j;
                            if(currx>=dimx){
                                currx=currx-dimx;
                            }
                            else if(currx<0){
                                currx=currx+dimx;
                            }
                            if(curry>=dimy){
                                curry=curry-dimy;
                            }
                            else if(curry<0){
                                curry=curry+dimy;
                            }

                            if(timePointBoundaryMatrix[currx][curry][currz]!=
                                currentStep || cellBoundaryMatrix[currx][curry][
                                currz]!=cell->id ){
                                if(timePointMatrix[currx][curry][currz]==
                                    currentStep && cellIndexMatrix[currx][curry][
                                    currz]==cell->id ){
                                    cell->canremodel_x.push_back(currx);
                                    cell->canremodel_y.push_back(curry);
                                    cell->canremodel_z.push_back(currz);
```

```
33                                      cellBoundaryMatrix[currx][curry][currz]=cell->id;
                                        timePointBoundaryMatrix[currx][curry][currz]=currentStep;
35                                  }
                                }
37                          }
                        }
39                  }
                }
41          }
        }
43  }
```

**Listing 6.2.26:** Organotypic Cell Boundary ECM Penetration Plugin - step 2

##### 6.2.5 Spheroid Cell Boundary Steppable

The spheroid context is given by SpheroidCellBoundaryECMPenetration with all functions described in SpheroidCellBoundaryECMPenetration.cpp. The only difference in the spheroid context is the lack of periodic conditions present in organotypic Listing 6.2.26 (lines 15-26) in the spheroid Listing 6.2.27.

```
1
        for(cInvItr=cellInventoryPtr->cellInventoryBegin() ; cInvItr != cellInventoryPtr->cellInventoryEnd() ;++cInvItr ){
3       cell=cellInventoryPtr->getCell(cInvItr);
        set<BoundaryPixelTrackerData> cellBoundaryPixels=boundaryPixelTrackerAccessorPtr->get(cell->extraAttribPtr)->pixelSet;
5           for(set<BoundaryPixelTrackerData>::iterator sitr=cellBoundaryPixels.begin() ; sitr != cellBoundaryPixels.end() ;++sitr){
                int xx=sitr->pixel.x;
7               int yy=sitr->pixel.y;
                int zz=sitr->pixel.z;
9               for(int i=-neighbourOrder1;i<neighbourOrder1+1;i++){
                    for(int j=-neighbourOrder1;j<neighbourOrder1+1;j++){
11                      for(int k=-neighbourOrder1;k<neighbourOrder1+1;k++){
                            int currz=zz+k;
13                          int curry=yy+j;
                            int currx=xx+i;
15                          if(currz>0 && currz<dimz && curry>0 && curry<dimy && currx>0 && currx<dimx){
                                if(timePointBoundaryMatrix[currx][curry][currz]!=currentStep || cellBoundaryMatrix[currx][curry][currz]!=cell->id ){
17                                  if(timePointMatrix[currx][curry][currz]==currentStep && cellIndexMatrix[currx][curry][currz]==cell->id ){
                                        cell->canremodel_x.push_back(currx);
19                                      cell->canremodel_y.push_back(curry);
                                        cell->canremodel_z.push_back(currz);
21                                      cellBoundaryMatrix[currx][curry][currz]=cell->id;
                                        timePointBoundaryMatrix[currx][curry][currz]=currentStep;
23                                  }
                                }
25                          }
                        }
27                  }
                }
29          }
        }
31  }
```

**Listing 6.2.27:** Spheroid Cell Boundary ECM Penetration Plugin - step 2

### 7 ECM Adhesion

#### 7.1 Methodology

The contact energy between a cell and the ECM is dependent on the mean concentration of ECM surrounding the cell. Let $J_{\tau_{\sigma_L},ECM}$ be lowest contact energy (the maximum adhesion) of cell $\sigma$, of cell type $\tau$, to the ECM. Regardless of ECM density, the cell matrix adhesion cannot increase beyond this value, representing the possible saturation of cellular integrins. Let $J_{\tau_{\sigma_H},ECM}$ be the highest contact energy of cell $\sigma$ of type $\tau$ to ECM. That is, the lowest adhesion to the ECM, when ECM density is close or equal to zero. In the organotypic context, there is a region of space above the ECM construct. Let $J_{\tau,\emptyset}$ be the adhesion of cell $\tau$ to the medium if the cell is located in this region where there is no ECM. It would be expected that $J_{\tau_{\sigma_H},ECM} \leq J_{\tau,\emptyset}$ reflecting the fact that a region of the tissue structure with zero ECM may be assumed to still have things to stick to that would clearly not exist in the space above ECM, or, at worst, the cells have no more to adhere to than when it is in that space. This leads to the result that for cell $\sigma$, of cell type $\tau$ at MCS, $t$, the contact energy to ECM is given by

$$J_{\tau_\sigma,\mathrm{ECM}}(t) = \begin{cases} J_{\tau,\emptyset} & \text{in space above ECM,} \\ \frac{J_{\tau_{\sigma_H},ECM} J_{\tau_L,ECM}}{(J_{\tau_{\sigma_H},ECM} - J_{\tau_L,ECM})\bar{c}_{ECM}(\sigma,t)^r + J_{\tau_{\sigma_L},ECM}} & \text{in ECM below space, } ECM \leq 1, \\ J_{\tau_{\sigma_L},ECM} & \text{in ECM below space, } ECM > 1, \end{cases} \tag{7.1}$$

in the organotypic context and

$$J_{\tau_\sigma,\mathrm{ECM}}(t) = \begin{cases} \frac{J_{\tau_{\sigma_H},ECM} J_{\tau_L,ECM}}{(J_{\tau_{\sigma_H},ECM} - J_{\tau_L,ECM})\bar{c}_{ECM}(\sigma,t)^r + J_{\tau_{\sigma_L},ECM}} & ECM \leq 1, \\ J_{\tau_{\sigma_L},ECM} & ECM > 1, \end{cases} \tag{7.2}$$

in the spheroid context. Here,

$$\bar{c}_{ECM}(\sigma, t) = \frac{1}{|P_{\sigma_{\mathrm{neigh}}}|} \sum_{p \in P_{\sigma_{\mathrm{neigh}}}} c_{ECM}(p, t) \tag{7.3}$$

gives the mean ECM concentration of the set of all surrounding pixels, $P_{\sigma_{\mathrm{neigh}}}$, of cell $\sigma$ at time $t$ and $|P_{\sigma_{\mathrm{neigh}}}|$ gives the total number of pixel neighbours of the cell. The parameter $r > 0$ determines the inverse rate of increasing contact energy with decreasing ECM concentration. For small $r$, the contact energy rapidly approaches the minimum contact energy (maximum adhesion strength) for even very small matrix density. For large $r$, the contact energy tends to the minimum contact energy (maximum adhesion strength) much more more slowly over the matrix density range. Thus, a small $r$ implies that a cell has good adhesion to the matrix until there is almost no matrix left. A large $r$ implies that this drop in adhesion occurs more uniformly as matrix density decreases.

#### 7.2 Code Documentation

##### 7.2.1 Organotypic Adhesion ECM Concentration Steppable

The steppable for calculating ECM adhesion in the organotypic context is given by OrganotypicAdhesionECMConcentration. All functions below can be found in OrganotypicAdhesionECMConcentration.cpp. The initiation function, Listing 7.2.1, initialises relevant plugins, in this case all relating to pixel tracking in order to assess the ECM concentration in pixels surrounding each cell.

```
void OrganotypicAdhesionECMConcentration::init(Simulator *simulator,
    CC3DXMLElement *_xmlData) {

    potts = simulator->getPotts();
    sim=simulator;
    cellFieldG = (WatchableField3D<CellG *> *)potts->getCellFieldG();
    simulator->registerSteerableObject(this);
    update(_xmlData);

    bool pluginAlreadyRegisteredFlag;


    pixelTrackerPlugin=(PixelTrackerPlugin*)Simulator::pluginManager.get("
        PixelTracker",&pluginAlreadyRegisteredFlag);
    if(!pluginAlreadyRegisteredFlag)
        pixelTrackerPlugin->init(simulator);

    boundaryPixelTrackerPlugin=(BoundaryPixelTrackerPlugin*)Simulator::
        pluginManager.get("BoundaryPixelTracker",&pluginAlreadyRegisteredFlag);
    if(!pluginAlreadyRegisteredFlag)
        boundaryPixelTrackerPlugin->init(simulator);

    pixelTrackerAccessorPtr=pixelTrackerPlugin->getPixelTrackerAccessorPtr();
    boundaryPixelTrackerAccessorPtr=boundaryPixelTrackerPlugin->
        getBoundaryPixelTrackerAccessorPtr();
}
```

**Listing 7.2.1:** Organotypic Adhesion ECM Concentration Steppable - init

Simulation inputs in the xml file are read by the update function, Listing 7.2.2, and allocated to the steppable. SCC and VCAF cell types alongside dimensions of the simulated system are included. The contact energy for each cell type to the medium in the space above ECM and the boundary of this space in the z dimension are included. As is the minimum adhesion (maximum contact energy) for SCCs and VCAFs when ECM concentration is at its lowest as well as input for the parameter $r$, in Equation (7.1) that determines how quickly cells lose adhesion to ECM as the density decreases. Finally, the name of the input adhesion file is included, allowing each individual cell to possess its own maximum cell-matrix adhesion.

```
void OrganotypicAdhesionECMConcentration::update(CC3DXMLElement *_xmlData, bool
    _fullInitFlag){
    if(_xmlData->findElement("SCCtype")){SCCtype=_xmlData->getFirstElement("
        SCCtype")->getDouble();}
    else{SCCtype=1;cerr<<"SCCtype not specified, default value set to: "<<SCCtype
        <<endl;}
    if(_xmlData->findElement("VCAFtype")){VCAFtype=_xmlData->getFirstElement("
        VCAFtype")->getDouble();}
    else{VCAFtype=2;cerr<<"VCAFtype not specified, default value set to: "<<
        VCAFtype<<endl;}
```

```
    if(_xmlData->findElement("dimx")){dimx=_xmlData->getFirstElement("dimx")->
        getDouble();}
    if(_xmlData->findElement("dimy")){dimy=_xmlData->getFirstElement("dimy")->
        getDouble();}
    if(_xmlData->findElement("dimz")){dimz=_xmlData->getFirstElement("dimz")->
        getDouble();}
    if(_xmlData->findElement("SCCECMZeroConcAdhesion")){SCCECMMinAdh=_xmlData->
        getFirstElement("SCCECMZeroConcAdhesion")->getDouble();}
    else{SCCECMMinAdh=50;cerr<<"SCCECMZeroConcAdhesion not specified, default
        value set to: "<<SCCECMMinAdh<<endl;}
    if(_xmlData->findElement("VCAFECMZeroConcAdhesion")){VCAFECMMinAdh=_xmlData->
        getFirstElement("VCAFECMZeroConcAdhesion")->getDouble();}
    else{VCAFECMMinAdh=50;cerr<<"VCAFECMZeroConcAdhesion not specified, default
        value set to: "<<VCAFECMMinAdh<<endl;}
    if(_xmlData->findElement("ReciprocalPower")){RecipPower=_xmlData->
        getFirstElement("ReciprocalPower")->getDouble();}
    else{RecipPower=0.25;cerr<<"ReciprocalPower not specified, default value set
        to: "<<RecipPower<<endl;}
    if(_xmlData->findElement("SCCSpaceAdhesion")){SCCSpaceAdhesion=_xmlData->
        getFirstElement("SCCSpaceAdhesion")->getDouble();}
    else{SCCSpaceAdhesion=50;cerr<<"SCCSpaceAdhesion not specified, default value
         set to: "<<SCCSpaceAdhesion<<endl;}
    if(_xmlData->findElement("VCAFSpaceAdhesion")){VCAFSpaceAdhesion=_xmlData->
        getFirstElement("VCAFSpaceAdhesion")->getDouble();}
    else{VCAFSpaceAdhesion=50;cerr<<"VCAFSpaceAdhesion not specified, default
        value set to: "<<VCAFSpaceAdhesion<<endl;}
    if(_xmlData->findElement("ECMSpaceBoundary")){SpaceBoundary=_xmlData->
        getFirstElement("ECMSpaceBoundary")->getDouble();}
    else{SpaceBoundary=10000;cerr<<"ECMSpaceBoundary not specified, default value
         set to: "<<SpaceBoundary<<endl;}
    if(_xmlData->findElement("VariableCellAdhesionInputFileName")){
        CellAdhesionInputFileName=_xmlData->getFirstElement("
        VariableCellAdhesionInputFileName")->getText();}
}
```

**Listing 7.2.2:** Organotypic Adhesion ECM Concentration Steppable - update

In the start function (Listing 7.2.3), time hash maps record the last timepoint that each gridpoint has been modified and by which cell, both for each entire cell (timePointMatrix and cellIndexMatrix respectively) and gridpoints on the cell boundary (timePointBoundaryMatrix and cellBoundaryMatrix respectively). This provides a faster way of checking (see Listing 6.2.4 of Subsection 6.2.1).

```

void OrganotypicAdhesionECMConcentration::start(){
    const int x=dimx;
    const int y=dimy;
    const int z=dimz;
    cellIndexMatrix = new int**[x];
    timePointMatrix  = new int**[x];
    cellBoundaryMatrix = new int**[x];
    timePointBoundaryMatrix = new int**[x];

    for(int i = 0; i < x; i++){
        cellIndexMatrix[i]= new int*[y];
        timePointMatrix[i]= new int*[y];
        cellBoundaryMatrix[i] = new int*[y];
        timePointBoundaryMatrix[i] = new int*[y];
        for (int j = 0; j< y; j++){
            cellIndexMatrix[i][j] = new int[z];
            timePointMatrix[i][j] = new int[z];
            cellBoundaryMatrix[i][j] = new int[z];
            timePointBoundaryMatrix[i][j] = new int[z];
        }
    }
    for (int i = 0; i< x; i++){
    for (int j = 0; j< y; j++){
    for (int k = 0; k< z; k++){
```

```
            cellIndexMatrix[i][j][k]=-1;
            timePointMatrix[i][j][k]=0;
            cellBoundaryMatrix[i][j][k] = -1;
            timePointBoundaryMatrix[i][j][k] = 0;
        }}}
}
```

**Listing 7.2.3:** Organotypic Adhesion ECM Concentration Steppable - start

For MCS zero, in the step function, the ECM adhesion input file is read to allocate the maximum ECM adhesion for each single cell (Listing 7.2.4). This adhesive allocation can be heterogeneous.

```

void OrganotypicAdhesionECMConcentration::step(const unsigned int currentStep){
    CellInventory *cellInventoryPtr=& potts->getCellInventory();
    CellInventory::cellInventoryIterator cInvItr;
    CellG *cell;

    if(currentStep==0){
        const char *inputname;
        inputname=CellAdhesionInputFileName.c_str();
        CellAdhesionInputFile.open(inputname,ifstream::in);
        if(!CellAdhesionInputFile.is_open()){cerr<<"could not open "<<
            CellAdhesionInputFile<<" !!"<<endl;}
        int counter=-1;
        int tempinput1;
        string tempinput2;
        float tempinput3;
        CellAdhesionInputFile>>tempinput2;
        CellAdhesionInputFile>>tempinput2;
        CellAdhesionInputFile>>tempinput2;
        CellAdhesionInputFile>>tempinput2;
        CellAdhesionInputFile>>tempinput2;
        CellAdhesionInputFile>>tempinput2;
        CellAdhesionInputFile>>tempinput2;
        CellAdhesionInputFile>>tempinput2;
        CellAdhesionInputFile>>tempinput2;
        CellAdhesionInputFile>>tempinput2;

        for(cInvItr=cellInventoryPtr->cellInventoryBegin() ; cInvItr !=
            cellInventoryPtr->cellInventoryEnd() ;++cInvItr ){
            cell=cellInventoryPtr->getCell(cInvItr);
            if (cell->type == SCCtype){
                cell->MaximumVCAFECMAdhesion=0.0;
                cell->CurrentVCAFECMAdhesion=0.0;
                cell->ZeroVCAFECMAdhesion=0.0;
                CellAdhesionInputFile>>tempinput1;
                CellAdhesionInputFile>>tempinput2;
                CellAdhesionInputFile>>tempinput3;
                CellAdhesionInputFile>>tempinput3;
                CellAdhesionInputFile>>cell->MaximumSCCECMAdhesion;
                CellAdhesionInputFile>>tempinput3;
                CellAdhesionInputFile>>tempinput3;
                CellAdhesionInputFile>>tempinput3;
                CellAdhesionInputFile>>tempinput3;
                CellAdhesionInputFile>>tempinput3;
            }
            else if(cell->type == VCAFtype){
                cell->MaximumSCCECMAdhesion=0.0;
                cell->CurrentSCCECMAdhesion=0.0;
                cell->ZeroSCCECMAdhesion=0.0;
                CellAdhesionInputFile>>tempinput1;
                CellAdhesionInputFile>>tempinput2;
                CellAdhesionInputFile>>tempinput3;
                CellAdhesionInputFile>>tempinput3;
                CellAdhesionInputFile>>tempinput3;
                CellAdhesionInputFile>>tempinput3;
                CellAdhesionInputFile>>tempinput3;
```

```
                CellAdhesionInputFile>>cell->MaximumVCAFECMAdhesion;
                CellAdhesionInputFile>>tempinput3;
                CellAdhesionInputFile>>tempinput3;
            }
        }
    CellAdhesionInputFile.close();
    }
```

**Listing 7.2.4:** Organotypic Adhesion ECM Concentration Steppable - step 1

For subsequent timepoints, the location of all cellular pixels are recorded alongside the current timestep at those locations ( Listing 7.2.5).

```

    for(cInvItr=cellInventoryPtr->cellInventoryBegin() ; cInvItr !=
        cellInventoryPtr->cellInventoryEnd() ;++cInvItr ){
        cell=cellInventoryPtr->getCell(cInvItr);
        set<PixelTrackerData> cellPixels=pixelTrackerAccessorPtr->get(cell->
            extraAttribPtr)->pixelSet;
        for(set<PixelTrackerData>::iterator sitr=cellPixels.begin() ; sitr !=
            cellPixels.end() ;++sitr){
            int xx=sitr->pixel.x;
            int yy=sitr->pixel.y;
            int zz=sitr->pixel.z;
            cellIndexMatrix[xx][yy][zz]=cell->id;
            timePointMatrix[xx][yy][zz]=currentStep;
        }
    }
```

**Listing 7.2.5:** Organotypic Adhesion ECM Concentration Steppable - step 2

In Listing 7.2.6 the minimum depth of the cell is recorded. A depth of zero corresponds to the bottom of the ECM whilst the maximum depth corresponds to being in the region of space above the ECM. The recording of the minimum depth of a cell allows the allocation of a different adhesion for cells that are in this space as opposed to in ECM.

```

    for(cInvItr=cellInventoryPtr->cellInventoryBegin() ; cInvItr !=
        cellInventoryPtr->cellInventoryEnd() ;++cInvItr ){
        cell=cellInventoryPtr->getCell(cInvItr);
        set<BoundaryPixelTrackerData> cellBoundaryPixels=
            boundaryPixelTrackerAccessorPtr->get(cell->extraAttribPtr)->pixelSet;
        for(set<BoundaryPixelTrackerData>::iterator sitr=cellBoundaryPixels.begin
            () ; sitr != cellBoundaryPixels.end() ;++sitr){
            if(sitr==cellBoundaryPixels.begin()){
                int zz=sitr->pixel.z;
                cell->MinZExtent=zz;
            }
            else{
                if(sitr->pixel.z<cell->MinZExtent){
                    cell->MinZExtent=sitr->pixel.z;
                }
            }
        }
    }
```

**Listing 7.2.6:** Organotypic Adhesion ECM Concentration Steppable - step 3

In Listing 7.2.7 the neighbouring pixels of a cell's boundary pixels are checked. If the neighbouring boundary pixel is not part of that cell then the ECM concentration at that pixel is recorded in ECMNeighbourConcentrations. The boundary pixel time hash maps stop a neighbouring pixel from being double counted. The mean ECM concentration is then allocated to each cell.

```
    for(cInvItr=cellInventoryPtr->cellInventoryBegin() ; cInvItr !=
        cellInventoryPtr->cellInventoryEnd() ;++cInvItr ){
        cell=cellInventoryPtr->getCell(cInvItr);
        cell->ECMNeighbourConcentrations.clear();
        set<BoundaryPixelTrackerData> cellBoundaryPixels=
            boundaryPixelTrackerAccessorPtr->get(cell->extraAttribPtr)->pixelSet;
        for(set<BoundaryPixelTrackerData >::iterator sitr=cellBoundaryPixels.begin
            () ; sitr != cellBoundaryPixels.end() ;++sitr){
            int xx=sitr->pixel.x;
            int yy=sitr->pixel.y;
            int zz=sitr->pixel.z;
            for(int i=-1;i<2;i++){
                for(int j=-1;j<2;j++){
                    for(int k=-1;k<2;k++){
                        int currz=zz+k;
                        if(currz>=0 && currz<dimz){
                            int currx=xx+i;int curry=yy+j;
                            if(currx>=dimx){
                                currx=currx-dimx;
                            }
                        else if(currx<0){
                            currx=currx+dimx;
                        }
                        if(curry>=dimy){
                            curry=curry-dimy;
                        }
                        else if(curry<0){
                            curry=curry+dimy;
                        }
                        if(timePointBoundaryMatrix[currx][curry][currz]!=
                            currentStep || cellBoundaryMatrix[currx][curry][currz
                            ]!=cell->id ){
                            if(timePointMatrix[currx][curry][currz]!=currentStep
                                || cellIndexMatrix[currx][curry][currz]!=cell->id
                                ){
                                cell->ECMNeighbourConcentrations.push_back(cell->
                                    ECMfieldbyCell[currx][curry][currz]);
                                    cellBoundaryMatrix[currx][curry][currz]=cell
                                        ->id;
                                    timePointBoundaryMatrix[currx][curry][currz]=
                                        currentStep;
                                }
                            }
                        }
                    }
                }
            }
        }
        cell->MeanSurroundingECMConc=std::accumulate(cell->
            ECMNeighbourConcentrations.begin(),cell->ECMNeighbourConcentrations.
            end(),0.0);
        cell->MeanSurroundingECMConc=cell->MeanSurroundingECMConc/cell->
            ECMNeighbourConcentrations.size();
    }
```

**Listing 7.2.7:** Organotypic Adhesion ECM Concentration Steppable - step 4

In Listing 7.2.8 the cell's ECM adhesion is updated according the the ECM concentration surrounding the cell. If the cell is located in the space above where the ECM begins then the ECM adhesion can be set at a fixed value that could be very low. If the cell is located partially, or totally within the region of ECM then the cell's ECM adhesion at that timepoint is set according to Equation (7.1).

```
    for(cInvItr=cellInventoryPtr->cellInventoryBegin() ; cInvItr !=
        cellInventoryPtr->cellInventoryEnd() ;++cInvItr ){
```

```
        cell=cellInventoryPtr->getCell(cInvItr);
        if(cell->MinZExtent>SpaceBoundary){
            if (cell->type == SCCtype){
                cell->CurrentSCCECMAdhesion=SCCSpaceAdhesion;
            }
            else if (cell->type == VCAFtype){
                cell->CurrentVCAFECMAdhesion=VCAFSpaceAdhesion;
            }
        }
        else{
            if (cell->type == SCCtype){
                MinAdh=SCCECMMinAdh;
                MaxAdh=cell->MaximumSCCECMAdhesion;
            }
            else if (cell->type == VCAFtype){
                MinAdh=VCAFECMMinAdh;
                MaxAdh=cell->MaximumVCAFECMAdhesion;
            }
            quadraticB=MaxAdh/(MinAdh-MaxAdh);
            quadraticA=quadraticB*MinAdh;
            if (cell->type == SCCtype){
                cell->CurrentSCCECMAdhesion=quadraticA/(pow(cell->
                    MeanSurroundingECMConc,RecipPower)+quadraticB);
                cell->CurrentVCAFECMAdhesion=0.0;
                if (cell->CurrentSCCECMAdhesion<cell->MaximumSCCECMAdhesion){
                    cell->CurrentSCCECMAdhesion=cell->MaximumSCCECMAdhesion;
                }
            }
            else if (cell->type == VCAFtype){
                cell->CurrentSCCECMAdhesion=0.0;
                cell->CurrentVCAFECMAdhesion=quadraticA/(pow(cell->
                    MeanSurroundingECMConc,RecipPower)+quadraticB);
                if (cell->CurrentVCAFECMAdhesion<cell->MaximumVCAFECMAdhesion){
                    cell->CurrentVCAFECMAdhesion=cell->MaximumVCAFECMAdhesion;
                }
            }
        }
    }
}
```

**Listing 7.2.8:** Organotypic Adhesion ECM Concentration Steppable - step 5

##### 7.2.2 Spheroid Adhesion ECM Concentration Steppable

The spheroid steppable is simpler than the organotypic steppable and as such we present a condensed version highlighting only the differences. These differences arise because firstly there is no region of space above the ECM in the spheroid context and secondly, there are no periodic boundary conditions in $x$ and $y$ dimensions. As such, in the spheroid update steppable there are no references to this region of space. The steppable for the spheroid environment is given by SpheroidAdhesionECMConcentration with the functions below in the SpheroidAdhesionECMConcentration.cpp file. In Listing 7.2.2, parameters relating to SCCSpaceAdhesion, VCAFSpaceAdhesion and ECMSpaceBoundary are removed (lines 16-21). The lack of a region of space above the ECM makes the code (Listing 7.2.6) related to recording the depth of each cell in the $z$-dimension redundant and it is thus removed in the spheroid. Listing 7.2.7 in the organotypic context is then reduced to Listing 7.2.9 in the spehroid context, removing periodic boundary conditions.

```
    for(cInvItr=cellInventoryPtr->cellInventoryBegin() ; cInvItr !=
        cellInventoryPtr->cellInventoryEnd() ;++cInvItr ){
```

```
        cell=cellInventoryPtr->getCell(cInvItr);
        cell->ECMNeighbourConcentrations.clear();
        set<BoundaryPixelTrackerData> cellBoundaryPixels=
            boundaryPixelTrackerAccessorPtr->get(cell->extraAttribPtr)->pixelSet;
        for(set<BoundaryPixelTrackerData >::iterator sitr=cellBoundaryPixels.begin
            () ; sitr != cellBoundaryPixels.end() ;++sitr){
            int xx=sitr->pixel.x;
            int yy=sitr->pixel.y;
            int zz=sitr->pixel.z;
            for(int i=-1;i<2;i++){
                for(int j=-1;j<2;j++){
                    for(int k=-1;k<2;k++){
                        int currx=xx+i;int curry=yy+j; int currz=zz+k;
                        if(currx>=0 && currx<dimx && curry>=0 && curry<dimy &&
                            currz>=0 && currz<dimz){
                            if(timePointBoundaryMatrix[currx][curry][currz]!=
                                currentStep || cellBoundaryMatrix[currx][curry][
                                currz]!= cell->id ){
                                if(timePointMatrix[currx][curry][currz]!=
                                    currentStep || cellIndexMatrix[currx][curry][
                                    currz]!= cell->id ){
                                    cell->ECMNeighbourConcentrations.push_back(
                                        cell->ECMfieldbyCell[currx][curry][currz
                                        ]);
                                    cellBoundaryMatrix[currx][curry][currz]=cell
                                        ->id;
                                    timePointBoundaryMatrix[currx][curry][currz]=
                                        currentStep;
                                }
                            }
                        }
                    }
                }
            }
        }
        cell->MeanSurroundingECMConc=std::accumulate(cell->
            ECMNeighbourConcentrations.begin(), cell->ECMNeighbourConcentrations.
            end(),0.0);
        cell->MeanSurroundingECMConc=cell->MeanSurroundingECMConc/cell->
            ECMNeighbourConcentrations.size();
    }
```

**Listing 7.2.9:** Spheroid Adhesion ECM Concentration Steppable - step 4

Finally, Listing 7.2.8 is reduced to Listing 7.2.10 due to Equation (7.1) reducing to (7.1) in spheroid context.

```

    for(cInvItr=cellInventoryPtr->cellInventoryBegin() ; cInvItr !=
        cellInventoryPtr->cellInventoryEnd() ;++cInvItr ){
        cell=cellInventoryPtr->getCell(cInvItr);
        if (cell->type == SCCtype){
            MinAdh=SCCECMMinAdh;
            MaxAdh=cell->MaximumSCCECMAdhesion;
            }
        else if (cell->type == VCAFtype){
            MinAdh=VCAFECMMinAdh;
            MaxAdh=cell->MaximumVCAFECMAdhesion;
        }
        quadraticB=MaxAdh/(MinAdh-MaxAdh);
        quadraticA=quadraticB*MinAdh;
        if (cell->type == SCCtype){
            cell->CurrentSCCECMAdhesion=quadraticA/(pow(cell->
                MeanSurroundingECMConc, RecipPower)+quadraticB);
            cell->CurrentVCAFECMAdhesion=0.0;
            if (cell->CurrentSCCECMAdhesion<cell->MaximumSCCECMAdhesion){
                cell->CurrentSCCECMAdhesion=cell->MaximumSCCECMAdhesion;
            }
        }
```

```
            else if (cell->type == VCAFtype){
                cell->CurrentSCCECMAdhesion=0.0;
                cell->CurrentVCAFECMAdhesion=quadraticA/(pow(cell->
                    MeanSurroundingECMConc,RecipPower)+quadraticB);
                if (cell->CurrentVCAFECMAdhesion<cell->MaximumVCAFECMAdhesion){
                    cell->CurrentVCAFECMAdhesion=cell->MaximumVCAFECMAdhesion;
                }
            }
    }
}
```

**Listing 7.2.10:** Spheroid Adhesion ECM Concentration Steppable - step 5

### 8 cell.h Header File

Finally, these steppables and plugins require additional cellular attributes to be added to the cell.h header file. Listing 8.0.1 shows these additional properties. The number of additional cellular properties required by these steppables and plugins is large. It includes such attributes as the ECM degradation and pushing rates kdeg and wpush, the random time to cell division TimeToMitosisCheck, the centroid location of the spheroid, SpheroidCentroidX, SpheroidCentroidY and SpheroidCentroidZ, and the current directionality of each cell, taxisidir.

```

#ifndef CELL_H
#define CELL_H

#include <iostream>
#include <vector>
using namespace std;


#ifndef PyObject_HEAD
struct _object; //forward declare
typedef _object PyObject; //type redefinition
#endif


struct VCAFpositionnode;

class BasicClassGroup;

namespace CompuCell3D {

/**
* A Potts3D cell.
*/

    class CellG{
    public:
        typedef unsigned char CellType_t;
    CellG():
        volume(0),
        targetVolume(0.0),
        lambdaVolume(0.0),
        surface(0),
        targetSurface(0.0),
        lambdaSurface(0.0),
        clusterSurface(0.0),
        targetClusterSurface(0.0),
        lambdaClusterSurface(0.0),
        type(0),
        xCM(0),yCM(0),zCM(0),
```

```
        xCOM(0),yCOM(0),zCOM(0),
        xCOMPrev(0),yCOMPrev(0),zCOMPrev(0),
        iXX(0), iXY(0), iXZ(0), iYY(0), iYZ(0), iZZ(0),
        lX(0.0),
        lY(0.0),
        lZ(0.0),
        lambdaVecX(0.0),
        lambdaVecY(0.0),
        lambdaVecZ(0.0),
        flag(0),
        id(0),
        clusterId(0),
        fluctAmpl(-1.0),
        extraAttribPtr(0),
        pyAttrib(0),

        //Attributes added for steppables and plugins in the paper ****
        VCAFListroot(0),
        previousCM(),
        lambdataxis(0.0),
        taxisidir(),
        lambdaECMPenetration(0.0),
        steppablesinitialised(0),
        TimeToMitosisCheck(100000),
        InitialTimeToMitosis(100000),
        DaughterCellTimer(100000),
        OriginalTargetVolume(1.0)

    {
        //Attributes added for steppables and plugins in the paper ****
        previousCM[0]=xCOM; previousCM[1]=yCOM; previousCM[2]=zCOM;
        lambdataxis=0;
        taxisidir[0]=1.0; taxisidir[1]=0.0; taxisidir[2]=0.0;
        TimeToMitosisCheck=100000;
        InitialTimeToMitosis=100000;
        DaughterCellTimer=100000;
        DivideThisCellInPython=false;
        just_divided=false;
        just_divided_taxis_label=false;
        OriginalTargetSurface=0;
        InitialConditionCell=false;
        MaxCellVolume=0;
        mitosistimer=0;
    }
        unsigned long volume;
        float targetVolume;
        float lambdaVolume;
        double surface;
        float targetSurface;
        float angle;
        float lambdaSurface;
        double clusterSurface;
        float targetClusterSurface;
        float lambdaClusterSurface;
        unsigned char type;
        unsigned char subtype;
        double xCM,yCM,zCM; // numerator of center of mass expression (components
            )
        double xCOM,yCOM,zCOM; // numerator of center of mass expression (
            components)
        double xCOMPrev,yCOMPrev,zCOMPrev; // previous center of mass
        double iXX, iXY, iXZ, iYY, iYZ, iZZ; // tensor of inertia components
        float lX,lY,lZ; //orientation vector components - set by MomentsOfInertia
             Plugin - read only
        float ecc; // cell eccentricity
        float lambdaVecX,lambdaVecY,lambdaVecZ; // external potential lambda
            vector components
        unsigned char flag;
        float averageConcentration;
        long id;
```

```
        long clusterId;
        double fluctAmpl;
        BasicClassGroup *extraAttribPtr;
        PyObject *pyAttrib;

        //Attributes added for steppables and plugins in the paper ****

        VCAFpositionnode *VCAFListroot;
        double previousCM[3];
        float lambdataxis;
        double taxisidir[3];
        float*** ECMfieldbyCell;
        float lambdaECMPenetration;
        int steppablesinitialised;

        vector <int> canremodel_x;
        vector <int> canremodel_y;
        vector <int> canremodel_z;
        int  TimeToMitosisCheck;
        int  DaughterCellTimer;
        bool  just_divided;
        bool  just_divided_taxis_label;
        float  OriginalTargetVolume;
        float  OriginalTargetSurface;
        int MaxCellVolume;
        bool DivideThisCellInPython;
        bool InitialConditionCell;
        int  InitialTimeToMitosis;
        int mitosistimer;
        float kdeg;
        float wpush;
        float MaximumSCCECMAdhesion;
        float CurrentSCCECMAdhesion;
        float ZeroSCCECMAdhesion;
        float MaximumVCAFECMAdhesion;
        float CurrentVCAFECMAdhesion;
        float ZeroVCAFECMAdhesion;
        float MeanSurroundingECMConc;
        vector <float> ECMNeighbourConcentrations;
        int MinZExtent;
        int  SpheroidCentroidX;
        int  SpheroidCentroidY;
        int  SpheroidCentroidZ;
        long parentid;
    };


    class Cell {
    };

    class CellPtr{
    public:
        Cell * cellPtr;
    };
};
#endif
```

**Listing 8.0.1:** cell.h header file
